# Mechanistic insight into the dual phosphorylation of ERK1 and ERK2 by MEK1

**DOI:** 10.64898/2026.03.13.710243

**Authors:** Ying Sun, Chen Peng, Shengjie Liu, Fayang Zhou, Gaoxingyu Huang, Jianbo Wang, Qi Hu

**Affiliations:** Qilu Hospital of Shandong University, Jinan, Shandong, China; Fudan University, Shanghai, China; State Key Laboratory of Gene Expression, School of Life Sciences, Westlake University, Hangzhou, Zhejiang, China; School of Future Biomedicine, Westlake University, Hangzhou, Zhejiang, China; Institute of Biology, Westlake Institute for Advanced Study, Hangzhou, Zhejiang, China

**Author notes:** These authors contributed equally (Ying Sun, Chen Peng, Shengjie Liu).

## Abstract

Activation of ERK1 and ERK2 through MEK1 and MEK2-catalyzed phosphorylation of conserved tyrosine and threonine residues in the ERK1/2 activation loops is a key step in the RAS-RAF-MEK-ERK signaling cascade. However, the molecular mechanisms underlying MEK1/2-catalyzed ERK1/2 phosphorylation remain unclear. Here, we report the cryo-EM structures of phosphorylated MEK1 (pMEK1) bound to unphosphorylated ERK1/2 and monophosphorylated ERK1, together with a crosslinked pMEK1:ERK1 complex. These structures elucidate pMEK1-catalyzed dual phosphorylation of ERK1/2: the ERK1/2 activation loops interact with the MEK1 activation loop to enable tyrosine phosphorylation, and adopt a distinct conformation by interacting with the ERK1/2 HRD motifs to promote threonine phosphorylation in the ERK1/2 activation loops. Notably, we find that pMEK1 catalyzes ERK1/2 dephosphorylation and reveal that it is attributed to the reverse transfer of the phosphate from ERK1/2 to ADP. These findings delineate the catalytic cycle of MEK1/2-catalyzed ERK1/2 phosphorylation and provide insights for targeting this oncogenic pathway.

## INTRODUCTION

Mitogen-activated protein kinases (MAPKs) are serine/threonine kinases that regulate essential cellular processes, such as proliferation and differentiation^1^. Among MAPKs, extracellular signal-regulated kinases ERK1 (MAPK3) and ERK2 (MAPK1) are the most extensively studied due to their role as downstream effectors of the RAS-RAF-MEK-ERK cascade^2^. Upon activation by guanine nucleotide exchange factors (GEFs), RAS GTPases (HRAS, NRAS, KRAS4A and KRAS4B) bind to and activate RAF kinases (ARAF, BRAF, CRAF)^3^. RAF kinases phosphorylate and therefore activate the dual-specificity kinases MEK1 and MEK2, which in turn phosphorylate ERK1/2, leading to their activation and translocation to the nucleus to modulate gene transcription^2^.

MEK1/2 are phosphorylated by RAF kinases at two conserved serine residues (S218/S222 in MEK1; S222/S226 in MEK2)^4^. A large number of crystal structures of unphosphorylated MEK1/2 have been determined^5–9^, however, there is a lack of structures of phosphorylated MEK1/2. This gap limits our understanding of how MEK1/2 achieve their active conformations.

Activated MEK1/2 phosphorylate ERK1/2 at a tyrosine residue (Y204 in ERK1 or Y187 in ERK2) and a threonine residue (T202 in ERK1 or T185 in ERK2) in their activation loops^1^. It has been reported that the phosphorylation of ERK1/2 occurs in a defined order: the tyrosine residue is phosphorylated first, followed by the threonine residue^10^. The dual phosphorylation increases the catalytic activity of ERK1/2 by approximately four orders of magnitude^11^. Structural studies of ERK2 reveal that phosphorylation stabilizes the activation loop in an active conformation^12,13^.

Previous studies reveal that MAPKs interact with their substrates and regulators mainly through two sites: one is a shallow groove adjacent to the hinge sequence connecting the N-lobe and C-lobe, which recognizes short peptides named the “D domain” or “DEJL motif” (docking sites for ERK and JNK, LXL), and thus called D-site recruitment site (DRS)^14–16^; the other is a pocket near the activation loop, called F-site recruitment site (FRS), which interacts with peptides named the “DEF motif” (docking sites for ERK, FXF)^14–16^. Crystal structures of ERK2 in complex with DEJL or DEF motifs of MEK2 and several other ERK-interacting proteins^17–20^, and with both the DEJL and DEF motifs of PEA-15 (15 kDa phosphoprotein enriched in astrocytes)^21^, have been reported. These structures demonstrate how the two docking sites (DRS and FRS) in ERK1/2 interact with multiple proteins. However, a full picture of how ERK1/2 interacts with MEK1/2—their most important regulators—remains unclear, particularly how phosphorylation sites in the activation loops of ERK1/2 are positioned toward the catalytic sites of MEK1/2 to facilitate the phosphorylation reaction.

In this article, we report the cryo-electron microscopy (cryo-EM) structures of phosphorylated MEK1 (pMEK1) bound to unphosphorylated ERK1 and ERK2, and monophosphorylated ERK1, and a crosslinked pMEK1:ERK1 complex. These structures demonstrate that pMEK1 in these complexes undergoes dramatic conformational changes in its N-lobe and activation loop in comparison with the structures of unphosphorylated MEK1 reported previously^6^ to adopt an active conformation, explaining how MEK1 is activated. Structural analysis reveals how ERK1 is dual-phosphorylated at Y204 and T202: for Y204 phosphorylation, the ERK1 activation loop interacts with that of MEK1 to orient Y204 toward the MEK1 catalytic center; and for T202 phosphorylation, the ERK1 activation loop interacts with the HRD motif of ERK1 to adopt a distinct conformation. In addition, we identify that pMEK1 can catalyze ERK1 Y204 dephosphorylation, and reveal that the dephosphorylation is attributed to the reverse transfer of the phosphate from Y204 to ADP to produce ATP, followed by hydrolysis of the ATP.

## RESULTS

### Overall structure of dual-phosphorylated MEK1 in complex with unphosphorylated ERK1

We overexpressed MEK1 and ERK1 in *E. coli* and purified their unphosphorylated forms (named uMEK1 and uERK1, respectively). By incubating uMEK1 with BRAF carrying the oncogenic mutation V600E, we obtained MEK1 phosphorylated at S218 and S222 (named pMEK1) (Figure S1A). We verified that only pMEK1 but not uMEK1 can catalyze the phosphorylation of ERK1 (Figure S1B). In consistent with the kinase activity data, we found that only pMEK1 but not uMEK1 can form a stable complex with uERK1 (Figures S1C and S1D).

We assembled the pMEK1:uERK1 complex by mixing uERK1 with pMEK1 in the presence of a non-hydrolysable analog of ATP, adenosine 5′-(β,γ-imido)triphosphate (also called AMP-PNP). Using cryo-EM, we solved the structure of this complex. The map resolution is 3.36 Å (Table S1, Figures S2 and S3). The cryo-EM structure shows that MEK1 forms a heterodimer with ERK1 (Figure 1A). In the heterodimer, the N-lobe and C-lobe of ERK1 interact with the N-lobe and C-lobe of MEK1, respectively. The N-terminal peptide of MEK1 docks into the D-site recruitment site (DRS) in ERK1, while a helix of MEK1 C-lobe binds to the F-site recruitment site (FRS) in ERK1 (Figure 1A). The αA-helix between the N-terminal peptide and the N-lobe of MEK1 is largely invisible in the cryo-EM map, except for a short segment whose main chain is resolved (Figures 1A and S4).

**Figure 1.**
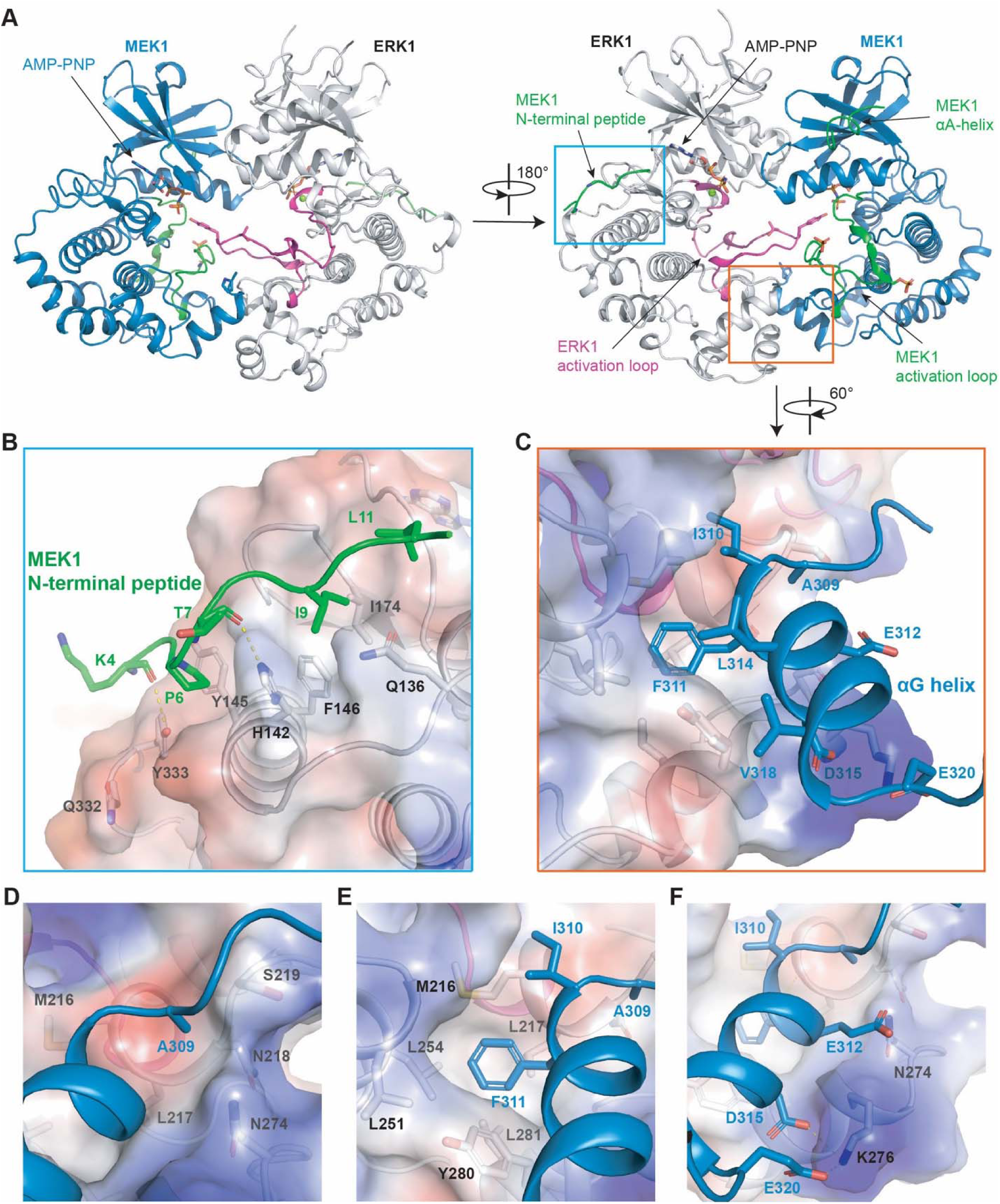
Cryo-EM structure of the pMEK1:uERK1 complex. (A) Overall structure of the pMEK1:uERK1 complex. Both pMEK1 and uERK1 in the complex bind to an AMP-PNP in their nucleotide binding pockets. (B) Interactions between the N-terminal peptide of MEK1 (colored green) and the D-site recruitment site (DRS) in ERK1. (C) Binding of the A309-E320 helix in the MEK1 C-lobe (colored blue) to the F-site recruitment site (FRS) in ERK1. (D–F) Close-up view of the binding of MEK1 A309 (D), I310 and F311 (E), and E312, D315 and E320 (F) to the FRS in ERK1. Hydrogen bonds are shown as yellow dash lines. All structural figures were made using PyMOL.

We also solved the cryo-EM structure of pMEK1 in complex with unphosphorylated ERK2 (uERK2) in the presence of AMP-PNP. The map resolution is 2.93 Å (Table S1, Figures S5 and S6). This structure closely resembles that of the pMEK1:uERK1 complex (Figures S7A and S7B), consistent with the high sequence identity (82.6%) between human ERK1 and ERK2.

### Interactions between MEK1 and two known docking sites in ERK1

The two docking sites, DRS and FRS, of ERK1/2 are known to be binding sites for many substrates and regulators of ERK1/2. In the pMEK1:uERK1 heterodimer, MEK1 also occupies the two docking sites.

As mentioned above, the N-terminal peptide (residues 4-12) of MEK1 binds to the DRS in ERK1 (Figure 1B). The interface area is 909 Å². This peptide interacts with ERK1 similarly to known DEJL motifs (Figure S8). Specifically, the side chains of P6 and I9 in this peptide dock into two hydrophobic pockets around H142 in ERK1. MEK1 L11 also interacts with ERK1 through hydrophobic interactions. Mutation of MEK1 I9 and L11 to alanine, as well as mutation of ERK1 H142 to glutamic acid, significantly inhibited MEK1-catalyzed ERK1 phosphorylation, especially the phosphorylation of ERK1 T202 (Figures S9A–S9D), supporting the importance of the MEK1 N-terminal peptide in recruiting ERK1 to MEK1 to facilitate phosphorylation. The interactions between MEK1 N-terminal peptide and ERK1 DRS are highly conserved in the pMEK1:uERK2 structure (Figure S7C). Mutation of MEK1 I9 and L11 to alanine also inhibited MEK1-catalyzed ERK2 phosphorylation (Figures S9E and S9F).

The A309-E320 helix in the MEK1 C-lobe, which is aligned to the αG helix in protein kinase A (PKA), binds to the FRS in ERK1 (Figure 1C) or ERK2 (Figure S7D). One face of this helix is enriched in hydrophobic residues: A309, I310, F311, L314, and V318. These residues mediate hydrophobic interactions with ERK1/2. Specifically, A309 inserts into a hydrophobic pocket formed by ERK1 residues M216-S219 and N274 (Figure 1D), while F311 occupies another hydrophobic pocket formed by ERK1 residues M216, L217, L251, L254, Y280, and L281 (Figure 1E) or the corresponding residues in ERK2 (Figures S7E and S7F). The MEK1 A309L mutation slightly decreased the kinase activity of MEK1 (Figures S9C–S9F). We also expressed the MEK1 F311A mutant, but found that it could not be phosphorylated by BRAF; therefore, its kinase activity was not evaluated. Notably, although MEK1 lacks the conserved FXFP motif found in ERK substrates^16^, I310 and F311 bind to positions in ERK1 analogous to the first and second phenylalanine residues in the FXFP motif, respectively (Figures S10A and S10B). The other face of MEK1 A309-E320 helix contains three negatively charged residues—E312, D315, and E320—that interact with the side chains of N274 and K276 in ERK1 through hydrogen bonding and electrostatic interactions (Figure 1F). The ERK1 K276D mutation almost completely blocked MEK1-catalyzed ERK1 phosphorylation (Figures S9G and S9H).

MEK1 binds to ERK1/2 in a manner similar to PEA-15, a regulator of ERK1/2 that has a death effector domain (DED) and a C-terminal tail^21^. Crystal structures reveal that the C-terminal tail of PEA-15 binds to the DRS in ERK2 (Figure S8F), while its DED binds to the FRS in ERK2 (Figures S10C and S10D)^21^. However, this DED binds to both the FRS and activation loop of ERK2 with two helices, which differs from how MEK1 binds the FRS in ERK1.

In addition to the above interactions, the MEK1 N-lobe is close to the ERK1 N-lobe in the complex. The interface, with an area of 892 Å², is mainly mediated by hydrophobic interactions. We attempted to perturb this interface by introducing the Y53A mutation, but found minimal effects on MEK1-catalyzed ERK1 phosphorylation (Figures S9A and S9B).

The overall structures of the pMEK1:uERK1 and pMEK1:uERK2 complexes are similar to that of an engineered MKK6 (dual specificity mitogen-activated protein kinase kinase 6) in complex with the T180V mutant of p38α (mitogen-activated protein kinase 14)^22^. However, the relative position between MEK1 and ERK1 is different from that between MKK6 and p38α: upon alignment of the C-lobe of ERK1 with that of p38α, the kinase domain of MEK1 exhibits a 9° rotation relative to that of MKK6 (Figure S11A). Another notable difference is in the conformations of the activation loops. The activation loops of ERK1 and MEK1 are well-resolved in the cryo-EM structure (Figure 2); in contrast, the activation loop of p38α adopts a different conformation while that of MKK6 is largely invisible (Figures S11B and S11C).

**Figure 2.**
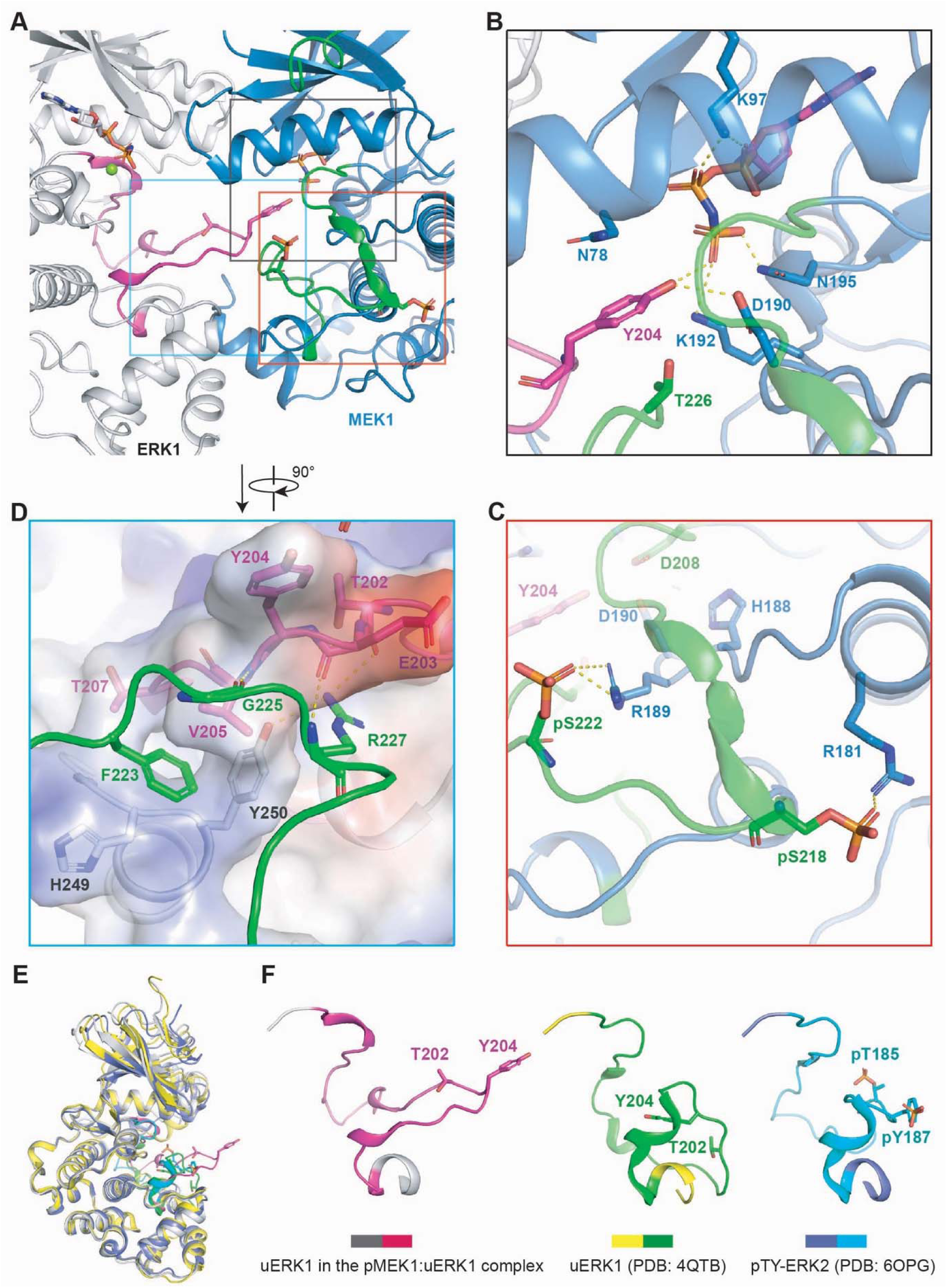
Interactions between the activation loop of uERK1 and pMEK1. (A) Overview of the interactions between uERK1 activation loop (colored magenta) and pMEK1 (colored blue/green). (B) Position of ERK1 Y204 (colored magenta) in the catalytic pocket of pMEK1. (C) Structure of the activation loop of pMEK1 in the pMEK1:uERK1 complex. MEK1 S218 and S222 are phosphorylated, and interact with MEK1 R181 and R189, respectively. (D) Interactions between pMEK1 activation loop (colored green) and ERK1. (E) Alignment of the cryo-EM structure of uERK1 (gray/magenta) in the pMEK1:uERK1 complex with the crystal structure of uERK1 alone (yellow/green, PDB: 4QTB) and that of dual-phosphorylated ERK2 (slate/cyan, PDB: 6OPG). (F) Structures of the activation loop of uERK1 in the pMEK1:uERK1 complex, of uERK1 alone, and of dual-phosphorylated ERK2.

### Binding of ERK1/2 activation loops to the catalytic pocket of MEK1

ERK1/2 activation requires phosphorylation of ERK1 T202 and Y204 or ERK2 T185 and Y187 in their activation loops by MEK1/2. Since the structure of pMEK1:uERK2 resembles that of pMEK1:uERK1, we will focus on the pMEK1:uERK1 structure to discuss the interactions between ERK1/2 activation loops and MEK1. In the pMEK1:uERK1 heterodimer, the ERK1 activation loop inserts into the catalytic pocket of MEK1 (Figure 2A). Specifically, the hydroxyl group of ERK1 Y204 points toward the γ-phosphate of AMP-PNP in the nucleotide-binding pocket of MEK1 (Figure 2B). The hydroxyl group of Y204 also forms a hydrogen bond with the side chain of D190 in the HRD motif in MEK1. The aromatic ring of Y204 sits between MEK1 N78 and T226. The γ-phosphate in MEK1 is stabilized by the side chains of K192 and N195. Such a structural arrangement prepares the transfer of the γ-phosphate to the hydroxyl oxygen of Y204. Mutations of MEK1 D190 or D208 (in the DFG motif) to alanine or mutation of K192 to methionine disrupted the kinase activity of MEK1, while the N78A mutation showed minimal effect (Figures S12A–S12F).

The activation loop of MEK1 plays an important role in positioning the activation loop of ERK1 into the catalytic pocket of MEK1. The MEK1 activation loop itself is stabilized by phosphorylated S218 and S222, which interact with the side chains of MEK1 R181 and R189, respectively, through charge-charge interactions and hydrogen bonding (Figure 2C). Three MEK1 residues downstream of S222, including F223, G225 and R227, directly interact with ERK1 activation loop (Figure 2D). Among them, F223 docks into a hydrophobic pocket formed by ERK1 V205, T207, H249, and Y250. MEK1 F223 also interacts with ERK1 Y250 through π-π stacking (Figure 2D). The MEK1 F223A mutation disrupted the kinase activity of MEK1 (Figures S9C–S9F). Additionally, the main chains of MEK1 G225 and R227 form two hydrogen bonds with the main chains of ERK1 V205 and E203, and the side chain of MEK1 R227 interacts with the main chain of ERK1 T202 and the side chain of ERK1 Y250 through hydrogen bonding (Figure 2D). The activation loop of ERK1 in the pMEK1:uERK1 complex is well ordered (Figures 2 and S11G). In contrast, in the previously reported structure of the chimeric MKK6:p38α(T180V) complex^22^, the p38α activation loop shows poor cryo-EM density, likely due to its flexibility (Figure S11H).

Structural alignment shows that the conformation of the ERK1 activation loop in the pMEK1-uERK heterodimer is distinct from that in reported ERK1/2 structures in either unphosphorylated or dual-phosphorylated states (Figures 2E and 2F), suggesting that binding to MEK1 induces conformational changes in the ERK1 activation loop to facilitate its phosphorylation.

### Conformation of MEK1 in its active state

In the pMEK1:uERK1 and pMEK1:uERK2 heterodimers, MEK1 adopts an active conformation. As no structure of active MEK1 had been reported previously, we aligned the structure of MEK1 in the pMEK1:uERK1 heterodimer with a crystal structure of inactive MEK1 to understand how MEK1 achieves its active conformation^6^. We identified three regions in MEK1 that undergo extensive conformational changes upon activation (Figure 3A).

**Figure 3.**
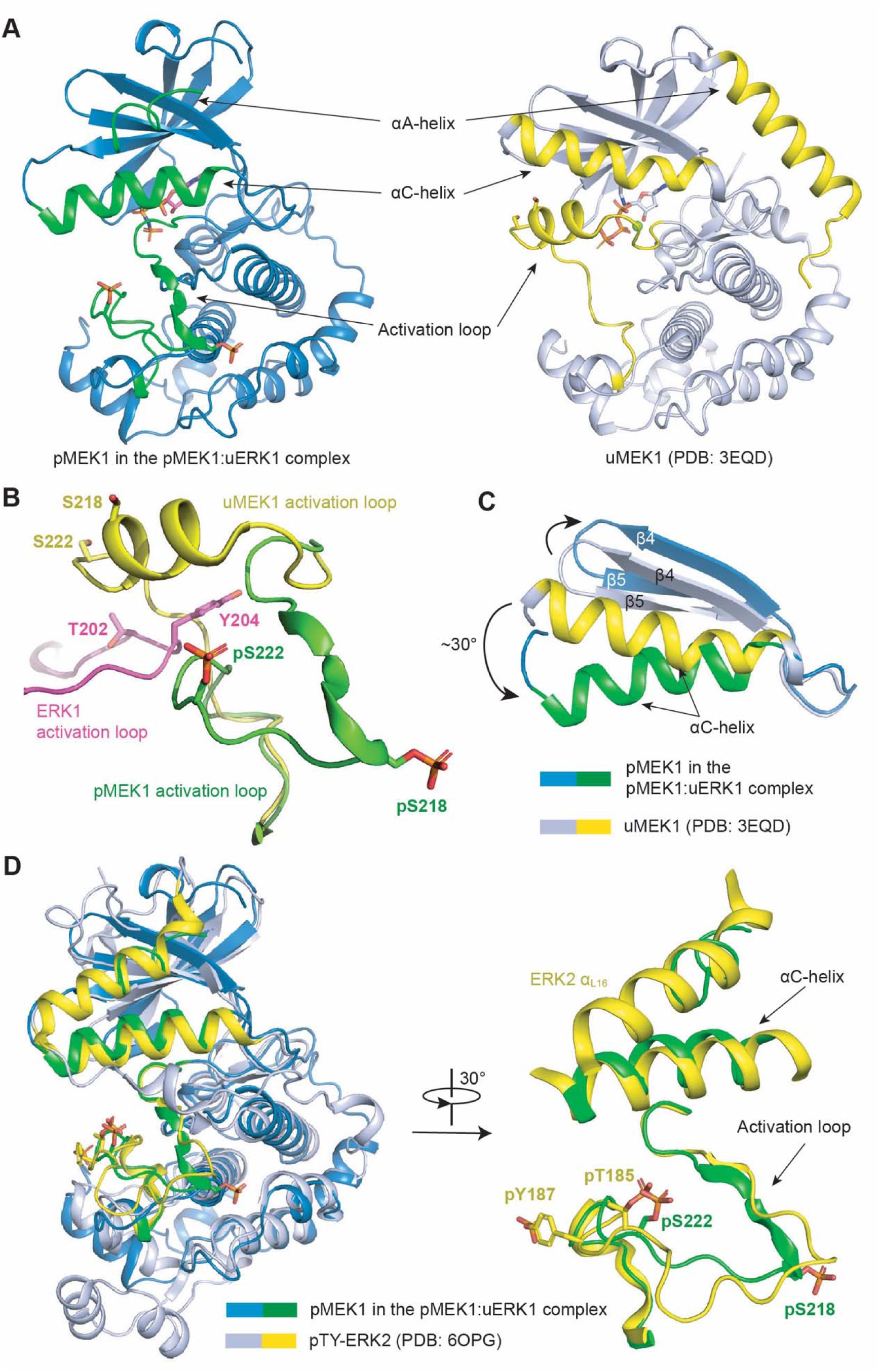
Structural reorganization of MEK1 underlying its activation. (A) Overall structure of pMEK1 in the pMEK1:uERK1 complex and that of uMEK1 alone (PDB: 3EQD). The C-lobes (residues 150-370) of MEK1 in the two structures were aligned. (B) Conformational differences of MEK1 activation loop between the inactive (yellow) and active (green) states. The activation loop of ERK1 bound to pMEK1 is colored magenta. (C) Conformational differences of MEK1 β4, β5 strands and αC-helix between the inactive (gray/yellow) and active (blue/green) states. (D) Alignment of the structure of pMEK1 in the pMEK1:uERK1 complex (blue/green) with that of dual-phosphorylated ERK2 (gray/yellow, PDB: 6OPG).

First, in inactive MEK1, the N-terminal half of its activation loop forms a helix that interacts with the αC-helix (Figure 3B). In contrast, in the pMEK1:uERK1 heterodimer, the MEK1 activation loop adopts an extended conformation and is anchored to the C-lobe via phosphorylated residues S218 and S222 (Figure 3B). Without this conformational change, the MEK1 activation loop would sterically clash with the ERK1 activation loop in the MEK:ERK complex (Figure 3B). Second, the αC-helix of MEK1 rotates ∼30° toward the C-lobe upon activation, while the β4 and β5 strands shift in the opposite direction, moving away from the αC-helix (Figure 3C). Third, the regulatory αA-helix extends toward the C-lobe in inactive MEK1 but becomes disordered in the active state (Figure 3A). In active MEK1, only the main chain of a short helical segment, which binds between the αC-helix and β4 strand, is resolved in the cryo-EM density map (Figures 3A and S4).

The conformational changes in the three regions are coupled with one another. Upon activation, movement of the MEK1 activation loop creates space for the αC-helix, which takes the position originally occupied by the N-terminal half of the activation loop in inactive MEK1. The rearrangements of the αC-helix and β4-β5 strands generate a shallow groove for the αA-helix to dock into. Binding of MEK1 N-terminal peptide to the DRS in ERK1 (Figure 1B) also contributes to the orienting direction of this αA-helix.

The conformation of active MEK1 resembles that of active ERK2 (Figure 3D). The overall topology and orientation of their activation loops are highly similar. Phosphorylated S222 in the MEK1 activation loop occupies a position structurally equivalent to phosphorylated T185 in ERK2 (T202 in ERK1). The position of the αC-helix in active MEK1 is almost identical to that in active ERK2. A segment of the αA-helix in active MEK1 mimics the C-terminal α-helix of ERK2 (α_L16_)^15^. Such structural similarity demonstrates the conservation of structural elements required for the kinase activity of MEK1/2 and ERK1/2.

### pMEK1 catalyzes the dephosphorylation of ERK1 pY204

The pMEK1:uERK1 structure reveals how ERK1 is recruited to MEK1 and how the first step of ERK1 phosphorylation (phosphorylation of Y204) occurs. To gain a comprehensive understanding of the MEK1-catalyzed phosphorylation process of ERK1, we sought to solve the structure of pMEK1 in complex with Y204 mono-phosphorylated ERK1 (pY-ERK1). We mixed pMEK1 with pY-ERK1 in an AMP-PNP-containing buffer and solved the cryo-EM structure (Table S1, Figures S2 and S13). Unexpectedly, the cryo-EM map suggests that ERK1 Y204 in the complex was unphosphorylated. That is, the pMEK1:pY-ERK1 complex actually yielded a cryo-EM structure of the pMEK1:uERK1 complex (Figures 4A and 4B).

**Figure 4.**
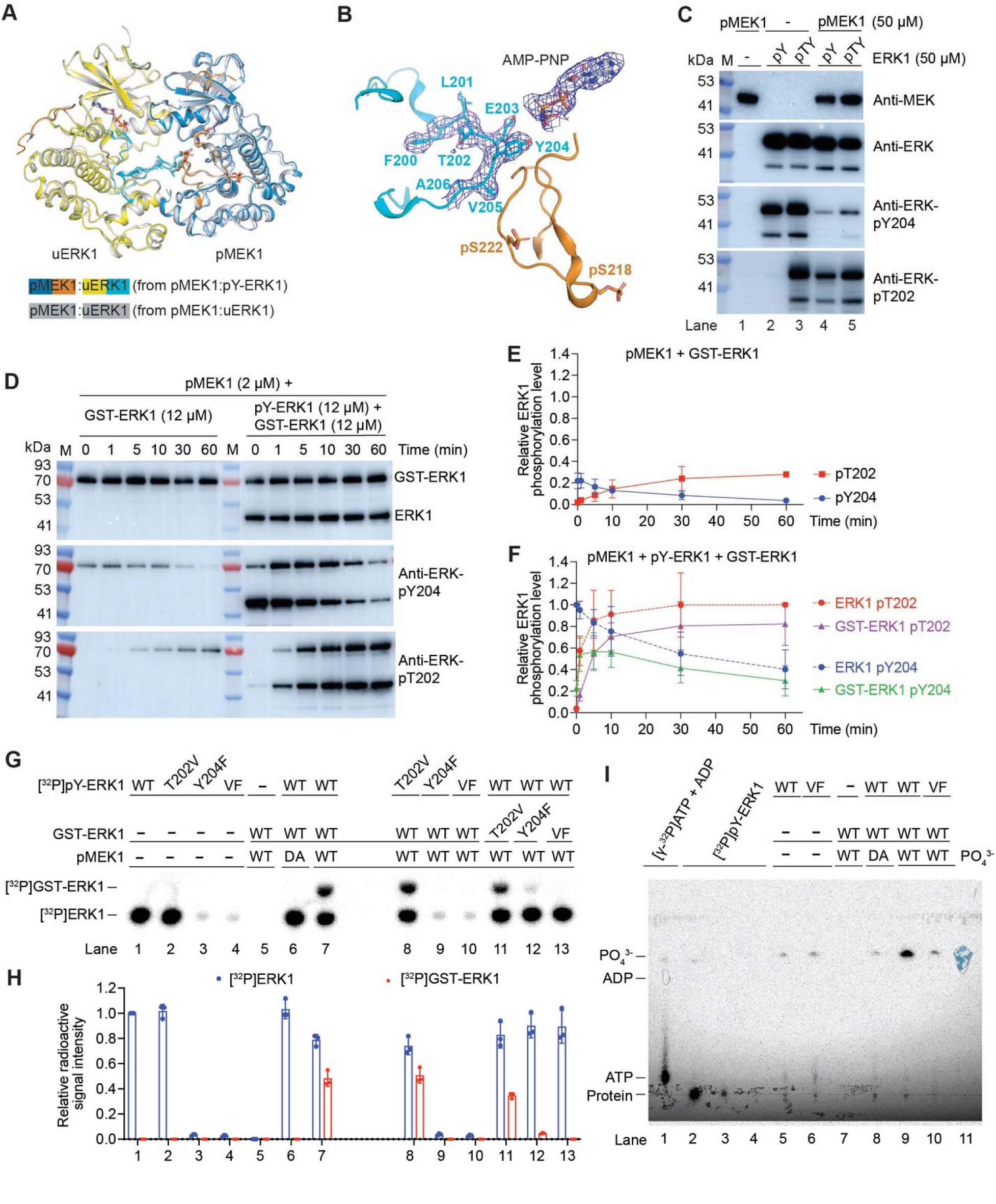
Phosphorylated MEK1 catalyzes ERK1 Y204 dephosphorylation. (A) Alignment of the cryo-EM structure of the pMEK1:uERK1 complex derived from the pMEK1:pY-ERK1 complex with the structure of the directly prepared pMEK1:uERK1 complex. (B) Confirmation of the activation loops of MEK1 and ERK1 in the pMEK1:uERK1 complex derived from the pMEK1:pY-ERK1 complex. (C) Dephosphorylation of pY-ERK1 and pTY-ERK1 by pMEK1 under the cryo-EM sample preparation condition, as evaluated by Western blotting for ERK1 phosphorylation. (D–F) pMEK1-catalyzed intramolecular and intermolecular phosphate group transfer from ERK1 Y204 to T202. In a nucleotide-free buffer, GST-ERK1 alone (slightly phosphorylated at ERK1 Y204) or together with pY-ERK1 was incubated with pMEK1 at 30 . The phosphorylation of ERK1 T202 and Y204 was evaluated using Western blotting (D) and quantified (E and F). (G and H) Evaluation of pMEK1-catalyzed phosphate transfer from pY-ERK1 to GST-ERK1 using a radioactive assay. ERK1 phosphorylated by [γ-^32^P]ATP at Y204 was incubated with unphosphorylated GST-ERK1 and wild-type (WT) or a catalytic-dead mutant (D208A, DA) mutant of pMEK1 in a nucleotide-free buffer for 5 min. Then the samples were analyzed by SDS–PAGE and subjected to radio imaging (G), and the radioactivity of each band was quantified (H). (I) Free phosphate (PO_4_ ^3-^) release following incubation of [^32^P]ERK1 with GST-ERK1 and pMEK1 for 5 min. After incubation, the samples were analyzed via thin-layer chromatography (TLC) and visualized by radio imaging. Lanes 1-4 and lane 11 are reference standards: [γ-^32^P]ATP and ADP mixture (lane 1), serially diluted [^32^P]ERK1 (lane 2-4), and PO_4_^3-^ (lane 11), respectively, which define the migration of ATP, ADP, proteins, and PO_4_^3-^ on the TLC plate. The results in (D), (G), and (I) are the results of a representative experiment out of three independent experiments. The data in (E), (F), and (H) represent the mean ± SD of three independent measurements.

Indeed, our Western blot data showed that under the condition used for preparing the cryo-EM samples (in the presence of 5 mM AMP-PNP), pY-ERK1 was almost completely dephosphorylated at Y204 by pMEK1 (Figure 4C). Under the same condition, T202/Y204 dual-phosphorylated ERK1 (pTY-ERK1) was also dephosphorylated at Y204 by pMEK1 (Figure 4C).

Interestingly, while pY-ERK1 was dephosphorylated at Y204, it was partially phosphorylated at T202 (Figure 4C, lane 4). We tested whether the phosphate transfer from ERK1 Y204 to T202 occurs intramolecularly or intermolecularly. We mixed pY-ERK1 with a GST-tagged ERK1 (GST-ERK1) in the presence of pMEK1 in a nucleotide-free buffer and then monitored the phosphate transfer. The GST-ERK1 purified from *E. coli* had a small fraction phosphorylated at Y204. When incubated alone with pMEK1, it was dephosphorylated at Y204 and phosphorylated at T202 (Figures 4D and 4E). When GST-ERK1 was incubated with pY-ERK1 and pMEK1, T202 in both GST-ERK1 and pY-ERK1 was gradually phosphorylated, and phosphorylation at Y204 in GST-ERK1 first increased, reached a plateau between 1 and 10 minutes, and then decreased (Figures 4D and 4F). The phosphorylation level of T202 in GST-ERK1 after incubation with pY-ERK1 was much higher than that in GST-ERK1 incubated without pY-ERK1 (Figures 4D–4F). Similar results were obtained when ERK1 was replaced by ERK2 (Figure S14). These results indicate that the phosphate group on ERK1 Y204 or ERK2 Y187 can be intermolecularly transferred to another ERK1 or ERK2 molecule.

We confirmed the intermolecular phosphate transfer using a radioactive assay. We prepared radiolabeled pY-ERK1 ([^32^P]pY-ERK1) by incubating uERK1 with [γ-^32^P]ATP, followed by removal of the excess [γ-^32^P]ATP using a desalting column (Figures 4G and 4H, lanes 1). We also incubated uERK1 mutants (T202V, Y204F, and the double mutant T202V/Y204F) with [γ-^32^P]ATP and found that radiolabeling was detected only in the T202V mutant (Figures 4G and 4H, lanes 2-4). Next, we incubated the radiolabeled pY-ERK1 with GST-ERK1 in a nucleotide-free buffer. GST-ERK1 was radiolabeled when incubated with wild-type or the T202V mutant of [^32^P]pY-ERK1, but not when incubated with the Y204F or T202V/Y204F double mutant (Figures 4G and 4H, lanes 7-10), indicating that [^32^P]pY204 in pY-ERK1 is the source of the radio signal transferred to GST-ERK1. The enzyme activity of pMEK1 is required for the radiolabeling of GST-ERK1, since it was not observed when a catalytic-dead mutant of pMEK1 (D208A) was used (Figures 4G and 4H, lanes 6).

The T202V mutant of GST-ERK1 was radiolabeled upon incubation with [^32^P]pY-ERK1 and pMEK1 for 5 min; the Y204F mutant was also radiolabeled, though with lower efficiency, and the T202V/Y204F double mutant was not radiolabeled under the same conditions (Figures 4G and 4H, lanes 11-13). The data are consistent with the Western blot data (Figure 4d) that the phosphate group on ERK1 Y204 was transferred to both Y204 and T202 of GST-ERK1, and at early time points, GST-ERK1 Y204 was phosphorylated faster than T202.

We also monitored the production of free phosphate using thin layer chromatography to track the movement of the radiolabeled phosphate group during ERK1 pY204 dephosphorylation (Figure 4I). Obvious production of free phosphate ([^32^P]PO_4_ ^3-^) was detected when [^32^P]pY-ERK1 was incubated with GST-ERK1 in the presence of the wild-type but not D208A mutant of pMEK1, suggesting that the dephosphorylation of ERK1 pY204 was accompanied by free phosphate production.

### ADP is used as the phosphate acceptor for pMEK1 to dephosphorylate ERK1 pY204

Previous studies have shown that protein phosphorylation catalyzed by some kinases (cAMP-dependent protein kinase, Src, EGFR, etc.) is reversible, with ADP acting as the phosphate acceptor^23–25^. Since ERK1 pY204 dephosphorylation was detected in a nucleotide-free buffer (Figures 4D–4H), we wonder whether trace amount of ADP present in the protein samples may contribute to this process. To test this, we used apyrase to hydrolyze ATP and ADP to AMP in the reaction system. Apyrase completely blocked both ERK1 pY204 dephosphorylation and T202 phosphorylation (Figure 5A). This finding supports the essential role of ADP in MEK1-catalyzed ERK1 dephosphorylation.

**Figure 5.**
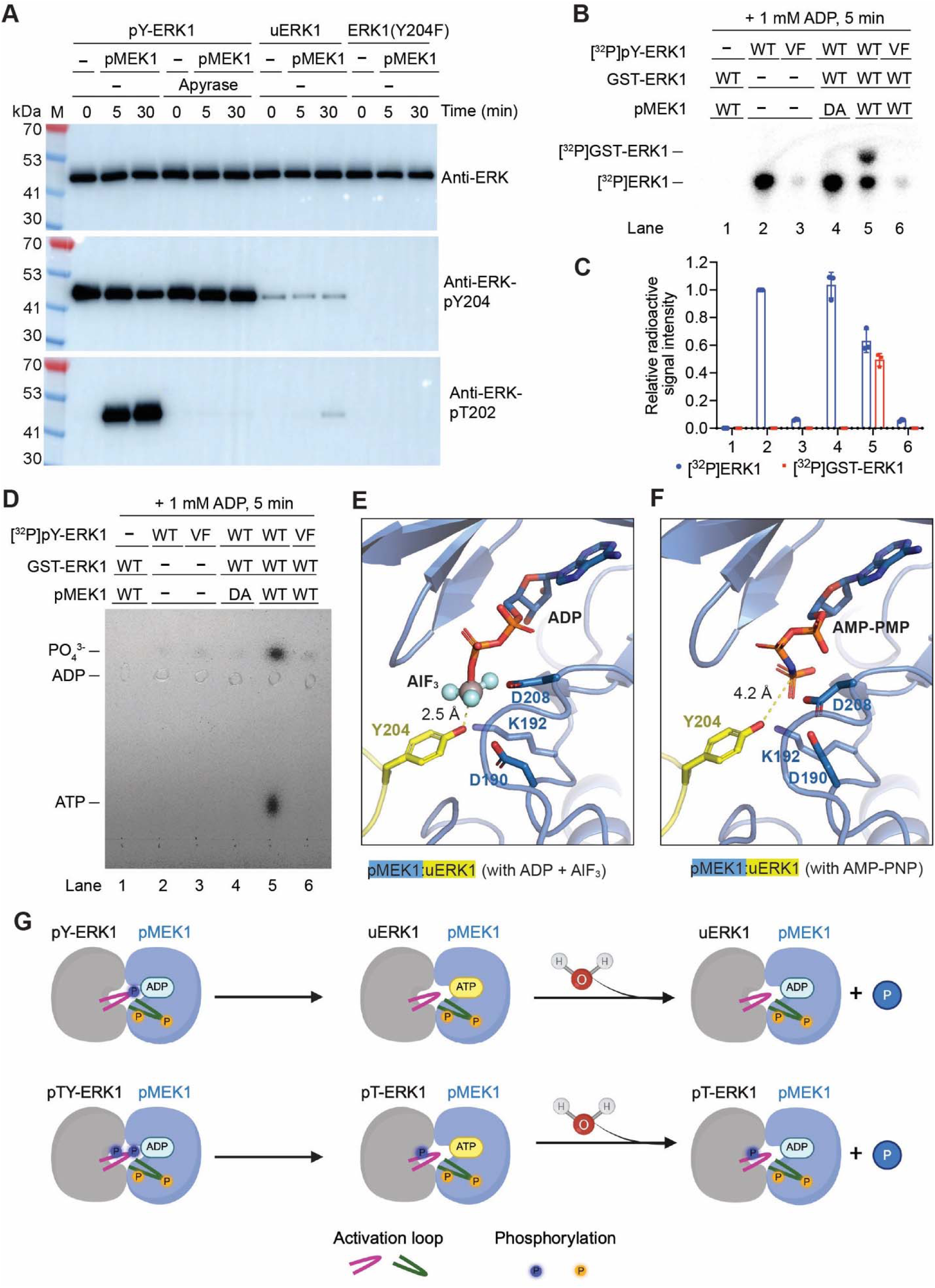
ADP is the phosphate acceptor for pMEK1 to dephosphorylate ERK1 pY204. (A) Removal of ADP blocked pMEK1-catalyzed dephosphorylation of ERK1 at Y204 and phosphate transfer to T202. In a nucleotide-free buffer, 12 µM pY-ERK1 was incubated with 2 µM pMEK1 at 30 ; at the indicated time points, the phosphorylation of ERK1 Y204 and T202 and was evaluated using Western blotting. (B–D) Evaluation of pMEK1-catalyzed ERK1 phosphate transfer using a radioactive assay in a buffer containing 1 mM ADP. The samples were analyzed by SDS–PAGE (B) or by TLC (D), and then subjected to radio imaging. The radioactivity of each band in (B) was quantified (C). The results in (A), (B) and (D) are the results of a representative experiment out of three independent experiments. The data in (C) represent the mean ± SD of three independent measurements. (E and F) Cryo-EM structure of the pMEK1:uERK1 complex bound to ADP and aluminum fluoride (AlF_3_) (E), or bound to AMP-PNP (F). (G) Illustration of pMEK1-catalyzed phosphate transfer from pY-ERK1 or pTY-ERK1 to ADP, followed by pMEK1-catalyzed hydrolysis of the newly generated ATP.

If ADP acts as the phosphate acceptor, dephosphorylation of ERK1 pY204 would generate ATP. However, we only detected production of [^32^P]PO_4_ ^3-^ but not [^32^P]ATP during ERK1 pY204 dephosphorylation in the nucleotide-free buffer (Figure 4I), probably because the newly generated ATP was below the detection limit. To verify ATP generation, we therefore added 1 mM ADP to the reaction system to drive ATP synthesis (Figures 5B–5D). Consistent with the results obtained in the nucleotide-free buffer, incubation of wild-type [^32^P]pY-ERK1 with GST-ERK1 and wild-type pMEK1 led to obvious dephosphorylation of [^32^P]pY-ERK1 accompanied by radiolabeling of GST-ERK1 (Figures 5B and 5C). Importantly, both [^32^P]PO_4_ ^3-^ and [^32^P]ATP were detected in this reaction (Figure 5D). These findings confirm that ADP functions as a phosphate acceptor to mediate ERK1 pY204 dephosphorylation.

We solved the cryo-EM structure of the pMEK1:uERK1 complex with ADP and aluminum fluoride (AlF_3_) to trap the transition state of reversible phosphate transfer between ERK1 Y204 and ADP (Table S1, Figures S5 and S15). The overall structure is almost identical to that of the pMEK1:uERK1 complex; however, in the transition state, the distance between the ERK1 Y204 hydroxyl group and AlF_3_ is 2.5 Å, which is closer than the 4.2 Å distance between the ERK1 Y204 hydroxyl group and the γ-phosphate of AMP-PNP bound to MEK1 in the pMEK1:uERK1 complex (Figures 5E and 5F). The mutations that disrupted the kinase activity of MEK1, including D190A, K192M, and D208A, also blocked the activity of MEK1 to catalyze the dephosphorylation of ERK1 pY204 (Figures S16A and S16B).

A previous study has shown that MEK1 exhibits ATPase activity^26^. We confirmed this activity in pMEK1 but not in uMEK1 (Figure S16C). These mutations that disrupted the kinase activity of pMEK1 also abolished its ATPase activity (Figure S16C). In addition, phosphorylated ERK1 also exhibited ATPase activity, but its activity was much lower than that of pMEK1. In contrast, the V600E mutant of BRAF, which is an active form of BRAF, showed negligible ATPase activity (Figure S16C).

Taken together, pMEK1 exhibits an unusual transferase activity to catalyze the reverse phosphorylation reaction, and a robust ATPase activity (Figure 5G). The ATP generated from ERK1 Y204 dephosphorylation can either be used to phosphorylate ERK1 Y204 and T202, or hydrolyzed by pMEK1, thereby explaining how ERK1 pY204 dephosphorylation and intermolecular phosphate transfer occur.

We further characterized the phosphatase-like activity of pMEK1 to understand its biological relevance. In the presence of 5 mM ADP, pMEK1 dephosphorylated pTY-ERK1 at Y204 in a concentration-dependent manner (Figures S17A and S17B). As ATP and ADP exist as a mixture in cells, we next examined how varying ATP ratios regulate the balance between the kinase and phosphatase-like activities of pMEK1. In an ATP–ADP-containing buffer with a fixed total nucleotide concentration of 5 mM, 1 µM pTY-ERK1 or pTY-ERK2 was incubated with 200 nM pMEK1 at 30 °C for 30 min. The ratio of pTY-ERK1 or pTY-ERK2 relative to total ERK1 or ERK2 dropped quickly once the ATP ratio fell below 25% (Figure S17C). Notably, when DUSP6, one of the phosphatases that dephosphorylate ERK1/2 in the cytosol^27^, was added to the reaction, pTY-ERK1 ratio became more sensitive to changes in the ATP ratio: pTY-ERK1 ratio decreased sharply once the ATP ratio fell below 50%, suggesting an additive effect of DUSP6 on the phosphatase-like activity of pMEK1 (Figure S17D). In the presence of DUSP6, when ADP in the buffer was replaced with AMP-PNP, pTY-ERK1 ratio decreased only when the ATP ratio was below 30% (Figure S17D), supporting the role of ADP in promoting pTY-ERK1/2 dephosphorylation. Cytosolic ATP is usually about 10-fold more abundant than ADP; however, under certain pathological conditions, such as hypoxia and ischemia^28,29^, the cytosolic ATP concentration can decrease to a level equal to or even lower than that of ADP, and pMEK1 may function as an ATP/ADP ratio sensor under such conditions.

### Structural basis for ERK1 T202 phosphorylation

After elucidating the mechanism of pMEK1-catalyzed ERK1 dephosphorylation, we attempted again to solve the structure of pMEK1 in complex with pY-ERK1 in the presence of AMP-PNP, aiming to capture the structure of ERK1 in which T202 is positioned to the MEK1 catalytic pocket for phosphorylation. By carefully controlling the incubation time of pMEK1 with pY-ERK1, we finally solved the structure of pMEK1 in complex with pY-ERK1 and AMP-PNP (Table S1, Figures S2 and S18). The overall structure of the pMEK1:pY-ERK1 complex closely resembles that of the pMEK1:uERK1 complex (Figure 6A). The activation loop of pY-ERK1 adopts a conformation almost identical to that of unphosphorylated ERK1. The side chain of phosphorylated Y204, but not unphosphorylated T202, in ERK1 points to the γ-phosphate of MEK1-bound AMP-PNP (Figure 6B).

**Figure 6.**
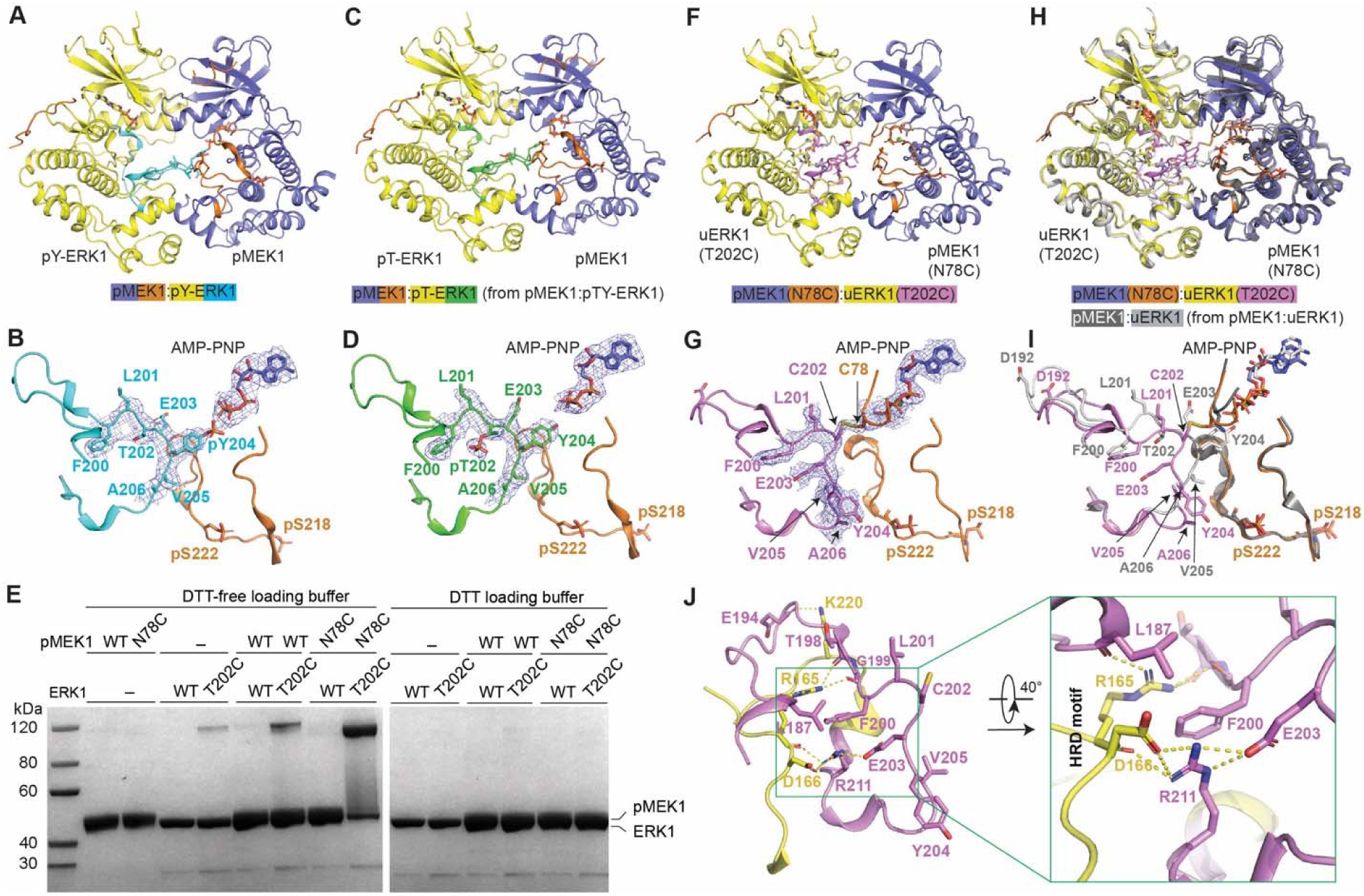
Structural basis for ERK1 T202 phosphorylation. (A and B) Overall structure of the pMEK1:pY-ERK1 complex bound to AMP-PNP (A) and conformation of the activation loops of MEK1 and ERK1 in this structure (B). (C and D) Overall structure of the pMEK1:pT-ERK1 complex bound to AMP-PNP (derived from the pMEK1:pTY-ERK1 complex) (C) and conformation of the activation loops of MEK1 and ERK1 in this structure (D). (E) Crosslinking of pMEK1(N78C) with uERK1(T202C) in the presence of 0.4 mM glutathione disulfide (GSSG). (F and G) Overall structure of the crosslinked pMEK1(N78C):uERK1(T202C) complex bound to AMP-PNP (F) and conformation of the activation loops of MEK1 and ERK1 in this structure (G). The cryo-EM maps of ERK1 F200-V205 and AMP-PNP bound to pMEK1 are contoured at 5.0 σ and shown as blue meshes. (H) Alignment of the cryo-EM structure of the crosslinked pMEK1(N78C):uERK1(T202C) complex with the structure of the pMEK1:uERK1 complex. (I) Comparison of the conformation of the ERK1 activation loop in the pMEK1(N78C):uERK1(T202C) complex with that in the wild-type pMEK1:uERK1 complex. (J) Interactions between the ERK1 activation loop and the ERK1 HRD motif in the pMEK1(N78C):uERK1(T202C) complex.

We also solved the structure of pMEK1 in complex with T202 mono-phosphorylated ERK1 (pT-ERK1). We attempted to solve the structure of the pMEK1:pTY-ERK1 complex, but finally found that it yielded a cryo-EM structure of pMEK1 in complex with pT-ERK1 (Table S1, Figures S2 and S19). The pMEK1:pT-ERK1 structure also closely resembles the structure of the pMEK1:uERK1 complex (Figure 6C). Phosphorylated ERK1 T202 remains distant from the γ-phosphate of MEK1-bound AMP-PNP, while the side chain of Y204 is in a position ready for phosphorylation (Figure 6D).

To overcome the difficulty in capturing the conformation of ERK1 T202 positioned toward the MEK1 catalytic pocket, we employed a crosslinking strategy. In the pMEK1:uERK1 complex, MEK1 N78 is at the entrance of the MEK1 catalytic pocket and is close to ERK1 Y204 (Figure 2B). We reasoned that ERK1 T202 should also be close to MEK1 N78 during its phosphorylation. Therefore, we mutated ERK1 T202 and MEK1 N78 to cysteine residues, and tested whether they could be crosslinked via a disulfide bond (Figure 6E). ERK1(T202C) and MEK1(N78C) each alone showed only a small fraction crosslinked in the presence of 0.4 mM glutathione disulfide (GSSG). When mixed together, ERK1(T202C) was almost completely crosslinked with pMEK1(N78C) (Figure 6E). These findings suggest that the ERK1 activation loop is in a dynamic state and T202 can be close to MEK1 N78.

Next, we solved the cryo-EM structure of the crosslinked pMEK1(N78C):uERK1(T202C) complex in an AMP-PNP-bound state at 2.33 Å resolution (Table S1, Figures S5 and S20). As anticipated, ERK1 C202 forms a disulfide bond with MEK1 C78 (Figures 6F and 6G). Structural alignment shows that the overall structure is quite similar to that of the pMEK1:uERK1 complex, with the major differences located in residues D192 to A206 in the activation loop of ERK1 (Figures 6H and 6I). ERK1 T202 points toward the ERK1 nucleotide-binding pocket in the pMEK1:uERK1 structure; by contrast, after T202 is mutated to a cysteine residue and crosslinked with pMEK1(N78C), it faces the MEK1 nucleotide-binding pocket (Figure 6I). Accompanied by this conformational change, ERK1 E203 flips about 180° and Y204 shifts away from the catalytic center of MEK1 (Figure 6I). These conformational changes explain how the ERK1 activation loop rearranges to enable T202 phosphorylation.

The conformation of the ERK1 activation loop in the crosslinked structure is stabilized via interactions with R165 and D166 in the HRD motif of ERK1 (Figure 6J). Specifically, the side chain of ERK1 R165 forms hydrogen bonds with the main chains of L187, T198 and G199, while both the side chain and main chain of ERK1 D166 form hydrogen bonds with the side chain of ERK1 R211. ERK1 E203, which undergoes a significant conformational change relative to its conformation in the pMEK1:uERK1 structure, also forms hydrogen bonds with ERK1 R211. Furthermore, the side chain of ERK1 F200 sits between the side chains of R165 and R211 to mediate strong cation-π interactions. These interactions are distinct from those stabilize the ERK1 activation loop in the pMEK1:uERK1 structure, in which this loop is stabilized mainly through interactions with the MEK1 activation loop (Figure 2D).

### ATP binding drives the MEK1-catalyzed ERK1 phosphorylation cycle

To get a full picture of the MEK1-catalyzed ERK1 phosphorylation cycle, we sought to elucidate how this catalytic cycle is driven by ATP binding, ERK1 phosphorylation, and ADP dissociation. To address this, we measured the binding affinities between MEK1 and unphosphorylated, mono-phosphorylated or dual-phosphorylated ERK1 in the presence and absence of AMP-PNP or ADP (Figures 7A and S21).

**Figure 7.**
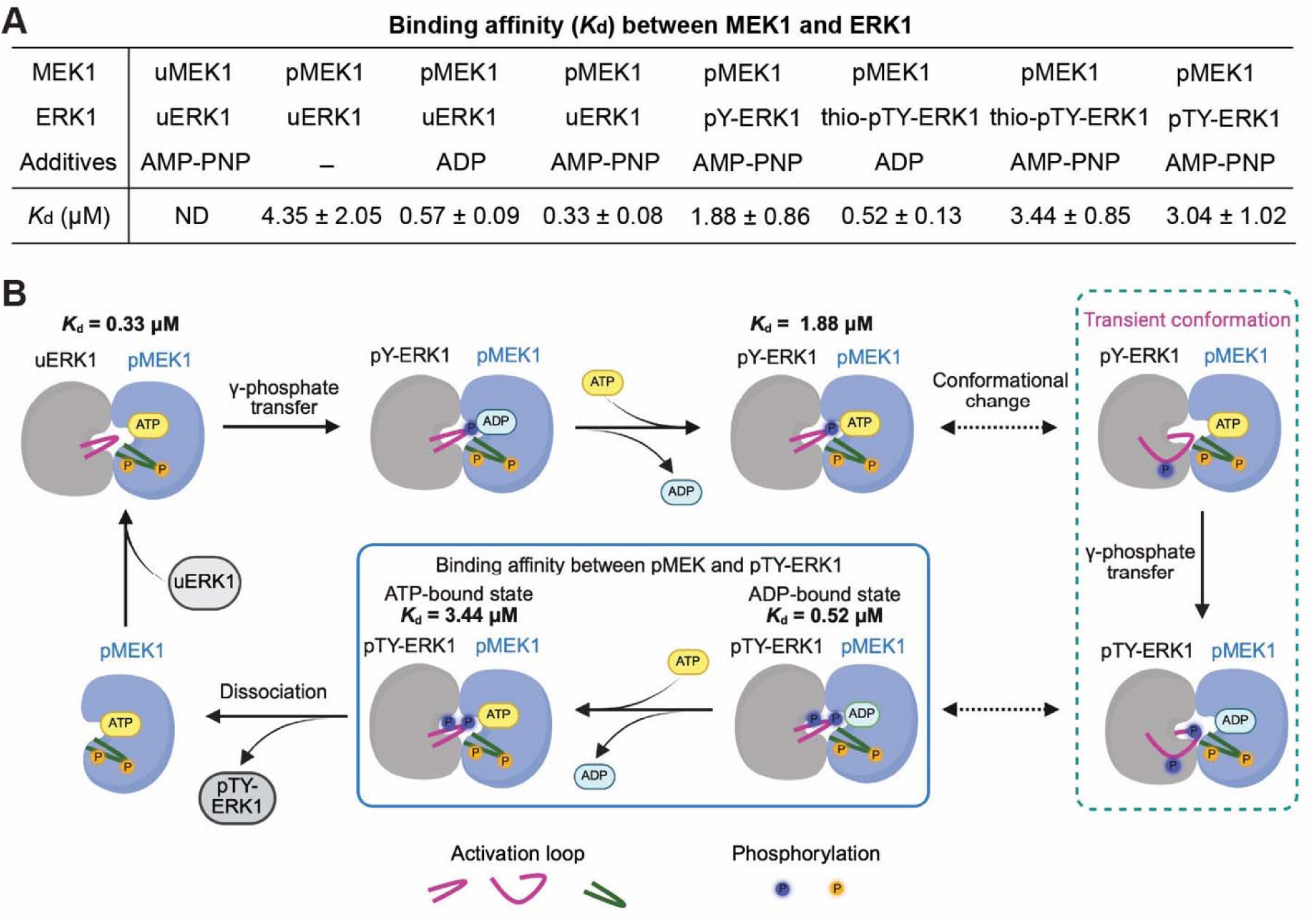
Catalytic cycle of the MEK1-catalyzed ERK1 phosphorylation reaction. (A) Summary of the dissociation constants (*K*_d_) for the binding between MEK1 (uMEK1 or pMEK1) and ERK1 (uERK1, pY-ERK1, thio-pTY-ERK1, or pTY-ERK1) in the absence or presence of ADP or AMP-PNP. The *K*_d_ values were measured using Isothermal Titration Calorimetry (ITC). (B) Illustration of the cycle of the pMEK1-catalyzed ERK1 phosphorylation reaction.

The interaction between uMEK1 and uERK1 was undetectable using isothermal titration calorimetry (ITC) (Figures 7A and S21), consistent with the gel filtration assay data (Figure S1). In contrast, pMEK1 binds to uERK1 with a dissociation constant (*K*_d_) of 4.35 µM in the absence of ADP or AMP-PNP, and their binding affinity increases by about 10-fold in the presence of AMP-PNP (*K*_d_ = 0.33 µM). The binding between pMEK1 and uERK1 can also be enhanced by ADP (*K*_d_ = 0.57 µM). Using apyrase to remove residual ADP, we measured the *K*_d_ between pMEK1 and pY-ERK1 in the presence of AMP-PNP; the value is 1.88 µM, which is higher than that between pMEK1 and uERK1 in the presence of AMP-PNP, suggesting weaker binding. The *K*_d_ between pMEK1 and pY-ERK1 in the presence of ADP was not measured, since pY-ERK1 can be quickly dephosphorylated by pMEK1 when ADP is present.

To measure the binding affinity between pMEK1 and pTY-ERK1 in the presence of either ADP or AMP-PNP, we prepared ERK1 thiophosphorylated at T202 and Y204 instead (thio-pTY-ERK1) by incubating uERK1 with an active form of MEK1 in an ATPγS-containing buffer. Thio-pTY-ERK1 has similar kinase activity to pTY-ERK1 (Figures S22A and S22B), but is less susceptible to being dephosphorylated by pMEK1 in comparison with pTY-ERK1 (Figures S22C and S22D). The *K*_d_ is 0.52 µM in the presence of ADP, but increases to 3.44 µM in the presence of AMP-PNP (Figures 7A and S21). Using apyrase to remove residual ADP, we also measured the *K*_d_ between pMEK1 and pTY-ERK1 in the presence of AMP-PNP; the value is 3.04 µM, which is similar to that between pMEK1 and thio-pTY-ERK1.

These binding measurements indicate that nucleotide-bound state and ERK1 phosphorylation status coordinately regulate the assembly and disassembly of the MEK1–ERK1 complex throughout catalysis. Based on the above data, we delineated the catalytic cycle of the MEK1-catalyzed ERK1 phosphorylation reaction (Figure 7B). Firstly, phosphorylated MEK1 recruits uERK1, with their binding affinity enhanced by ATP, and then ERK1 Y204 is phosphorylated by accepting the γ-phosphate from ATP, forming the pMEK1:pY-ERK1:ADP complex. Secondly, ADP in pMEK1 is replaced by ATP, and the ERK1 activation loop undergoes conformational changes to orient T202 toward the MEK1 catalytic center, and then ERK1 T202 is phosphorylated by accepting the γ-phosphate from ATP. Lastly, ADP in the pMEK1:pTY-ERK1:ADP complex is replaced by ATP, and the binding affinity between pMEK1 and pTY-ERK1 decreases 6.6-fold, triggering the dissociation of pTY-ERK1 from pMEK1. Then pMEK1 can enter another catalytic cycle.

## DISCUSSION

The RAS-RAF-MEK-ERK cascade is a major focus of cancer research due to its critical role in controlling cancer development and progression. Structural biology studies have revealed how RAS binds to RAF, how RAF is activated by phosphorylation, and how RAF binds to MEK. However, the mechanisms by which MEK binds to and catalyzes ERK phosphorylation remain poorly understood. We filled this gap by solving the cryo-EM structures of phosphorylated MEK1 in complex with ERK1 in different phosphorylation states and with unphosphorylated ERK2, as well as the structure of pMEK1(N78C) crosslinked with unphosphorylated ERK1(T202C), and by performing phosphorylation assays to investigate the sequential phosphorylation of ERK1 Y204 and T202.

These cryo-EM structures share a quite similar overall structure. The phosphorylated MEK1 interacts with ERK1/2 primarily through two interfaces. The first interface is between the N-terminal peptide of MEK1 and the D-site recruitment site (DRS) in ERK1; the second is between the A309-E320 helix in the C-lobe of MEK1 and the F-site recruitment site (FRS) in ERK1/2 (Figure 1). These highly specific interactions explain the high specificity of MEK1/2 towards ERK1/2. The cryo-EM structures also reveal how MEK1 undergoes conformational changes for activation. Unlike the activation of ERK1/2, which mainly involves conformational changes in the activation loop of ERK1/2, the activation of MEK1 involves large conformational changes in three regions: the activation loop, the αC-helix, and the αA-helix (Figure 3). These conformational changes, together with the binding of the MEK1 N-terminal peptide to the ERK1 DRS, allow the catalytic pocket of MEK1 to achieve its active conformation.

More importantly, these structures reveal the structural basis for the dual phosphorylation of ERK1 catalyzed by MEK1. In all these structures except that of the crosslinked pMEK1(N78C):uERK1(T202C) complex, the ERK1/2 activation loops interact with the MEK1 activation loop, positioning the side chain hydroxyl group of ERK1 Y204 or ERK2 Y187 to the γ-phosphate of AMP-PNP bound to MEK1, explaining how the tyrosine residues are phosphorylated. This structural preference is consistent with the report that in the ERK1/2 activation loops, the tyrosine residues are phosphorylated first, followed by the threonine residues^10^. In contrast, in the pMEK1(N78C):uERK1(T202C) complex, the ERK1 activation loop, instead of interacting with the activation loop of MEK1, is stabilized by the ERK1 HRD motif, and consequently, adopts a distinct conformation to orient C202, the substituent residue of T202, to the catalytic center of MEK1 (Figure 6F–6J). Taken together, these structures demonstrate how ERK1 dynamically rearranges its activation loop to enable phosphorylation at Y204 and T202, and explain the dual specificity of MEK1.

While this manuscript was under submission, von Velsen *et al*. deposited a preprint on bioRxiv that reported cryo-EM structures of a chimeric MEK1 in complex with the ERK2 T185V mutant^30^. Surprisingly, these structures are quite different from those of the MEK1:ERK1/2 complexes solved in our study. We compared one of their structures (PDB: 9TYH), which was reported as the active state, with the structure of the pMEK1:uERK2 complex from our study, and found that when C-lobe of ERK2 in the two structures are aligned, the kinase domain of the chimeric MEK1 in 9TYH is rotated by 39° relative to that of MEK1 in the pMEK1:uERK2 complex (Figure S23A–S23C). Furthermore, when the C-lobes of MEK1 in the two structures are aligned, the αC-helix of MEK1 in 9TYH is rotated by 8° relative to that in the pMEK1:uERK2 complex, and the β-sheet in the N-lobe of MEK1 is rotated to by 7° to the opposite direction (Figure S23D–S23F). In addition, most residues in the MEK1 activation loop in 9TYH are invisible, in contrast to the well-resolved MEK1 activation loop in our pMEK1:uERK2 structure. (Figure S23D-S23F). One explanation for these differences is that von Velsen *et al*. used an engineered MEK1, in which the N-terminal peptide of MEK1 was replaced with the kinase interaction motif (KIM) from the *Toxoplasma gondii* effector protein GRA24 to increase the binding affinity with ERK2, and S218 and S222 in MEK1 were mutated to aspartate to mimic the phosphorylated state of the two residues. Such engineering may distort the interactions between MEK1 and ERK2. In contrast, we used wild-type MEK1 phosphorylated by BRAF instead of incorporating phosphomimetic mutations. Our structures therefore represent the native states of MEK1 in complex with ERK1/2.

The structural basis for the phosphorylation of MAPKs by MAP kinase kinases (MAP2Ks) has also been studied by Juyoux *et al*. through characterization of an engineered MKK6:p38α(T180V) complex with cryo-EM and molecular dynamics (MD) simulations^22^. In comparison with this previous study, we have two different findings. First, in the cryo-EM structures of the pMEK1:ERK1 complexes, the ERK1 activation loop is stabilized by the MEK1 activation loop to enable phosphorylation of ERK1 Y204 (Figure 2D), and stabilized by the ERK1 HRD motif to enable phosphorylation of ERK1 T202 (Figure 6J). In contrast, in the cryo-EM structure of the chimeric MKK6:p38α(T180V) complex, the activation loop of MKK6 is only partially visible, and that of p38α has a poor resolution, leading Juyoux *et al*. to conclude that the loop regions lack tight interactions^22^. Second, from the conformation for Y204 phosphorylation to the conformation for T202 phosphorylation, the overall structure of ERK1 and its interaction with pMEK1 remain largely unchanged, with the only significant conformational changes in the ERK1 activation loop (Figures 6H and 6I). In contrast, Juyoux *et al*. did MD simulations and found that for p38α T180 (corresponding to ERK1 T202) to approach the catalytic center of MKK6, the p38α N-lobe rotated about 50 to 80° relative to its C-lobe^22^. It should be noted that Juyoux *et al.* used a strategy similar to that used by von Velsen *et al*. for MEK1:ERK2 structure determination to obtain the MKK6:p38α(T180V) structure^22^. The native KIM in MKK6 was replaced with the KIM from the *Toxoplasma gondii* effector protein GRA24, and S207 and T211 of MKK6 were mutated to aspartate to stabilize the MKK6:p38α(T180V) complex. It is important to solve structures of the native MKK6:p38α complex to clarify whether the differences between the MAPK phosphorylation mechanism revealed by our study and that revealed by the chimeric MKK6:p38α(T180V) structure are due to the intrinsic differences between MKK6:p38α and MEK1:ERK1/2 or due to the protein engineering of MKK6.

Another important finding of our study is that pMEK1 can catalyze the dephosphorylation of ERK1 at Y204 (and ERK2 at Y187) by transferring the phosphate group to ADP to generate ATP (Figures 4 and 5). The reversible phosphorylation activity has been observed in several other kinases^23–25^; however, considering the high ATP/ADP ratio in cells, the biological significance of the reverse phosphorylation remains unclear. We showed that together with the unexpected reverse phosphorylation activity, pMEK1 exhibits robust ATPase activity (Figure S16C). Therefore, pMEK1 transfers the phosphate group from ERK1 to ADP to generate ATP and subsequently catalyzes ATP hydrolysis. Through the two sequential reactions, pMEK1 drives ERK1 dephosphorylation to release free phosphate, thereby exhibiting phosphatase-like activity. We also showed that the phosphatase-like activity of pMEK1 can be additive with the phosphatase activity of DUSP6, enabling pMEK1 to exert phosphatase-like activity at a higher ATP/ADP ratio (Figure S17D). Therefore, we speculate that pMEK1, in coordination with ERK1/2 phosphatases, may function as an ATP/ADP ratio sensor under certain pathological conditions to regulate cell signaling.

MEK1/2 are important anti-cancer drug targets. Several inhibitors of MEK1/2, such as trametinib, cobimetinib, and selumetinib, have been approved for treating cancers with abnormal activation of the RAS-RAF-MEK-ERK cascade^31^. All of them are non-ATP-competitive inhibitors. Based on our findings that pMEK1 can catalyze the dephosphorylation of ERK1/2 and that ADP, but not AMP-PNP, acts as the phosphate group acceptor, we propose that developing MEK1/2 inhibitors to mimic the effect of ADP, which accepts phosphate groups from substrate proteins, would be a promising strategy targeting the RAF-MEK-ERK cascade. Such inhibitors may abolish the kinase activity of MEK1/2 while maintaining their phosphatase activity to inactivate ERK1/2. This strategy should be applicable to inhibit other kinases that can also catalyze the reverse phosphorylation.

## Acknowledgments

We would like to thank Profs. Hongtao Yu, Kevan M. Shokat and Jing Huang for providing suggestions on this study; the Cryo-EM Facility and HPC Center of Westlake University for providing cryo-EM and computation support, the Mass Spectrometry & Metabolomics Core Facility at the Center for Biomedical Research Core Facilities of Westlake University for sample analysis. This work was supported by Central Guidance on Local Science and Technology Development Fund of Zhejiang Province (2026ZY01027), State Key Laboratory of Gene Expression, Westlake Education Foundation, and “Pioneer” and “Leading Goose” R&D Program of Zhejiang (2023C03109 and 2024SSYS0036).

## Author Contributions Statement

Q.H. conceived and supervised the project; Y.S. and C.P. purified the proteins, performed the biochemical assays; Y.S., C.P. and F.Z. prepared the cryo-EM samples and collected the cryo-EM data; S.L. calculated the cryo-EM map with the supervision of G.H.; C.P. refined the cryo-EM structures; J.W. provided supervision to Y.S.; all authors contributed to data analysis; Q.H. wrote the manuscript with inputs from other authors.

## Data availability

The cryo-EM structures of human MEK1:ERK1/2 complexes have been deposited in the Protein Data Bank with the accession codes 9UUR, 9UW4, 9UW3, 9UUX, 25GJ, 25HU and 43YB. The cryo-EM maps have been deposited in the Electron Microscopy Data Bank with the accession codes EMD-64516, EMD-64545, EMD-64544, EMD-64519, EMD-80086, EMD-80120 and EMD-82382. All other data are available in the manuscript or the supplementary materials.

## Materials and Methods

### Cell culture

The Expi293F cells were cultured in SMM293-TII medium (Sino Biological, Cat# M293TII) supplemented with 1% penicillin-streptomycin solution (Solarbio, Cat# P1400) at 37 °C under 5% CO_2_ in a shaking incubator at 120 RPM.

### Genes and Cloning

The constitutively active mutation V600E was introduced into human BRAF (accession number in PubMed: NP_004324.2), and the gene encoding human BRAF(V600E) with an N-terminal FLAG tag and an 8×His tag was cloned into a pcDNA3.1 vector. The gene encoding human MEK1 (accession number in PubMed: NP_002746.1) without stop codon was cloned into a modified pET21b vector. A constitutively active form of MEK1 (residues 32-51 truncated, S218E/S222D), also called ΔN3-MEK1-SESD, was cloned into both a pBB75 vector and a modified pET21b vector. The genes encoding human ERK1 (accession number in PubMed: NP_002737.2) and ERK2 (accession number in PubMed: NP_002736.3) with or without a C-terminal 6× His tag were cloned into a pGEX vector. Site directed mutagenesis was performed to generate the expression constructs of MEK1 and ERK1 mutants. The gene encoding human DUSP6 (accession number in PubMed: NP_001937.2) with a C-terminal 6×His tag was cloned into a pGEX vector for protein expression.

### Protein expression and purification

Human BRAF(V600E) was transiently expressed in Expi293F cells and purified by Anti-FLAG Affinity Resin (GenScript, Cat# L00432-25). The 293F cells at a density of 2× 10^6^ cells/mL in SMM293-TII medium were transfected with 1-1.5 mg per liter of the pcDNA3.1 plasmid and linear Polyethylenimine 25,000 (DNA mass: PEI mass = 1:2). The transfected cells were cultured for 48 hours, and then harvested by centrifugation at 4,000 g for 10 min at 4 °C. The cell pellets were resuspended in a lysis buffer (25 mM Tris–HCl pH 8.0, 150 mM NaCl, 5% glycerol) supplemented with 1.3 µg/mL Aprotinin (VWR, Cat# E429), 1 µg/mL Pepstain (VWR, Cat# J583), 5 µg/mL Leupeptin (VWR, Cat# J580), 0.5% *v*/*v* Triton X-100 and 1 mM PMSF (Macklin, Cat# P6140), and lysed by ultrasonication. The cell lysates were centrifuged at 22,000 g at 4 °C for 1 hour (BECKMAN Avanti JXN-26 Centrifuge), and then the target protein in the supernatants was purified using anti-FLAG affinity resin following the manufacturer’s protocol. The eluate was supplemented with 5 mM ATP (Macklin, Cat# A6363) and 5 mM MgCl_2_, then the sample was further purified using TALON Metal Affinity Resin (TaKaRa, Cat# 635504) to remove HSP70.

The C-terminal His-tagged wild-type MEK1 and its mutants, and the constitutively active form of MEK1 (ΔN3-MEK1-SESD-His), were each transformed separately into *E. coli* BL21(DE3) cells. The cells were cultured in LB medium supplemented with 0.1 mg/mL ampicillin at 37 °C until OD600 reached 1-1.5, then cooled to 18 °C followed by addition of 200 µM isopropyl β-D-thiogalactopyranoside (IPTG). Then the cells were cultured at 18 °C overnight. The cells were harvested by centrifugation at 4,000 g for 10 min at 4 °C and resuspended in the lysis buffer supplemented with 1 mM PMSF and lysed by ultrasonication. The cell lysates were centrifuged at 22,000 g at 4 °C for 1 hour, then the target proteins in the supernatants were purified using TALON Metal Affinity Resin following the manufacturer’s protocol.

The wild-type and mutant ERK1/2, and DUSP6 with a GST tag at their N-terminus and a 6×His tag at their C-terminus were also separately overexpressed in BL21(DE3) and purified by using TALON Metal Affinity Resin. Then the eluates were supplemented with 5 mM dithiothreitol (DTT) and treated with DrICE enzyme to remove the GST tag. Then the samples were further purified using Glutathione Sepharose 4B resin (GE Healthcare, Cat# 17075605) to remove the GST tag following the manufacturer’s protocol. ERK1/2 with only a GST tag at its N-terminus (GST-ERK1/2) was overexpressed in BL21(DE3) and purified using Glutathione Sepharose 4B resin following the manufacturer’s protocol.

All the eluates from affinity resins were supplemented with 5 mM Dithiothreitol (DTT) and loaded into a Source-15Q anion exchange column (GE Healthcare) and eluted by a linear gradient from 100% buffer A (25 mM Tris–HCl pH 8.0, 5 mM MgCl_2_, 5% glycerol) to 55% buffer B (25 mM Tris–HCl pH 8.0, 1 M NaCl, 5 mM MgCl_2_, 5% glycerol). The peak fractions were pooled, supplemented with 5 mM DTT, concentrated using a centrifugal filter (Amicon Ultra, 30 kDa MWCO) and further purified using Superdex 200 Increase 10/300 GL column (GE Healthcare) and eluted by SEC buffer (25 mM HEPES pH 7.4, 150 mM NaCl, 5 mM MgCl_2_). The peak fractions were pooled for biochemical assays and cryo-EM sample preparation.

### Preparation of phosphorylated MEK1 (pMEK1)

Phosphorylated MEK1 and its mutants were prepared by incubation with BRAF(V600E). The wild-type or each MEK1 mutant was mixed with purified BRAF(V600E) in a reaction buffer containing 50 mM Tris–HCl pH 8.0, 4.35% glycerol, 1 mM DTT, 150 mM NaCl, 5 mM ATP, and 10 mM MgCl_2_. The final concentrations of MEK1 and BRAF(V600E) in the reaction mixture were 10 µM and 0.5 µM, respectively. The reaction was carried out at 4 °C and monitored by LC-MS. Once the protein was nearly completely phosphorylated at both S218 and S222 sites, the reaction mixture was loaded into a Source-15Q column and eluted by a linear gradient from 100% buffer A to 55% buffer B. The phosphorylation state of MEK1 in each fraction was confirmed by LC-MS. The fractions corresponding to the pMEK1 peak were pooled, supplemented with 5 mM DTT, concentrated using a centrifugal filter and further purified using Superdex 200 Increase 10/300 GL column and eluted by SEC buffer (25 mM HEPES pH 7.4, 150 mM NaCl, 5 mM MgCl_2_). The peak fractions were pooled for biochemical assays and cryo-EM sample preparation.

### Preparation of phosphorylated ERK1 at Y204 or ERK2 at Y187 (pY-ERK1 or pY-ERK2)

The C-terminal His-tagged ERK1/2 (uERK1/2) purified via a Source-15Q column was concentrated to 600 µM and then mixed with an ATP-containing buffer to achieve a final concentration of 300 µM ERK1/2, 25 mM Tris–HCl pH 8.0, 2.5% glycerol, 150 mM NaCl, 20 mM ATP, 40 mM MgCl_2_, and 5 mM DTT in the reaction mixture. The mixture was incubated at room temperature. The phosphorylation level of ERK1/2 was monitored by LC-MS. Once the protein was nearly completely mono-phosphorylated, the reaction mixture was concentrated using a centrifugal filter and desalted using a HiTrap Desalting column equilibrated with SEC buffer to remove residual ATP or ADP. The desalted protein was then loaded onto a Source-15Q anion exchange column and eluted with a linear gradient from 100% buffer A to 40% buffer B to separate pY-ERK1/2 from residual uERK1/2. The phosphorylation state of ERK1/2 in each fraction was confirmed by LC-MS. The fractions corresponding to the pY-ERK1/2 peak were pooled, supplemented with 5 mM DTT, and further purified using a Superdex 200 Increase 10/300 GL size-exclusion chromatography column equilibrated with SEC buffer. The peak fractions were pooled for biochemical assays and cryo-EM sample preparation.

### Preparation of ERK1 phosphorylated at Y204 and T202 (pTY-ERK1)

The GST-ERK1-His in a pGEX vector and FLAG-MEK-ΔN3-SESD in a pBB75 vector were co-transformed into BL21(DE3) cells. The transformed cells were grown in LB medium supplemented with 50 µg/mL ampicillin and 25 µg/mL kanamycin at 37 °C until the OD600 reached 1.5. The cultures were then cooled to 18 °C for 1 hour, followed by the addition of 200 µM IPTG. The cells were cultured overnight at 18 °C. The cells were harvested by centrifugation at 4,000 g for 10 min at 4 °C and resuspended in the lysis buffer supplemented with 1 mM PMSF and lysed by ultrasonication. The cell lysates were centrifuged at 22,000 g at 4 °C for 1 hour, then ERK1 in the supernatants was purified using TALON Metal Affinity Resin following the manufacturer’s protocol. Next, the eluate was supplemented with 5 mM DTT and treated with DrICE enzyme to cleave the GST tag. Then the sample was further purified using Glutathione Sepharose 4B resin (GE Healthcare, Cat# 17075605) to remove the GST tag following the manufacturer’s protocol.

The flow through was loaded into a Source-15Q column (GE Healthcare) and eluted by a linear gradient from 100% buffer A to 40% buffer B to separate pTY-ERK1 from uERK1 and pY-ERK1. The phosphorylation state of ERK1 in each fraction was confirmed by LC-MS. The fractions containing pTY-ERK1 were pooled, supplemented with 5 mM DTT, and further purified using a Superdex 200 Increase 10/300 GL size-exclusion chromatography column equilibrated with SEC buffer. The peak fractions were pooled for biochemical assays and cryo-EM sample preparation.

### Preparation of ERK2 at Y187 and T185 phosphorylated (pTY-ERK2)

The uERK2 purified via a Source-15Q column was mixed with MEK-ΔN3-SESD in an ATP-containing buffer to achieve a final concentration of 100 µM ERK2, 3 µM MEK-ΔN3-SESD, 25 mM Tris–HCl pH 8.0, 5% glycerol, 150 mM NaCl, 10 mM ATP, 20 mM MgCl_2_, and 5 mM DTT in the reaction mixture. The mixture was incubated at room temperature. The phosphorylation level of ERK2 was monitored by LC-MS. Once the protein was nearly completely phosphorylated at both Y187 and T185, the reaction mixture was loaded into a Source-15Q anion exchange column and eluted with a linear gradient from 100% buffer A to 40% buffer B to separate pTY-ERK2 from uERK2, pY-ERK2, pT-ERK2 and MEK-ΔN3-SESD. The phosphorylation state of ERK2 in each fraction was confirmed by LC-MS. The fractions corresponding to the pTY-ERK2 peak were pooled, supplemented with 5 mM DTT, and further purified using a Superdex 200 Increase 10/300 GL size-exclusion chromatography column equilibrated with SEC buffer. The peak fractions were pooled for biochemical assays.

### Preparation of thio-pTY-ERK1

The purified uERK1 was concentrated to 600 µM, and mixed with MEK-ΔN3-SESD in an ATPγS-containing buffer to achieve a final concentration of 140 µM ERK1, 7 µM MEK-ΔN3-SESD, 25 mM Tris–HCl pH 8.0, 1% glycerol, 150 mM NaCl, 2 mM ATPγS, 10 mM MgCl_2_, and 5 mM DTT in the reaction mixture. The mixture was incubated at room temperature. The phosphorylation level of ERK1 was monitored by LC-MS. Once the protein was nearly completely thiophosphorylated at both Y204 and T202, the reaction mixture was concentrated using a centrifugal filter and desalted using a HiTrap Desalting column equilibrated with SEC buffer to remove residual ATPγS or ADP. The desalted protein was then loaded into a Source-15Q anion exchange column and eluted with a linear gradient from 100% buffer A to 40% buffer B to separate thio-pTY-ERK1 from uERK1, thio-pY-ERK1, thio-pT-ERK1 and MEK-ΔN3-SESD. The phosphorylation state of ERK1 in each fraction was confirmed by LC-MS. The fractions corresponding to the thio-pTY-ERK1 peak were pooled, supplemented with 5 mM DTT, and further purified using a Superdex 200 Increase 10/300 GL size-exclusion chromatography column equilibrated with SEC buffer. The peak fractions were pooled for biochemical assays and cryo-EM sample preparation.

### MEK1 kinase activity assay

Purified C-terminal His-tagged phosphorylated MEK1 (pMEK1), C-terminal His-tagged unphosphorylated MEK1 (uMEK1) and other indicated mutants were diluted to 10 nM using a reaction buffer (25 mM HEPES pH 7.4, 150 mM NaCl, 10 mM MgCl_2_, 5 mM DTT). Similarly, purified C-terminal His-tagged ERK1/2 (uERK1/2) or the indicated mutants was diluted to 4 µM using the reaction buffer. ATP (Sigma-Aldrich, Cat# A6419) was prepared as 100 mM stock in aqueous solution, adjusted to pH 7.0 by NaOH. ATP stock was diluted to 10 mM using the reaction buffer.

For the kinase reaction, 80 µL of 4 µM uERK1/2 was mixed with 80 µL of 10 nM MEK1. The mixture was mixed with 160 µL of 10 mM ATP to initiate the reaction and incubated at 30 °C. The final concentrations of MEK1, ERK1/2 and ATP in the reaction mixture were 2.5 nM, 1 µM and 5 mM, respectively. At different time points, 10 µL of the mixture was immediately added into 90 µL of 2× SDS loading buffer and heated at 95 °C for 2 min to terminate the reaction. Then the samples were subjected to immunoblotting. The bands on the immunoblotting image were quantified using ImageJ (version 1.54g).

### ERK1 kinase activity assay

The ERK1 activity was measured using an ADP-Glo^TM^ Kinase Assay (Promega, Cat# V6930) following the manufacturer’s protocol. The ATP + ADP standards were prepared as described in the protocol. Purified uERK1, pY-ERK1, pTY-ERK1 and thio-pTY-ERK1 were individually diluted to concentrations ranging from 0 to 50 nM using buffer K (50 mM HEPES pH 7.4, 150 mM NaCl, 0.01% Triton X-100, 0.01% BSA, 10 mM MgCl_2_, 5 mM DTT). The MBP peptide FFKNIVTPRTPPPSQGK (5 mM, in H_2_O) was used as the substrate, and was diluted to 200 µM using buffer K. The ATP stock was also diluted to 20 µM using buffer K.

For the kinase reaction, 12 µL of diluted ERK1 was mixed with 12 µL of MBP-peptide solution, followed by the addition of 24 µL of diluted ATP solution. The reaction mixture was then incubated at room temperature for 40 minutes. After incubation, 10 µL of the reaction mixture was transferred to a 384-well plate. Next, 10 µL of ADP-Glo^TM^ Reagent was added to each well in order to deplete unconsumed ATP. The plate was then incubated at room temperature for 40 minutes. After that, 20 µL of Kinase Detection Reagent was added to each well to convert ADP back to ATP and to introduce luciferase and luciferin for ATP detection. The plate was incubated again at room temperature for 20 minutes. Finally, luminescence was measured using a Tecan Spark plate reader with an integration time of 800 ms per well. The concentration of ATP consumed by ERK1 was calculated based on a luminescence curve of ATP + ADP standards. This assay was performed with three independent measurements, each consisting of technical triplicates, and was statistically analyzed using GraphPad Prism 10.

### MEK1 ATPase activity assay

Proteins after size exclusive chromatography (SEC) were thawed at room temperature and put at 4 . All proteins were diluted to 400 nM, 200 nM, 100 nM, 50 nM, 25 nM and 12.5 nM in a 2-fold serial dilution. Dilution was conducted in the same buffer used in SEC (25 mM HEPES pH 7.4, 150 mM NaCl, 5 mM MgCl_2_). ATP stock was diluted to 100 μM by SEC buffer. Then 60 μL proteins dilution was mixed with 60 μL of 100 μM ATP, and incubated at 37 for 30 min. 50 μL of each sample was transferred to 96-well UV transparent plate (Corning, 3695) in duplicates. The free phosphate was detected by PiColorLock^TM^ phosphate assay kit (Abcam, ab270004). Calibration curve, control group, volume and incubation time of detection were set according to the product protocol. In brief, after hydrolysis, 12.5 µL of PiColorLock mix was added into each well and incubated for 5 mins before adding 5 µL of PiColorLock stabilizer. After adding stabilizer, mixture was allowed to incubate at room temperature for 30 mins before absorption detection. Absorption was detected at 650 nm, using a microplate reader (Spark, Tecan). The data was analyzed by using GraphPad Prism 10.

### MEK1 dephosphorylation activity assay

To assess the dephosphorylation activity of MEK1 under the condition used for preparing the cryo-EM samples (in the presence of 5 mM AMP-PNP), purified pMEK1, pY-ERK1, and pTY-ERK1 were each diluted to a 2× stock using a reaction buffer (25 mM HEPES pH 7.4, 150 mM NaCl, 10 mM MgCl_2_, 5 mM DTT) supplemented with 5 mM AMP-PNP. The reactions were initiated by mixing 10 µL of pY-ERK1 or pTY-ERK1 with 10 µL of either the reaction buffer or pMEK1, and then incubated on ice.

To assess the pMEK1-catalyzed transfer of the phosphate group from pY-ERK1 to GST-uERK1/2, purified pMEK1 was prepared as a 2× stock using the reaction buffer, and purified GST-uERK1/2 and pY-ERK1/2 were each prepared as a 4× stock using the same buffer. The reactions were initiated by mixing GST-uERK1/2 with pY-ERK1/2 or the same volume of the reaction buffer, and then mixed with pMEK1 at 30 °C.

For assays performed in the presence of apyrase (Sigma, A6535), proteins were prepared in a 2× stock in buffer containing 25 mM HEPES (pH 7.4), 150 mM NaCl, 5 mM MgCl_2_, 5 mM DTT, and 5 mM CaCl_2_. To deplete residual ATP and ADP, 20 µL of pMEK1 or ERK1 was firstly incubated with 1 µL apyrase (0.2 U) or apyrase buffer (25 mM HEPES pH 7.4 and 5 mM CaCl_2_) at 30 °C for 20 min. Reactions were initiated by mixing 15 µL of ERK1 with 15 µL of pMEK1 from the pre-incubation step, or with an equal volume of buffer (control), followed by incubation at 30 °C. The final concentrations of pMEK1 and ERK1 in the reaction mixture were 2 µM and 12 µM, respectively.

At each time point, 2–6 µL of the reaction mixture was removed and immediately diluted by 10 mM Tris–HCl pH 8.0, mixed with SDS loading buffer and heated at 95 °C for 2 min. Then the samples were subjected to immunoblotting and quantified with ImageJ (version 1.54g).

### Dephosphorylation of pTY–ERK1/2 at varying nucleotide ratios

Purified pMEK1 was prepared as a 2× stock in a reaction buffer containing 25 mM HEPES (pH 7.4), 150 mM NaCl, 10 mM MgCl_2_, and 5 mM DTT. pTY-ERK1/2 were each prepared as 4× stocks in the same buffer. Nucleotide mixtures with varying ATP/ADP ratios (0–100% ATP) were prepared by combining appropriate volumes of ATP and ADP stock solutions to a final total nucleotide concentration of 20 mM, and prepared as 4× stocks in the same buffer. Reactions were initiated by first mixing pTY-ERK1/2 with the nucleotide mixtures, followed by addition of pMEK1, and incubated at 30 °C. The final concentrations of pMEK1, pTY-ERK1/2, and total nucleotides were 200 nM, 1 µM, and 5 mM, respectively.

For assays including DUSP6, purified pMEK1 and DUSP6 were each prepared as 4× stocks in the reaction buffer, and pTY-ERK1 was prepared as a 4× stock. Nucleotide mixtures with varying ATP/ADP or ATP/AMP-PNP ratios (0–100% ATP; total nucleotide concentration of 20 mM) were prepared as described above. Reactions were initiated by first mixing pTY-ERK1 with the nucleotide mixtures, followed by addition of pre-mixed pMEK1–DUSP6, and incubated at 30 °C. The final concentrations of pMEK1, DUSP6, pTY-ERK1/2, and total nucleotides were 200 nM, 200 nM, 1 µM, and 5 mM, respectively.

After 30 min, 20 µL of each reaction mixture was removed and quenched with 1 % (*v*/*v*) formic acid and then analyzed by LC-MS.

### [^32^P]pY-ERK1 preparation

Autophosphorylation reactions were performed at 37 °C for 1 h in a total volume of 120 µL for four groups: ERK1, ERK1(T202V), ERK1(Y204F), and ERK1(T202V/Y204F). Reaction mixtures contained ERK1 or the indicated ERK1 mutants (48 µM), ATP (1 mM), and [γ-^32^P]ATP (41.25 nM; typically, 15 µCi per reaction; PerkinElmer, BLU502A250UC) in buffer containing 25 mM HEPES (pH 7.4), 150 mM NaCl, 5 mM MgCl_2_, and 5 mM DTT.

Following incubation, samples were centrifuged at 22,000 × g for 3 min. The supernatant (120 µL) was subjected to desalting using Zeba Spin Desalting Columns (7K MWCO; Thermo Scientific, 89882) to remove excess ATP. To further eliminate residual [γ-^32^P]ATP, samples were supplemented with ATP to a final concentration of 1 mM after the first round of desalting purification and incubated for 10 min at room temperature, followed by four additional rounds of desalting (five rounds total).

To assess residual free [γ-^32^P]ATP, 5 µL of desalted protein sample was mixed with either 5 µL of 100% acetonitrile to precipitate proteins or 5 µL of the reaction buffer (control). After centrifugation, the supernatant was collected, mixed with 100 µL of scintillation cocktail, and radioactivity was measured using a MicroBeta2 scintillation counter (PerkinElmer) to obtain CPM values. The measured CPM values (protein vs. protein-precipitated supernatant) were as follows: [^32^P]pY-ERK1, 9071 vs. 264; [^32^P]pY-ERK1(T202V), 10,190 vs. 300; ERK1(Y204F), 805 vs. 364; ERK1(T202V/Y204F), 377 vs. 206. Protein concentrations were determined by Bradford assay using ERK1 proteins as standards, and the final concentrations of [^32^P]pY-ERK1 and the indicated mutants were approximately 24 µM.

### Radioactive assay

To assess pMEK1-catalyzed phosphate transfer from ^32^P-labeled ERK1 ([^32^P]pY-ERK1) to GST-ERK1, purified [^32^P]pY-ERK1 or the indicated ERK1 mutants (after desalting) were prepared as 2× stocks. Purified GST-ERK1 (or corresponding mutants) and pMEK1 or pMEK1(D208A) were each prepared as 4× stocks in the same buffer. Reactions were initiated by first mixing GST-ERK1 with [^32^P]pY-ERK1, followed by addition of pMEK1, and incubated at 30 °C. The final concentrations of pMEK1, [^32^P]pY-ERK1 and GST-ERK1 in the reaction mixture were 2 µM, 12 µM and 12 µM, respectively. For assays performed in the presence of ADP (Selleck, Cat# S6325, 100 mM stock in H_2_O), reactions were carried out under identical conditions with a final concentration of 1 mM ADP.

After 5 min, 18 µL of the reaction mixture was mixed with 18 µL of 2× SDS loading buffer and heated at 95 °C for 2 min to terminate the reaction. 20 µL of each sample was resolved by SDS–PAGE, and the gel was exposed to phosphor screens (BAS-SR 2025, Fujifilm) for 4 h prior to imaging using a phosphor imager (Amersham Typhoon IP, Cytiva). An additional 8 µL of each sample was analyzed by SDS–PAGE followed by Coomassie Brilliant Blue staining.

In parallel, 5 µL of the reaction mixture was mixed with 5 µL of 100% acetonitrile to precipitate proteins. Following centrifugation, the supernatant was collected for thin-layer chromatography (TLC) analysis. Equal amounts of [^32^P]pY-ERK1 (after desalting), as well as 1:10 and 1:100 dilutions and [γ-^32^P]ATP, were included as standards. TLC was performed largely as previously described, with minor modifications^32^. Briefly, 1 µL aliquots of reaction mixtures or indicated standards were spotted twice (2 µL total) onto 20 × 20 cm PEI cellulose F TLC plates (Sigma, 1055790001). Plates were developed for 40 min at room temperature in a solvent system containing 0.5 M LiCl and 1.5 M formic acid. After drying, phosphor screens were exposed for 14 h prior to imaging using a phosphor imager. ATP and ADP standards were visualized under UV light at 254 nm and marked with pencil. Free inorganic phosphate (Pi) standards were detected by spraying the plates with a solution containing 1% (*w*/*v*) ammonium molybdate, 3 % (*v*/*v*) H_2_SO_4_, and 5.2% (*w*/*v*) sodium ascorbate.

Quantification was performed using the original phosphor images in ImageJ and analyzed by using GraphPad Prism 10. Phosphor images, pencil-marked photographs, and spray-stained images were overlaid using Adobe Photoshop for comparison.

### Immunoblotting

10 µL of each sample was subjected to 4-20% SDS–PAGE (GenScript, Cat# M00657) and transferred to 0.2 µm pore size PVDF membrane (Merck Millipore). The membrane was blocked with 5% non-fat milk (Beyotime, Cat# P0216-1500g) in TBST buffer (25 mM Tris–HCl 7.4, 140 mM NaCl, 30 mM KCl, 0.1% Tween 20) for 1 hour at room temperature with shaking, and then cut horizontally according to the molecular weight of target proteins, followed by incubation with primary antibodies for overnight at 4 °C with shaking. The membrane was then washed 5 times with TBST buffer, incubated with appropriate HRP-conjugated secondary antibodies (Merck Millipore, Cat# AP127P or Cat# AP156P) for 1 hour at room temperature, and washed 5 times with TBST buffer. The secondary antibodies on the PVDF membrane were visualized by adding ECL western blot reagents (FDbio, Cat# FD8020), followed by detection of the luminance signal using Amersham Imager 680. The bands on the immunoblotting image were quantified using ImageJ (version 1.54g).

Primary antibodies used in this study include Anti-MEK1/2 (CST, Cat# 4694S), Anti-phospho-MEK1/2 (Ser217/221) (CST, Cat# 9154S), Anti-MAPK (ERK1/2) (CST, Cat# 4695S), Anti-phospho-MAPK (ERK1/2) (Thr202) (CST, Cat# 4370S) and Anti-phospho-MAPK (ERK1/2) (Tyr204) (CST, Cat# 4377S). The 4370S antibody detects mono-phosphorylated ERK1 (at Thr202) and dually phosphorylated ERK1 (at Thr202 and Tyr204). The 4377S antibody detects mono-phosphorylated ERK1 (at Tyr204) and dually phosphorylated ERK1 (at Thr202 and Tyr204).

### Crosslinking assay

For disulfide crosslinking experiments, purified pMEK1 (WT or N78C) and ERK1 (WT or T202C) proteins in SEC buffer containing 25 mM HEPES (pH 7.4), 150 mM NaCl, and 5 mM MgCl_2_ were mixed at a final concentration of 10 µM each in the presence of 1 mM AMP-PNP. Oxidative crosslinking was initiated by addition of GSSG (BD02544320, 10 mM stock prepared in SEC buffer) to a final concentration of 0.4 mM, and reactions were incubated at room temperature for 2 h. Following incubation, samples were mixed with 2× SDS loading buffer under reducing (200 mM DTT) or non-reducing conditions and analyzed by SDS–PAGE to assess disulfide-dependent complex formation.

### Liquid chromatography–mass spectrometry (LC-MS)

Chromatographic elution was performed on an UPLC system (Waters, USA) equipped with Time-of-Flight Mass Spectrometry (Waters Xevo G2 XS, USA) with an electrospray ion (ESI) source in positive ion modes. The separation was carried out on the BEH C4 column (50 mm × 2.1 mm, 1.7 µm) at 40 °C. The mobile phase A was ultra-pure water containing 0.1% formic acid, and the mobile phase B was acetonitrile containing 0.1% formic acid. The flow rate was 0.3 mL/min, and the gradient of mobile phase B was held at 5% for 1 min, then 5% to 95% in 4 min, and finally held at 95% for 2 min. The sample volume injected was 1 µL (0.5 µM). A Waters TOF-MS system in the ESI+ modes was used for the detection. The parameters used were as follows: Capillary, 3 kV; Sample Cone, 50 V; Source Offest, 30 V; Source temperature, 140 °C; Desolvation temperature 600 °C. nebulizer gas, 35 psi; Cone gas flow rate, 50 L/h; and Desolvation gas flow rate 800 L/h. A full scan was run with a mass range from *m*/*z* 200 to 3000. Masslynx was used for data acquisition and data analysis.

### Gel filtration assay

The binding of uMEK1 or pMEK1 to GST-ERK1 in the presence of ADP was analyzed using gel filtration. Purified uMEK1 or pMEK1 was mixed with purified GST-ERK1 and ADP in SEC buffer to a final volume of 400 µL. The final concentrations of MEK1, ERK1, and ADP in the mixture were 2 µM, 10 µM, and 5 mM, respectively. Control groups were prepared by replacing either MEK1 or GST-ERK1 stock with an equivalent volume of SEC buffer. The mixtures were incubated at 4 °C for 2 hours before gel filtration. After incubation, each mixture was loaded into a Superdex 200 Increase 10/300 GL column equilibrated with a buffer containing 25 mM HEPES pH 7.4, 150 mM NaCl, 1 mM DTT, 5 mM MgCl_2_, and 5 mM ADP. The eluted fractions were analyzed by SDS–PAGE.

### Isothermal Titration Calorimetry (ITC)

ITC experiments were done with the isothermal titration calorimeter MicroCal PEAQ-ITC (Malvern Panalytical). In these experiments, a 20 µM solution of either pMEK1 or uMEK1 in the gel filtration buffer (25 mM HEPES pH 7.4, 150 mM NaCl, 5 mM MgCl_2_) was titrated with a 200 µM solution of either uERK1 or thio-pTY-ERK1 in the same buffer. The titrations were performed in the presence or absence of 1 mM AMP-PNP (Sigma, Cat# A2647, 100 mM stock in H_2_O) or ADP.

For assays performed in the presence of apyrase (Sigma, A6535), a 20 µM solution of pMEK1 in buffer (25 mM HEPES pH 7.4, 150 mM NaCl, 5 mM MgCl_2_, and 5 mM CaCl_2_) was titrated with a 200 µM solution of uERK1, pY-ERK or pTY-ERK1 in the same buffer. Titrations were carried out in the presence of 1 mM AMP-PNP and 0.004 U/µL apyrase. The phosphorylation state of ERK1 after titration was validated by mass spectrometry.

All experiments were conducted at a constant temperature of 25 °C. The resulting ITC data were analyzed using the MicroCal PEAQ-ITC analysis software to determine the binding affinity.

### Cryo-EM sample preparation and data acquisition

To prepare cryo-EM samples of the AMP-PNP-bound pMEK1:uERK1/2 complex, purified pMEK1 and uERK1/2 were mixed in a buffer containing 25 mM HEPES pH 7.4, 150 mM NaCl, 10 mM MgCl_2_ and 5 mM DTT. AMP-PNP was added to the mixture to achieve a final concentration of 5 mM. The mixture was then incubated on ice for 2 hours.

To prepare the cryo-EM sample of the pMEK1:uERK1 complex (derived from AMP-PNP-bound pMEK1:pY-ERK1 complex) or the pMEK1:pT-ERK1 complex (derived from AMP-PNP-bound pMEK1:pTY-ERK1 complex), purified pMEK1 were mixed with either pY-ERK1 or pTY-ERK1 in the reaction buffer. The final concentrations of pMEK1, pY-ERK1 and pTY-ERK1 were all 50 µM. AMP-PNP (100 mM stock in H2O) was added to the mixture to achieve a final concentration of 5 mM. The mixture was then incubated on ice for 2 hours. The phosphorylation status of ERK1 in the samples was assessed using immunoblotting and LC-MS.

To prepare the cryo-EM sample of the AMP-PNP-bound pMEK1:pY-ERK1 complex, 10 µL of pMEK1 (100 µM in the reaction buffer) was mixed with 10 µL of pY-ERK1 (100 µM in the reaction buffer) in the presence of 2 mM AMP-PNP, and then the mixture was immediately loaded to glow discharged grids.

To prepare the cryo-EM sample of the ADP-AlF_3_-bound pMEK1:uERK1 complex, pMEK1 was first equilibrated in buffer containing 25 mM HEPES (pH 7.4), 150 mM NaCl, 5 mM MgCl_2_, 2 mM DTT, 1 mM ADP, 50 mM NaF, and 1 mM AlCl_3_. uERK1, prepared in the same buffer, was then mixed with pMEK1. The final concentrations of both pMEK1 and uERK1 were 50 µM. The mixture was incubated on ice for 10 min.

To prepare the cryo-EM sample of the crosslinked pMEK1(N78C):uERK1(T202C) complex, purified pMEK1(N78C) was incubated with ERK1(T202C) in a buffer containing 25 mM HEPES (pH 7.4), 150 mM NaCl, 5 mM MgCl_2_, and 1 mM AMP-PNP. Disulfide crosslinking was induced by addition of GSSG to a final concentration of 0.4 mM. The mixture was incubated at room temperature for 3 h to allow formation of the disulfide bond-linked pMEK1–ERK1 complex.

To prepare cryo-EM sample grids, the grids (holey carbon film on 300 mesh Au, R 1.2/1.3, Quantifoil) were glow discharged with a medium RF power for 30 s by a plasma cleaner (PDC-32G-2, Harrick Plasma) after vacuuming for 1 min. After pretreatment, 4 μL of protein sample was loaded to a glow discharged grid and blotted for 3.5 s with zero force immediately at 100% humidity and 8 °C chamber temperature, then flash frozen in liquid ethane using Vitrobot (Mark IV, Thermo Fisher Scientific). The cryo-EM grids were stored in liquid nitrogen before being used for data collection.

For all samples except the crosslinked pMEK1(N78C):uERK1(T202C) complex, after quality evaluation with a Glacios TEM (FEI), the grids were transferred to a Titan Krios TEM (FEI) operating at 300 kV equipped with a K3 Summit direct electron detector (Gatan) mounted on a GIF Quantum energy filter (Gatan). Zero-loss movie stacks were automatically collected using EPU software (FEI) with a slit width of 20 eV on the energy filter and a defocus range from -1.1 µm to -1.8 µm in super-resolution mode at a nominal magnification of 81,000 ×. Each stack, which contained 32 frames, was exposed for 2.56 s with a total electron dose of ∼ 50 e-/Å^2^. The movie stacks were imported to cryoSPARC v4^33^ (Structura Biotechnology) for downstream processing. For the crosslinked pMEK1(N78C):uERK1(T202C) complex, after quality evaluation with a Glacios TEM (FEI), the grid was transferred to a Titan Krios TEM (FEI) operating at 300 kV equipped with a Falcon4i Summit direct electron detector mounted on a Selectris X energy filter. Zero-loss movie stacks were automatically collected using EPU software (FEI) with a slit width of 10 eV on the energy filter and a defocus range from -1 µm to - 1.9 µm in super-resolution mode at a nominal magnification of 215,000×. Each stack was exposed for 1.99 s with a total electron dose of ∼ 50 e-/Å^2^. The movie stacks were imported to cryoSPARC v4^33^ (Structura Biotechnology) for downstream processing.

### Structure determination and model building

The flowcharts of cryo-EM data processing are illustrated in Figures S2 and S5. All the steps were performed in cryoSPARC v4^33^ unless otherwise stated.

For the data processing of the pMEK1:pT-ERK1 complex (derived from pMEK1:pTY-ERK1 complex) in dataset D, a total of 4,198 movie stacks, with a pixel size of 1.0773 Å, were motion-corrected using MotionCor2^34^ with dose-weighting, followed by patch-based contrast transfer function estimation (Patch CTF Estimation). Initial particle picking yielded 4,226,273 particles, with a box size of 192 pixels cropped to 128 pixels. After 2D classification, 4,170,611 particles were selected for ab-initio reconstruction. One dominant structural class, containing 2,525,371 particles, underwent further 2D classification and ab-initio reconstruction. Of those, 742,809 particles were subjected to Topaz training. The Topaz picking model was then used for particle extraction from the 4,198 micrographs, resulting in 5,259,866 particles with a box size of 192 pixels cropped to 128 pixels for downstream processing. For ab-initio reconstruction, a good class containing 227,714 particles was further refined by non-uniform refinement^35^, resulting in a 3.56 A map (Fourier shell correlation (FSC) 0.143). With this map as a good reference and several maps from ab-initio reconstruction as bad references, heterogeneous refinement was performed to obtain more high-quality particles from the 2,525,371 particles. The target particles were re-extracted with a box size of 192 pixels for non-uniform refinement, resulting in a 3.23 A map (FSC 0.143) using 523,917 particles. To further improve the map quality, multiple rounds of 2D classification and heterogeneous refinement were performed after Topaz extraction. Subsequently, non-uniform refinement was performed with 555,122 selected particles after re-extraction with a box size of 192 pixels, resulting in a final map at 3.12 A resolution (FSC 0.143). All the map figures were generated using ChimeraX^36^.

For the data processing of the pMEK1:uERK1 complex (derived from pMEK1:pY-ERK1 complex) in dataset B, the pixel size of the movie stacks in dataset B was 1.087 A, differing from that in dataset D. To get a high-quality map based on the well-characterized particles and the final map from dataset D, 5,618 dose-weighted micrographs with a pixel size of 1.0773 Å were utilized for patch CTF estimation. These micrographs were subsequently combined with those from dataset D for particle picking and extraction, yielding 25,107,479 particles with a box size of 192 pixels cropped to 32 pixels. After three rounds of 2D classification, 17,001,789 particles were selected for heterogeneous refinement and ab-initio reconstruction, yielding a good class of 8,244,378 particles with a box size of 192 pixels cropped to 128 pixels after re-extraction. Two additional rounds of 2D classification were then performed, followed by heterogeneous refinement. The particles of the target class were reassigned to micrographs from dataset B, with a pixel size of 1.087 Å, and re-extracted with a box size of 192 pixels for non-uniform refinement. This process yielded a final map at 3.20 Å resolution (FSC 0.143) using 235,273 particles.

The data processing of the pMEK1:uERK1 complex in dataset A and pMEK1:pY-ERK1 complex in dataset C were similar. Briefly, 5,966 and 6,617 micrographs in dataset A and C, respectively, with a pixel size of 1.0773 Å, were utilized for particle picking and extraction, yielding 24,513,458 and 11,179,582 particles with a box size of 192 pixels cropped to 32 pixels, respectively. After 2D classification, the selected particles in dataset A were re-extracted with a box size of 192 pixels cropped to 128 pixels for further 2D classification, while the selected particles in dataset C were re-extracted with a box size of 192 pixels cropped to 48 pixels. Subsequently, 8,322,704 and 9,585,072 particles in datasets A and C, respectively, were selected for several rounds of heterogeneous refinement. The particles of the target class were extracted with a box size of 192 pixels for non-uniform refinement, resulting in final maps at 3.36 Å resolution (FSC 0.143) using 319,939 particles for dataset A and 3.36 Å resolution (FSC 0.143) using 216,414 particles for dataset C.

For the data processing of the pMEK1:uERK2 complex, a total of 7,970 micrographs with a pixel size of 1.0773 Å were used for particle picking and extraction, yielding 12,045,667 particles. The initial extraction box size was 192 pixels, which was subsequently cropped to 128 pixels. After three rounds of 2D classification, 3,503,882 particles were re-extracted using a box size of 192 pixels cropped to 48 pixels for ab-initio reconstruction under different conditions. The good classes were then extracted with a box size of 192 pixels and subjected to non-uniform refinement. After removal of duplicate particles, 1,729,222 particles were subjected to heterogeneous refinement, followed by non-uniform refinement. The final map, calculated from 609,769 particles, reached a resolution of 2.93 Å (FSC 0.143).

For the data processing of the pMEK1:uERK1 complex with ADP and AlF_3_, a total of 8,032 micrographs with a pixel size of 1.0773 Å were used for particle picking and extraction, yielding 11,784,916 particles. The initial extraction box size was 192 pixels, which was subsequently cropped to 128 pixels. After three rounds of 2D classification, 3,269,105 particles were re-extracted using a box size of 192 pixels cropped to 48 pixels for ab-initio reconstruction. One good class containing 879,372 particles was then extracted with a box size of 192 pixels and subjected to non-uniform refinement, resulting in a 3.21 Å resolution map (FSC 0.143). Simulated particles derived from this map were combined with the initial particle set for another round of 2D classification. From these, 1,788,389 particles were used in parallel for two independent approaches: ab-initio reconstruction, and heterogeneous refinement using the 3.21 Å map as a good reference and several ab-initio maps from the first reconstruction as bad references. The best particle classes obtained from both approaches were combined, yielding a target class of 1,054,683 particles. This combined particle set was then further processed through an additional round of heterogeneous refinement, followed by ab-initio reconstruction and non-uniform refinement. The final map, calculated from 403,726 particles, reached a resolution of 2.96 Å (FSC 0.143).

For the data processing of the pMEK1(N78C):uERK1(T202C) complex, a total of 25,750 micrographs with a pixel size of 0.57 Å were used for particle picking and extraction, yielding 52,692,322 particles. The initial extraction box size was 384 pixels, which was subsequently cropped to 96 pixels. After three rounds of 2D classification, 2,619,359 particles were re-extracted with a box size of 384 pixels cropped to 48 pixels for ab-initio reconstruction under different conditions. The good classes were then extracted with a box size of 384 pixels and subjected to non-uniform refinement, yielding maps with resolutions ranging from 2.67 to 2.88 Å (FSC 0.143). In parallel, duplicate particles were removed, and 1,341,851 particles from these maps were subjected to heterogeneous refinement using the best map as a good reference and several ab-initio volumes as bad references. Subsequent non-uniform refinement of 591,862 particles yielded a 2.55 Å resolution map (FSC 0.143). In an alternative approach, simulated particles derived from these maps were combined with the initial particle set for another round of 2D classification, from which 2,556,020 particles were subjected to heterogeneous refinement using the 2.55 Å map as a good reference, followed by non-uniform refinement. The final map, calculated from 1,082,206 particles, reached a resolution of 2.33 Å (FSC 0.143), and was subsequently flipped in ChimeraX^36^ for model building and further structural analysis.

The models of human MEK1:ERK1/2 complexes were manually built using Coot^37^. AlphaFold3 prediction^38^ of the human pMEK1:uERK1 complex was manually fitted into the density map to serve as the starting model. The model was then manually adjusted, using aromatic residues as landmarks due to their clear visibility in the cryo-EM map. Each residue was carefully inspected, with their chemical properties considered during modelling. The model was subsequently refined in Phenix^39^, employing with secondary structure and geometry restraints to prevent overfitting. Statistics related to data collection, 3D reconstruction, and model refinement can be found in Table S1. All figures in this article related to MEK1:ERK1/2 structures were generated using PyMOL^40^, Chimera^41^, and ChimeraX^36^.

