## Supplementary material for "Mechanistic insight into the dual phosphorylation of ERK1 and ERK2 by MEK1": Figures S1-S23 and Table S1

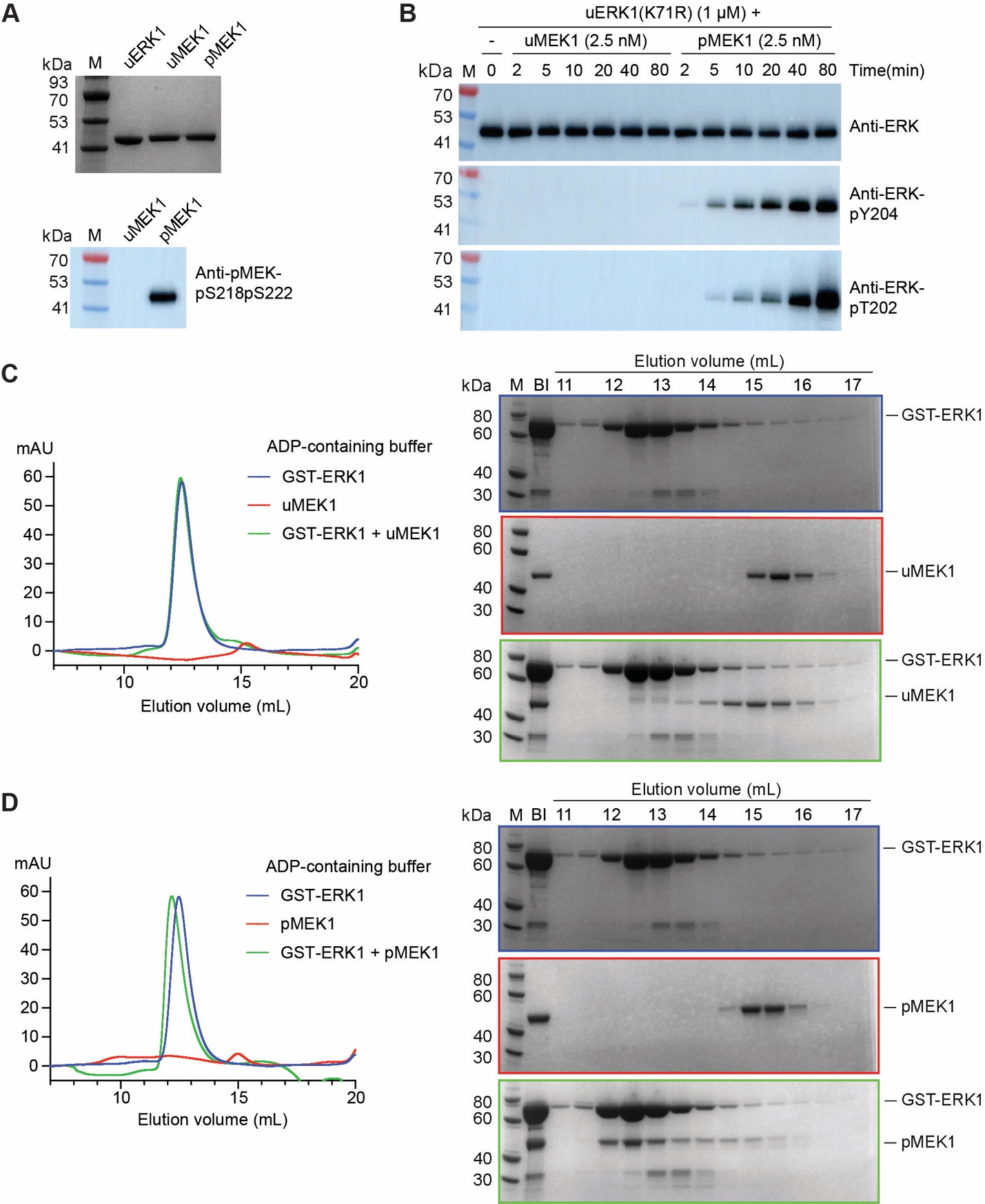
 Figure S1. Characterization of purified human MEK1 and ERK1

(A) Purified unphosphorylated ERK1 (uERK1) and MEK1 (uMEK1) and phosphorylated MEK1 (pMEK1) were examined by SDS–PAGE and visualized by Coomassie blue staining (upper panel). The phosphorylation states of uMEK1 and pMEK1 were confirmed by Western blotting (lower panel).

(B) The kinase activity of uMEK1 and pMEK1 was evaluated using a catalytic dead mutant of ERK1 (K71R) as the substrate protein. Phosphorylation of ERK1(K71R) was examined by Western blotting.

(C) uMEK1 cannot form a stable complex with ERK1. uMEK1 alone, GST tagged unphosphorylated ERK1 (GST-ERK1) alone, or a mixture of uMEK1 and GST-ERK1, was analyzed by gel filtration (left panel) and examined by SDS–PAGE and visualized by Coomassie blue staining (right panel). BI: before injection.

(D) pMEK1 can form a stable complex with ERK1. pMEK1 alone, GST-ERK1 alone, or a mixture of pMEK1 and GST-ERK1, was analyzed by gel filtration (left panel) and examined by SDS–PAGE and visualized by Coomassie blue staining (right panel).

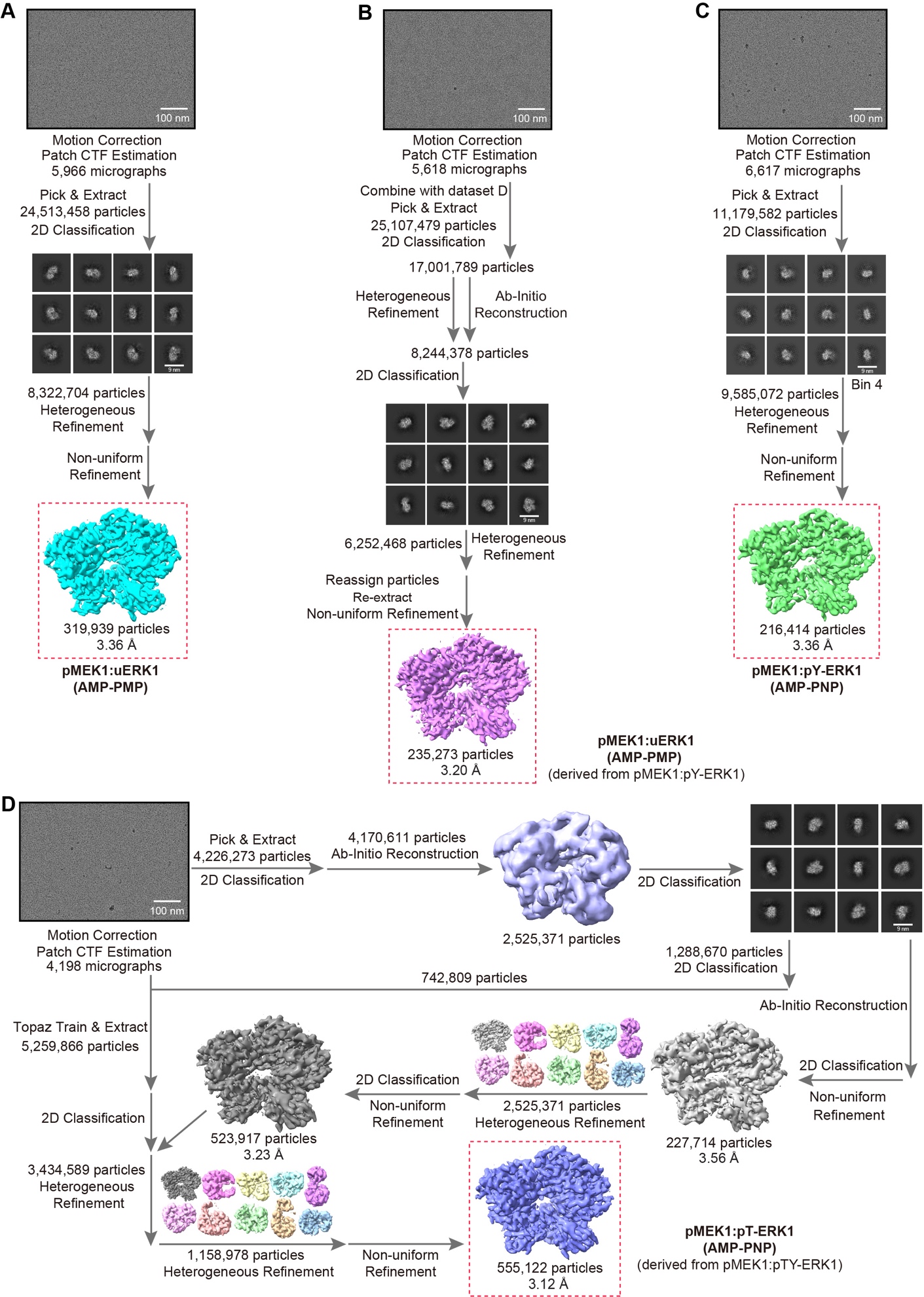
 Figure S2. Workflow of the EM data processing of the human MEK1:ERK1 complexes

(A) pMEK1:uERK1 complex.

(B) pMEK1:uERK1 complex, derived from the pMEK1:pY-ERK1 sample.

(C) pMEK1:pY-ERK1 complex.

(D) pMEK1:pT-ERK1 complex, derived from the pMEK1:pTY-ERK1 sample.

Detailed descriptions of the workflow are available in the ‘Materials and Methods’ section.

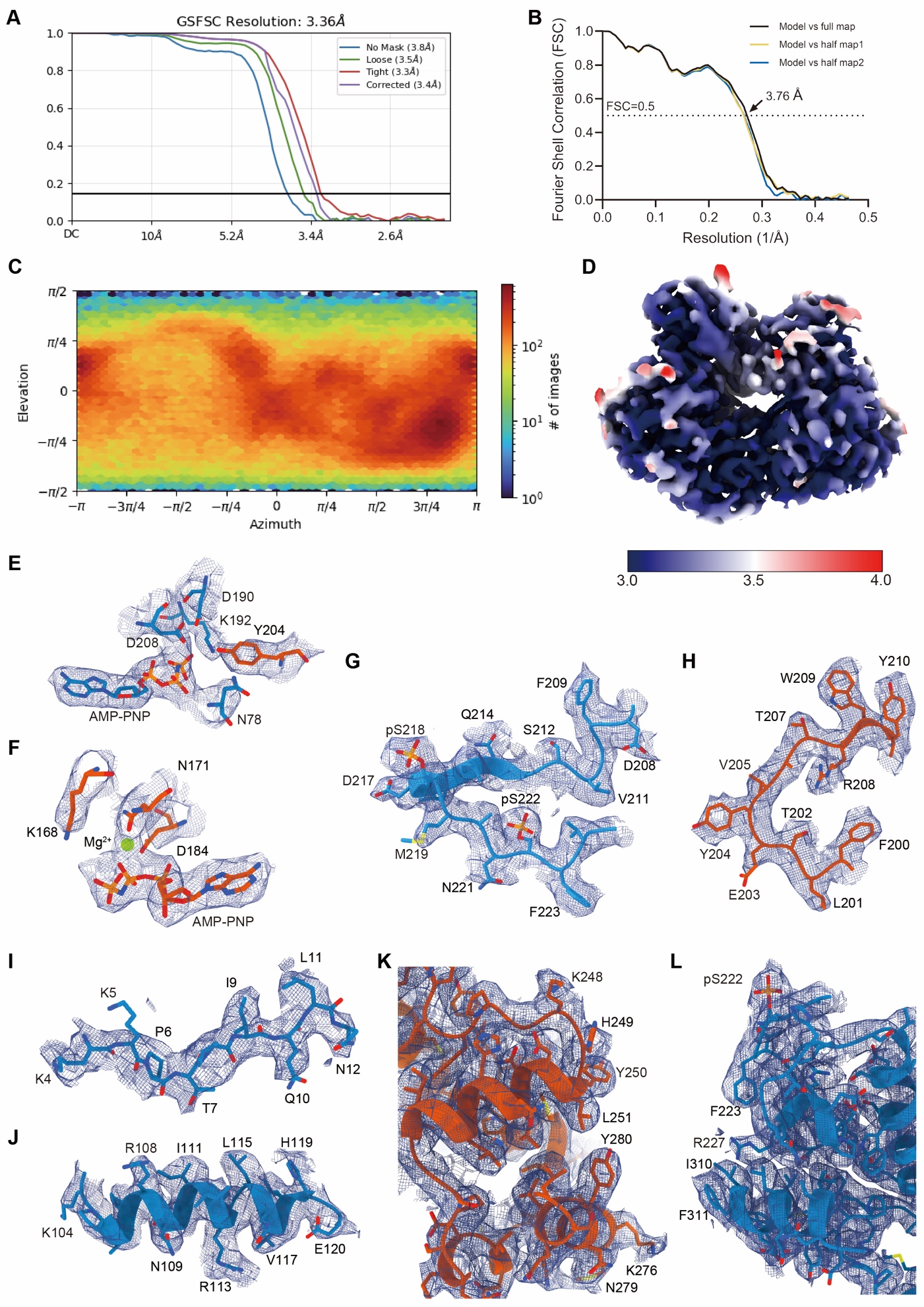

Figure S3. Cryo-EM analysis of the AMP-PNP-bound human pMEK1:uERK1 complex

(A) FSC curve for the final reconstruction of the pMEK1:uERK1 complex generated by CryoSPARC. Four curves correspond to the measurements with no mask (3.9 Å), loose mask (3.5 Å), tight mask (3.3 Å) and corrected mask (3.4 Å).

(B) FSC curves of the final refined model versus the summed map that it was refined against (black); of the model versus the first half-map (yellow); and of the model versus the second half-map (blue). The small difference between the yellow and blue curves indicates that the refinement of the atomic coordinates did not suffer from overfitting.

(C) Euler-angle distributions of the particles used for the final reconstruction.

(D) Local resolution of the reconstructed map.

(E–L) Representative cryo-EM maps of MEK1 (blue) and ERK1 (orange), at a contour level of 0.66. (E, F) ATP binding pockets. (G, H) activation loops. (I) N-terminal ERK docking site on MEK1. (J) αC helix of MEK1. (K, L) C-lobes near interface. The cryo-EM maps are shown as blue meshes.

The structural figures were made using UCSF ChimeraX (1.9).

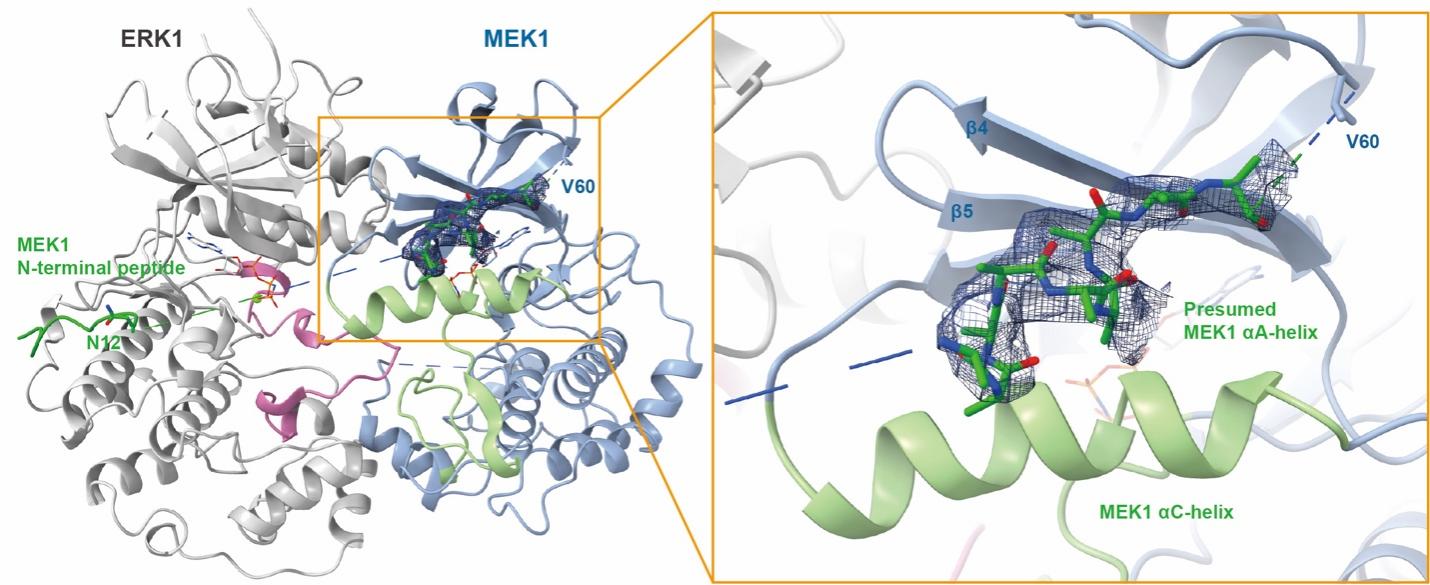

Figure S4. Structure of the αA-helix of MEK1 in the pMEK1:uERK1 complex

The cryo-EM map of the αA-helix of MEK1 was contoured at 0.66 and shown as a blue mesh. Only a short segment of the αA-helix is visible in the cryo-EM map, and for this short segment, only its main chain is resolved. This figure was made using UCSF ChimeraX (1.9).

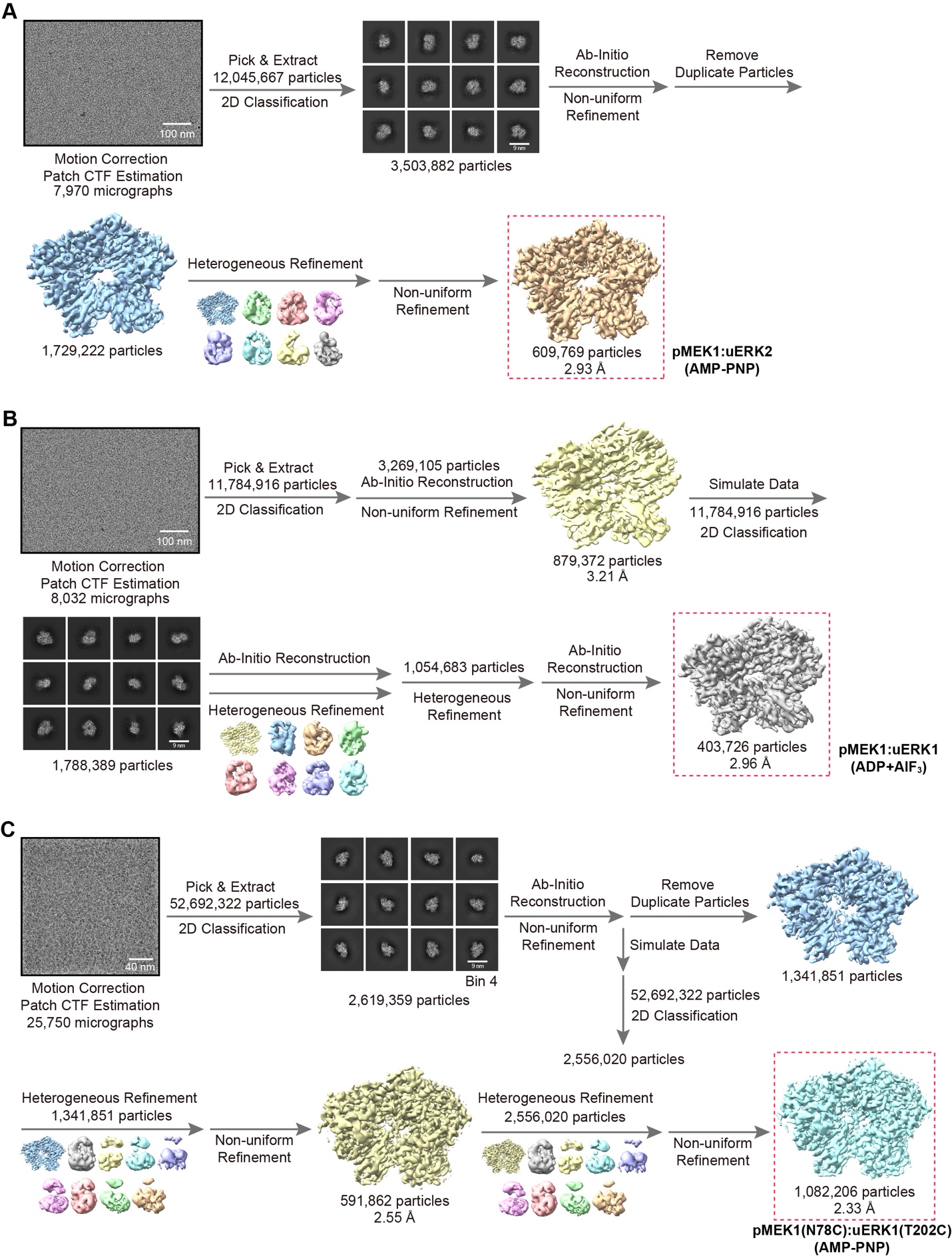

Figure S5. Workflow of the EM data processing of the human MEK1:ERK2 and MEK1:ERK1 complexes

(A) pMEK1:uERK2 complex in the presence of AMP-PNP.

(B) pMEK1:uERK1 complex in the presence of ADP and AlF_3_.

(C) pMEK1(N78C):uERK1(T202C) crosslinked complex in the presence of AMP-PNP.

Detailed descriptions of the workflow are available in the ‘Materials and Methods’ section.

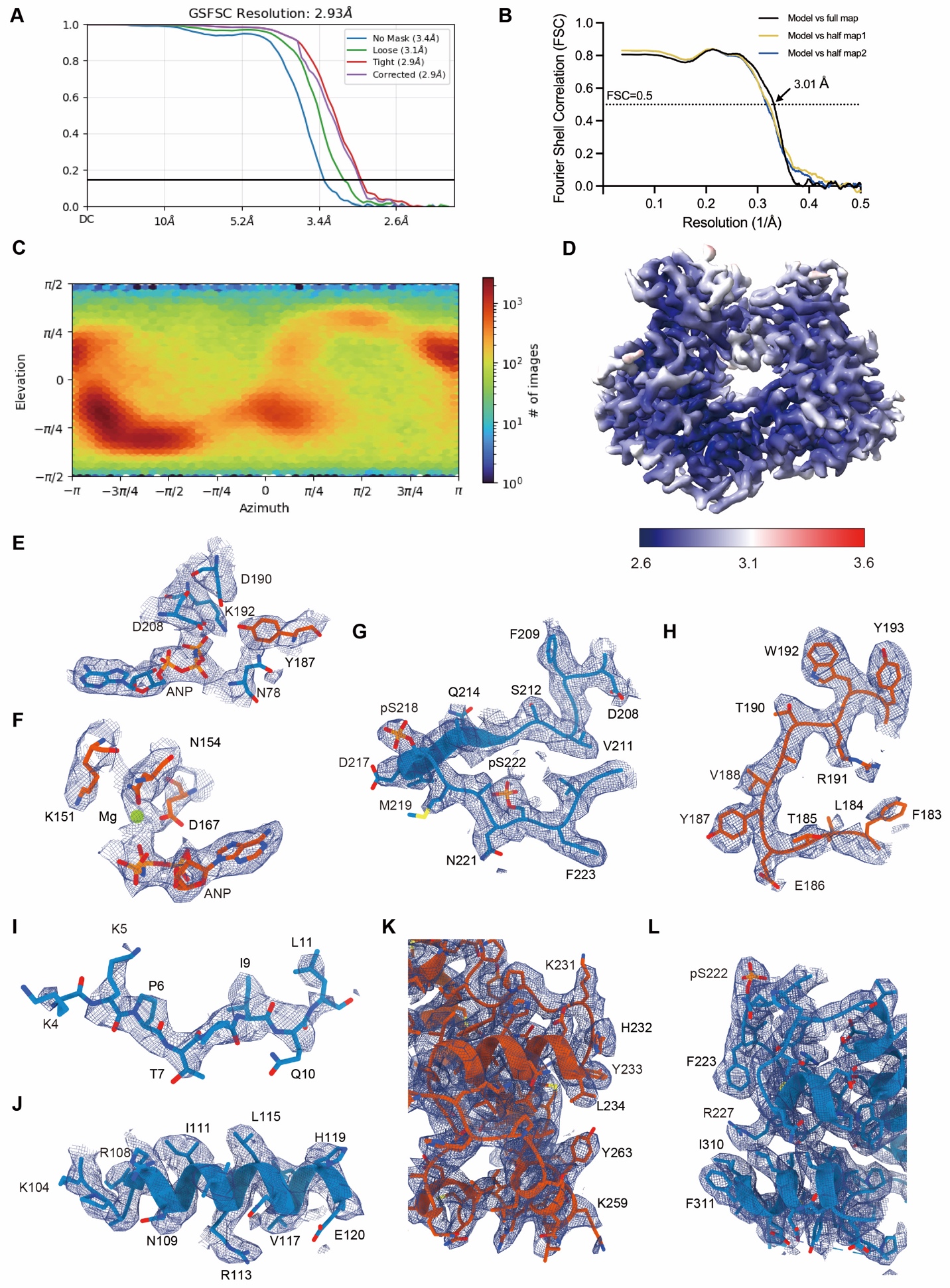

Figure S6. Cryo-EM analysis of the AMP-PNP-bound human pMEK1:uERK2 complex

(A) FSC curve for the final reconstruction of the pMEK1:uERK2 complex generated by CryoSPARC. Four curves correspond to the measurements with no mask (3.4 Å), loose mask (3.1 Å), tight mask (2.9 Å) and corrected mask (2.9Å).

(B) FSC curves of the final refined model versus the summed map that it was refined against (black); of the model versus the first half-map (yellow); and of the model versus the second half-map (blue). The small difference between the yellow and blue curves indicates that the refinement of the atomic coordinates did not suffer from overfitting.

(C) Euler-angle distributions of the particles used for the final reconstruction.

(D) Local resolution of the reconstructed map.

(E–L) Representative cryo-EM maps of MEK1 (blue) and ERK2 (orange), at a contour level of 0.41. (E, F) ATP binding pockets. (G, H) activation loops. (I) N-terminal ERK docking site on MEK1. (J) αC helix of MEK1. (K, L) C-lobes near interface. The cryo-EM maps are shown as blue meshes.

The structural figures were made using UCSF ChimeraX (1.9).

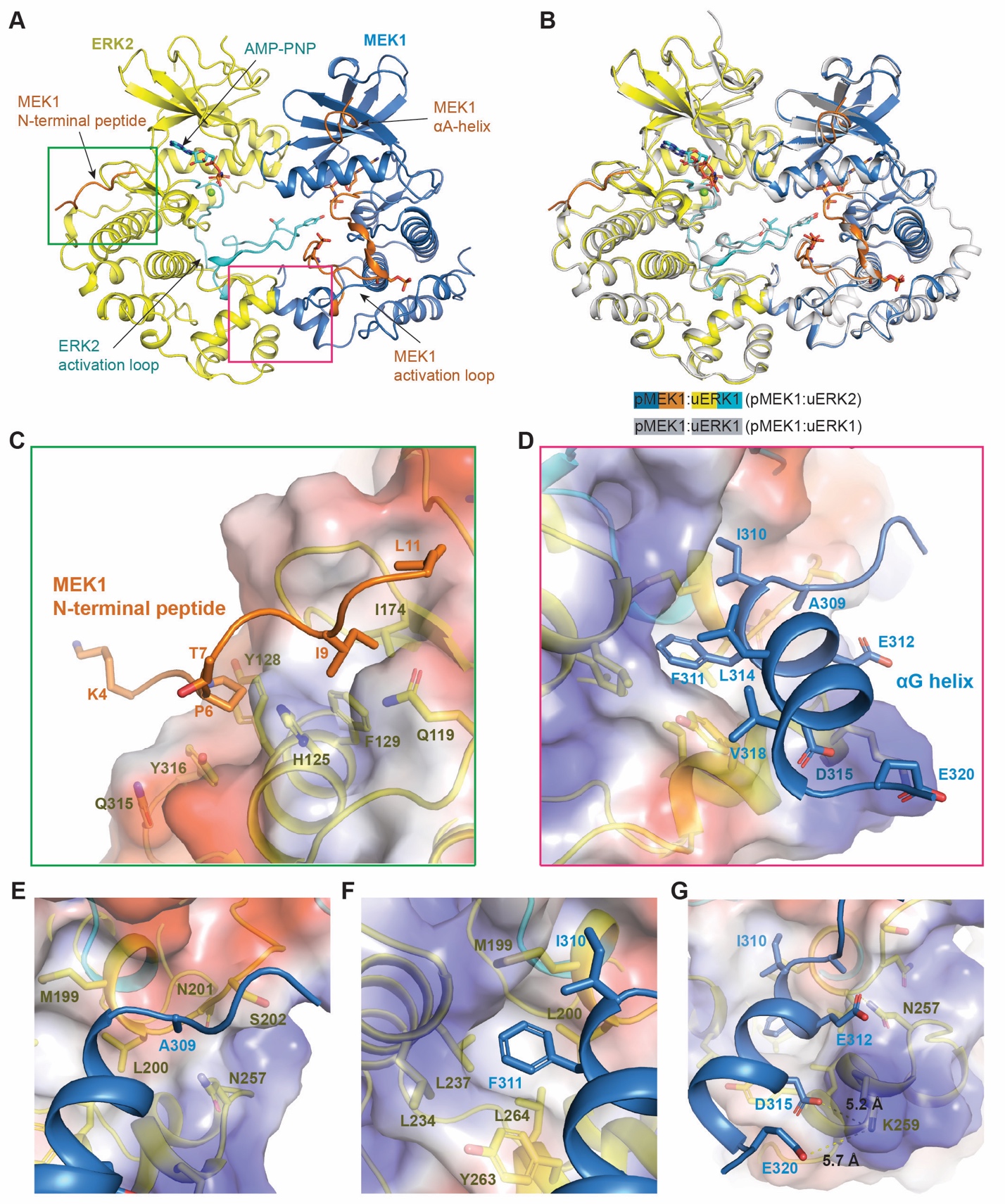

Figure S7. Cryo-EM analysis of the AMP-PNP-bound human pMEK1:uERK2 complex

(A) Overall structure of the pMEK1:uERK2 complex. Both pMEK1 and uERK2 in the complex bind to an AMP-PNP in their nucleotide binding pockets.

(B) Alignment of the structure of the pMEK1:uERK2 complex with that of the pMEK1:uERK1 complex.

(C) Interactions between the N-terminal peptide of MEK1 (colored green) and the D-site recruitment site (DRS) in ERK2.

(D) Binding of the A309-E320 helix in the MEK1 C-lobe (colored blue) to the F-site recruitment site (FRS) in ERK1.

(E–G) Close-up view of the binding of MEK1 A309 (E), I310 and F311 (F), and E312, D315 and E320 (G) to the FRS in ERK2. Hydrogen bonds are shown as yellow dash lines. The protein contact potential of ERK2 was generated in PyMOL.

All structural figures were made using PyMOL.

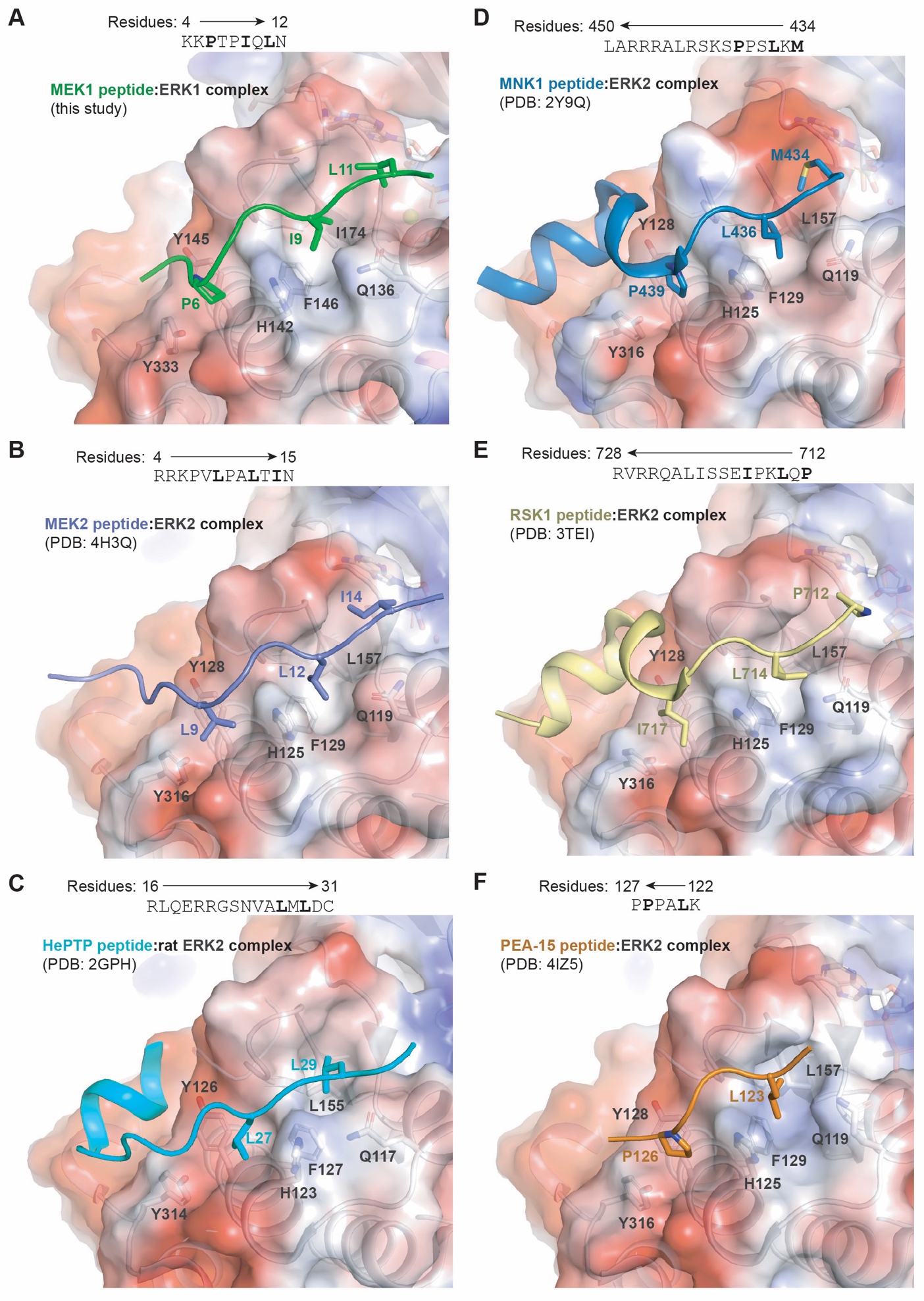

Figure S8. Comparison of the binding mode between the MEK1 N-terminal peptide and ERK1 DRS with that between reported DEJL motifs and ERK2 DRS

(A) Interaction between the N-terminal peptide (residues 4-12, colored green) of human MEK1 and human ERK1 DRS (cryo-EM structure of the pMEK1:uERK1 complex, this study).

(B) Interaction between the N-terminal peptide (residues 4-15, colored slate) of human MEK2 and human ERK2 DRS (PDB: 4H3Q).

(C) Interaction between a peptide (RLQERRGSNVALMLDV, colored cyan) of HePTP (hematopoietic tyrosine phosphatase, also named tyrosine-protein phosphatase non-receptor type 7) and rat ERK2 DRS (PDB: 2GPH).

(D) Interaction between a peptide (residues 434-450, colored blue) of human MNK1 (MAP kinase-interacting serine/threonine-protein kinase 1) and human ERK2 DRS (PDB: 2Y9Q).

(E) Interaction between a peptide (residues 712-728, colored yellow) of human RSK1 (ribosomal protein S6 kinase alpha-1) and human ERK2 DRS (PDB: 3TEI).

(F) Interaction between a peptide (residues 122-127, colored orange) of human PEA-15 (15 kDa phosphoprotein enriched in astrocytes) and ERK2 DRS (PDB: 4IZ5). The protein contact potential of ERK1 and ERK2 was generated in PyMOL.

All structural figures were made using PyMOL.

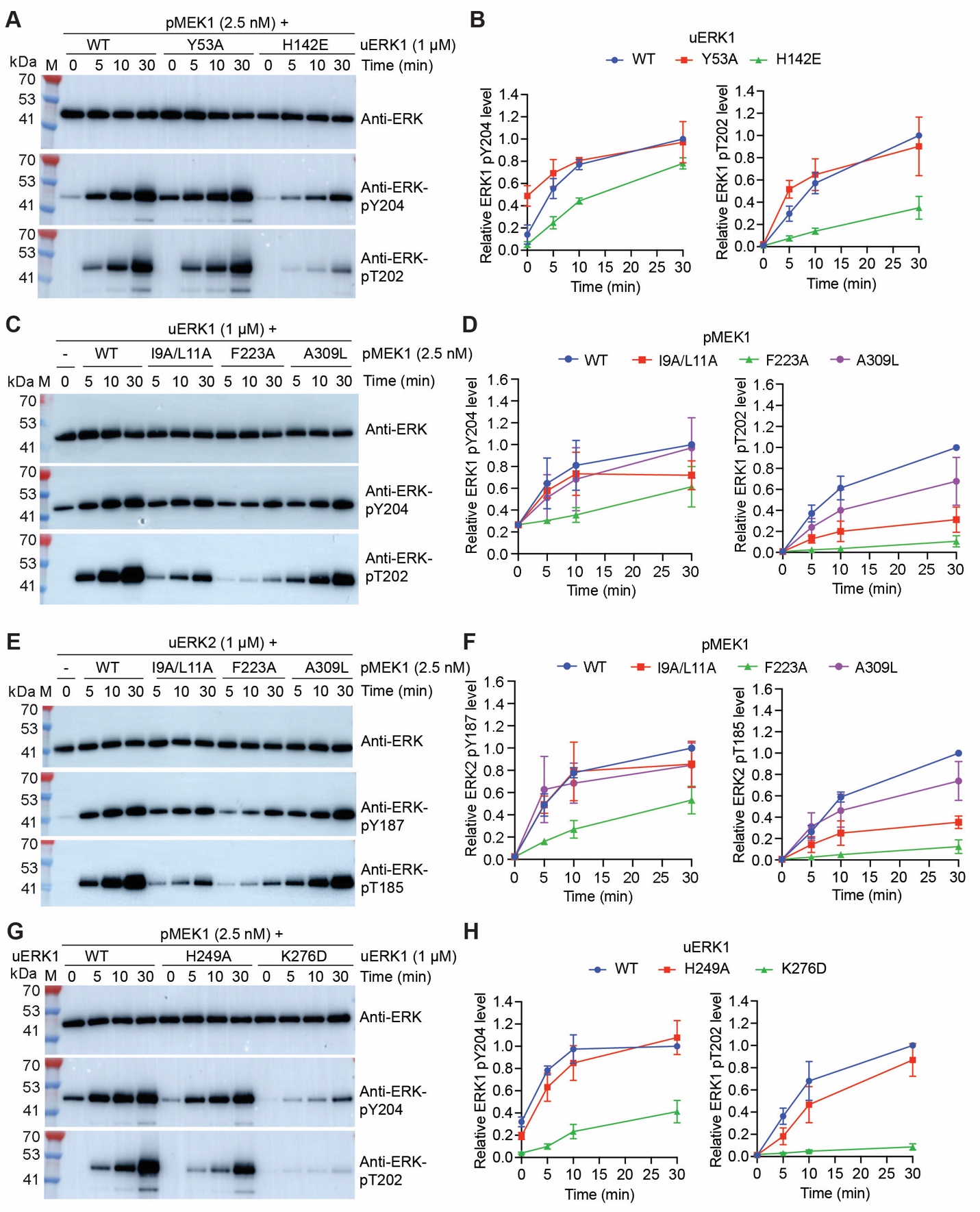

Figure S9. Identification of mutations in ERK1/2 and MEK1 that affect pMEK1-catalyzed ERK1 phosphorylation

In a buffer containing 5 mM ATP, 1 µM uERK1 (or uERK2) was incubated with 2.5 nM pMEK1 at 30 ℃, and then the phosphorylation of ERK1 T202 and Y204 (or ERK2 T185 and Y187) was monitored using Western blotting (A, C, E, G) and quantified (B, D, F, H).

(A and B) Effect of Y53A and H142E mutations in ERK1 on pMEK1-catalyzed phosphorylation of ERK1.

(C and D) Effect of I9A/L11A double mutation, F223A and A309L mutation in MEK1 on pMEK1-catalyzed phosphorylation of ERK1.

(E and F) Effect of I9A/L11A double mutation, F223A and A309L mutation in MEK1 on pMEK1-catalyzed phosphorylation of ERK2.

(G and H) Effect of H249A and K276D mutations in ERK1 on pMEK1-catalyzed phosphorylation of ERK1.

The gels in (A), (C), (E) and (G) are the results of a representative experiment out of three independent experiments. The data in (B), (D), (F) and (H) represent the mean ± SD of three independent measurements.

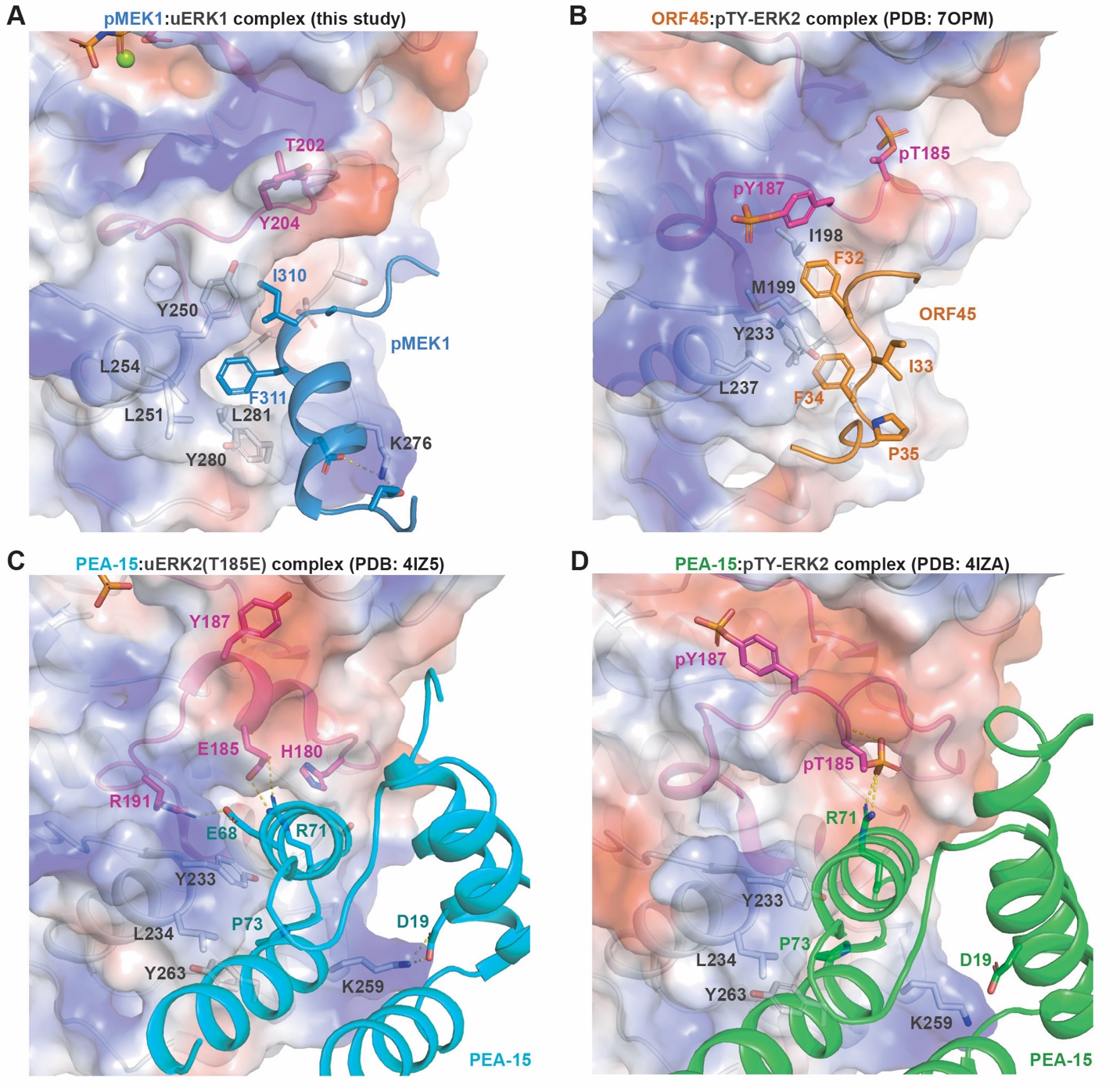

Figure S10. Comparison of the binding mode between ERK1 FRS and pMEK1 with that between ERK2 FRS and two of its binders

(A) Interaction between the A309-E320 helix (colored blue) in the human MEK1 C-lobe and the FRS in human ERK1 (cryo-EM structure of the pMEK1:uERK1 complex, this study).

(B) Interaction between Kaposi's sarcoma associated herpesvirus protein ORF45 (colored orange) and the FRS in human ERK2 (PDB: 7OPM).

(C) Interaction between human PEA-15 (colored cyan) and the FRS in the T185E mutant of unphosphorylated human ERK2 (PDB: 4IZ5).

(D) Interaction between human PEA-15 (colored green) and the FRS in the T185/Y187 dual-phosphorylated human ERK2 (PDB: 4IZA).

The protein contact potential of ERK1 and ERK2 was generated in PyMOL. All structural figures were made using PyMOL.

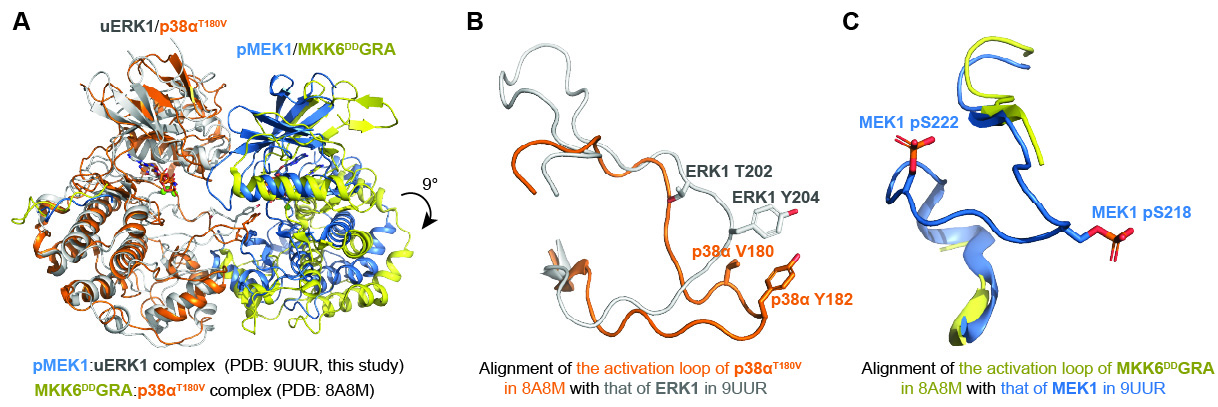

Figure S11. Comparison of the cryo-EM structure of the pMEK1:uERK1 complex (PDB: 9UUR, this study) with that of the MKK6^DD^GRA:p38α^T180V^ complex (PDB: 8A8M, Juyoux *et al*.^1^)

(A) Comparison of the overall structure of the pMEK1:uERK1 complex with that of the MKK6^DD^GRA:p38α^T180V^ complex. When the C-lobe of p38α is aligned with that of ERK1, the kinase domain of MKK6 is rotated by 9° relative to that of MEK1.

(B) The activation loop of ERK1 in the pMEK1:uERK1 complex and that of p38α in the MKK6^DD^GRA:p38α^T180V^ complex adopt distinct conformations.

(C) The activation loop of MKK6 in the MKK6^DD^GRA:p38α^T180V^ complex is largely invisible, while the activation loop of MEK1 in the pMEK1:uERK1 complex is well-resolved.

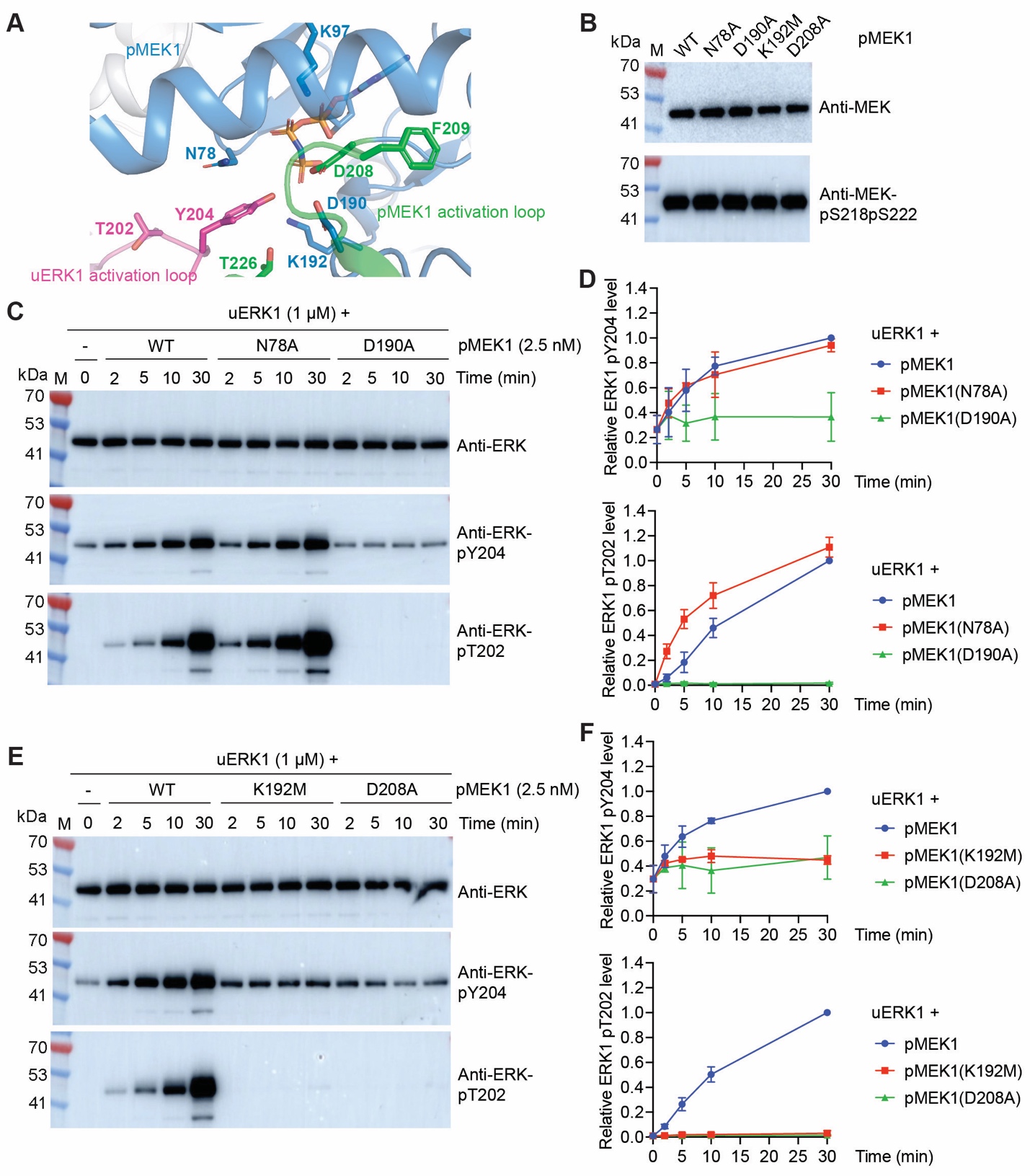

Figure S12. Effect of mutations of residues forming the catalytic pocket of MEK1 on the kinase activity of pMEK1

(A) Binding of uERK1 Y204 (colored magenta) to the catalytic pocket of pMEK1 in the cryo-EM structure of the pMEK1:uERK1 complex.

(B) The phosphorylation state of wild-type and mutant pMEK1 was analyzed by Western blotting. (C and D) Effect of N78A and D190A mutations in MEK1 on the kinase activity of pMEK1. In a buffer containing 5 mM ATP, 1 µM uERK1 was incubated with 2.5 nM pMEK1 at 30 ℃, and then the phosphorylation of ERK1 T202 and Y204 was monitored using Western blotting (C) and quantified (D).

(E and F) Effect of K192M and D208A mutations in MEK1 on the kinase activity of pMEK1.

The gels in (C) and (E) are the results of a representative experiment out of three independent experiments. The data in (D) and (F) represent the mean ± SD of three independent measurements.

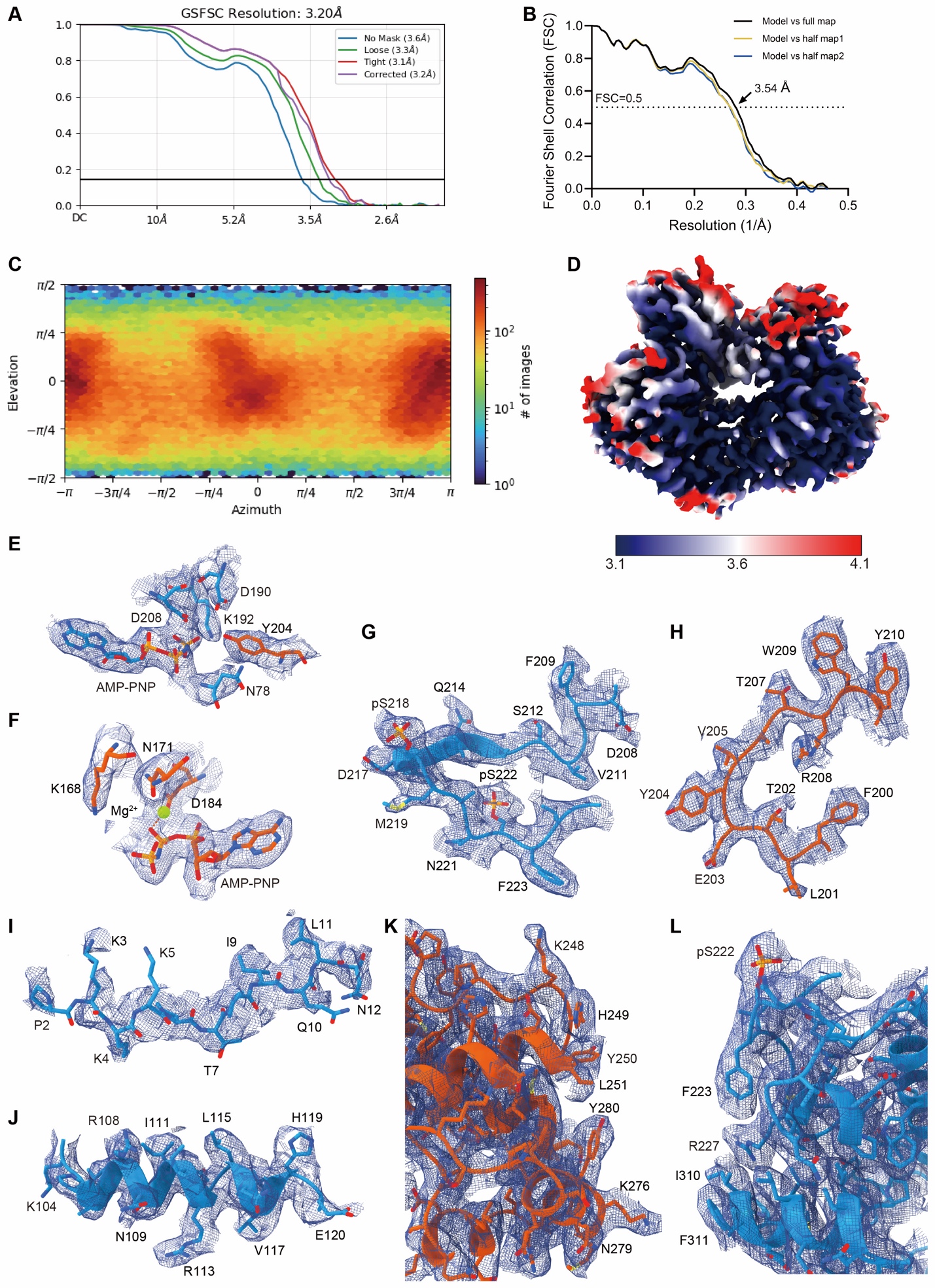

Figure S13. Cryo-EM analysis of the AMP-PNP-bound pMEK1:uERK1 complex (derived from AMP-PNP-bound pMEK1:pY-ERK1 complex)

(A) FSC curve for the final reconstruction of the pMEK1:uERK1 complex generated by CryoSPARC. Four curves correspond to the measurements with no mask (3.6 Å), loose mask (3.3 Å), tight mask (3.1 Å) and corrected mask (3.2 Å).

(B) FSC curves of the final refined model versus the summed map that it was refined against (black); of the model versus the first half-map (yellow); and of the model versus the second half-map (blue). The small difference between the yellow and blue curves indicates that the refinement of the atomic coordinates did not suffer from overfitting.

(C) Euler-angle distributions of the particles used for the final reconstruction.

(D) Local resolution of the reconstructed map.

(E–L) Representative cryo-EM maps of MEK1 (blue) and ERK1 (orange), at a contour level of 0.48. (E, F) ATP binding pockets. (G, H) activation loops. (I) N-terminal ERK docking site on MEK1. (J) αC helix of MEK1. (K, L) C-lobes near interface.

The cryo-EM maps of human pMEK1:uERK1 are shown as blue meshes. The structural figures were made using UCSF ChimeraX (1.9).

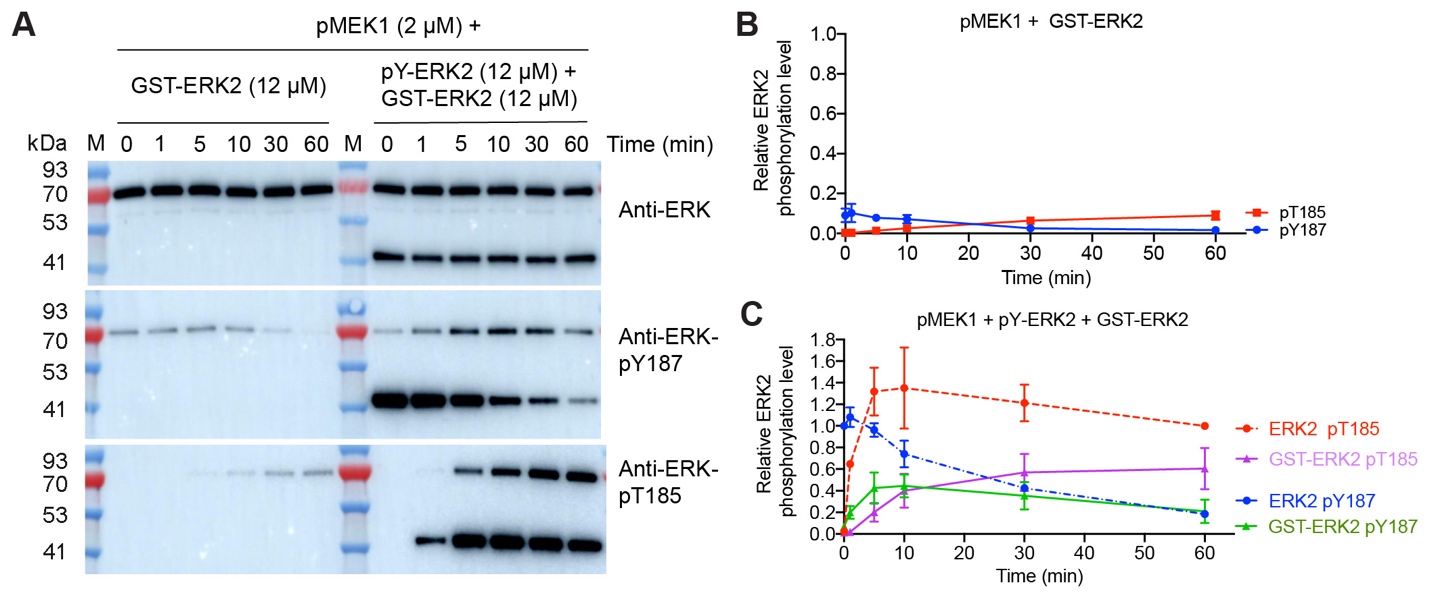

Figure S14. Dephosphorylation of ERK2 and phosphate transfer from ERK2 Y187 to T185 catalyzed by pMEK1

(A–C) pMEK1-catalyzed intramolecular and intermolecular phosphate group transfer from ERK2 Y187 to T185. In a nucleotide-free buffer, GST-ERK2 alone (slightly phosphorylated at ERK2 Y187) or together with pY-ERK2 was incubated with pMEK1 at 30 ℃. The phosphorylation of ERK1 T185 and Y187 was evaluated using Western blotting (A) and quantified (B and C).

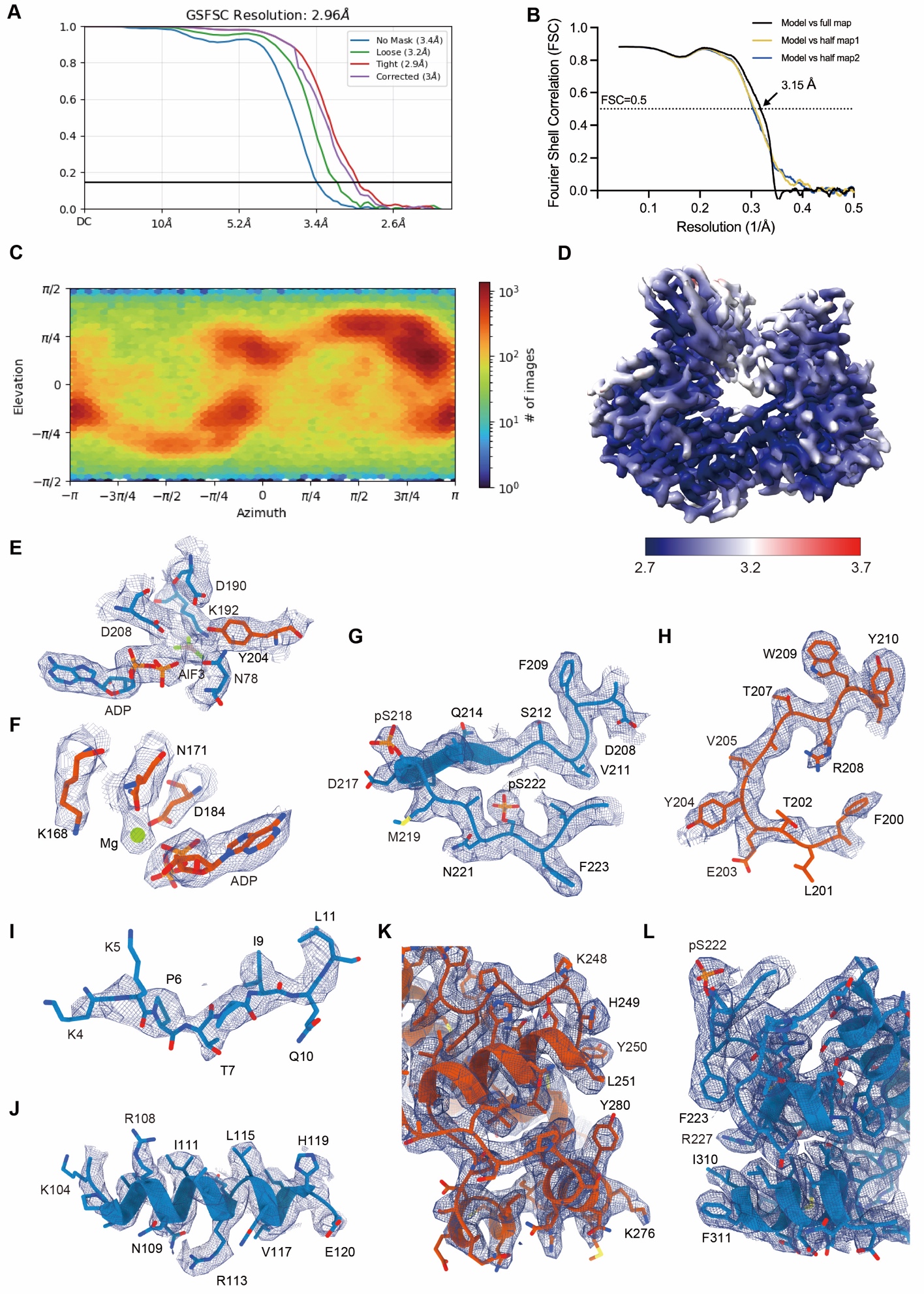

Figure S15 | Cryo-EM analysis of the ADP and AlF_3_-bound pMEK1:uERK1 complex.

(A) FSC curve for the final reconstruction of the pMEK1:uERK1 complex generated by CryoSPARC. Four curves correspond to the measurements with no mask (3.4 Å), loose mask (3.2 Å), tight mask (2.9 Å) and corrected mask (3.0 Å).

(B) FSC curves of the final refined model versus the summed map that it was refined against (black); of the model versus the first half-map (yellow); and of the model versus the second half-map (blue). The small difference between the yellow and blue curves indicates that the refinement of the atomic coordinates did not suffer from overfitting. (C) Euler-angle distributions of the particles used for the final reconstruction.

(D) Local resolution of the reconstructed map.

(E–L) Representative cryo-EM maps of MEK1 (blue) and ERK1 (orange), at a contour level of 0.43. (E, F) ATP binding pockets. (G, H) activation loops. (I) N-terminal ERK docking site on MEK1. (J) αC helix of MEK1. (K, L) C-lobes near interface.

The cryo-EM maps of human pMEK1:pY204ERK1 are shown as blue meshes. All structural figures were made using UCSF ChimeraX (1.9).

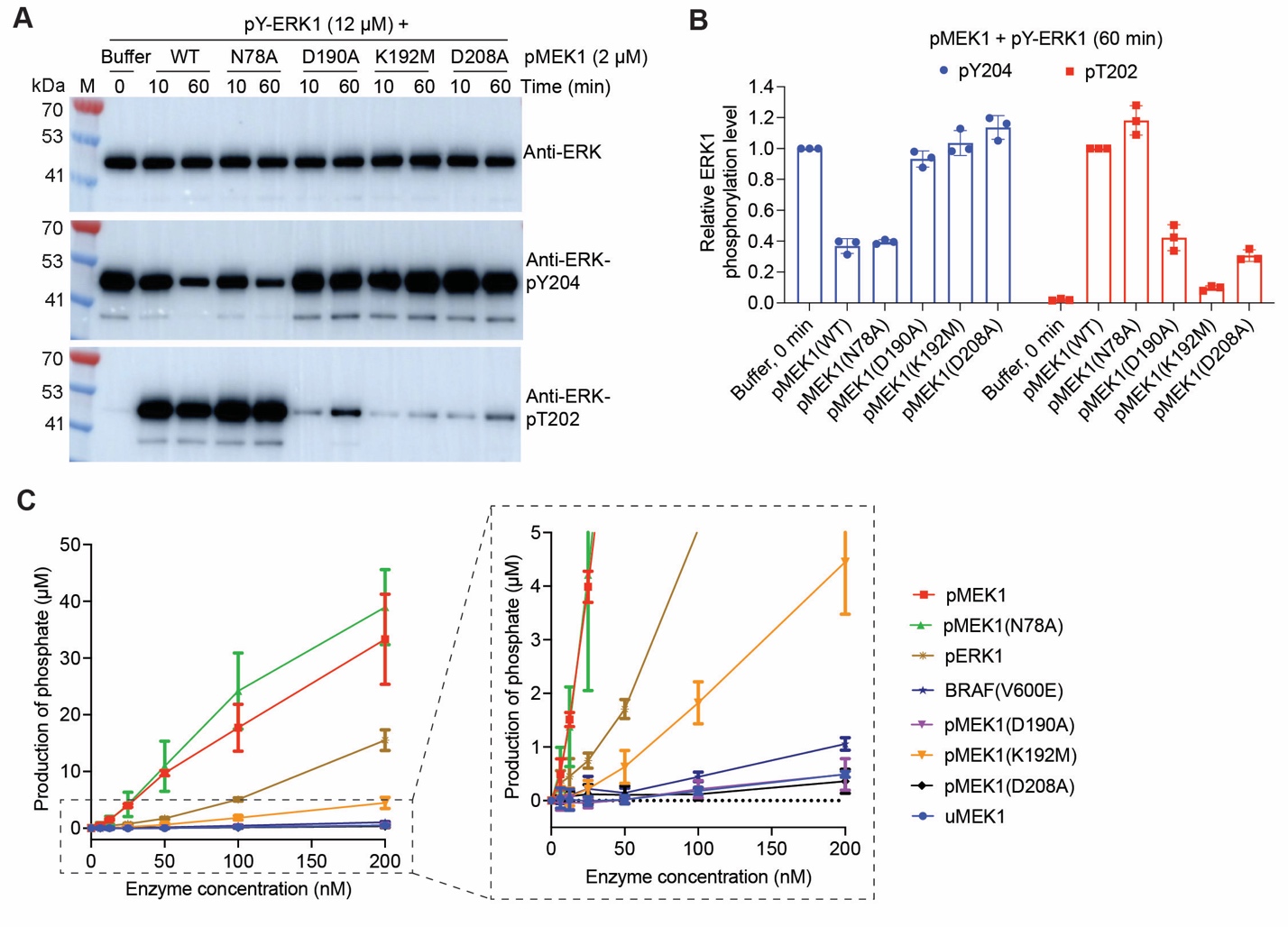

Figure S16. Effect of mutations of residues forming the catalytic pocket of MEK1 on the phosphate transfer and ATPase activity of pMEK1

(A and B) Effect of mutations of the nucleotide-binding pocket of MEK1 on the activity of pMEK1 to catalyze phosphate group transfer from ERK1 Y204 to T202. In a nucleotide-free buffer, pY-ERK1 was incubated with wild-type (WT) or mutant pMEK1 at 30 ℃, and then the phosphorylation of ERK1 T202 and Y204 was evaluated using Western blotting (A) and quantified (B).

(C) Comparison of the ATPase activity of wild-type pMEK1 with that of uMEK1, pMEK1 mutants, pERK1, and BRAF(V600E).

The gels in (A) are the results of a representative experiment out of three independent experiments. The data in (B) and (C) represent the mean ± SD of three independent measurements.

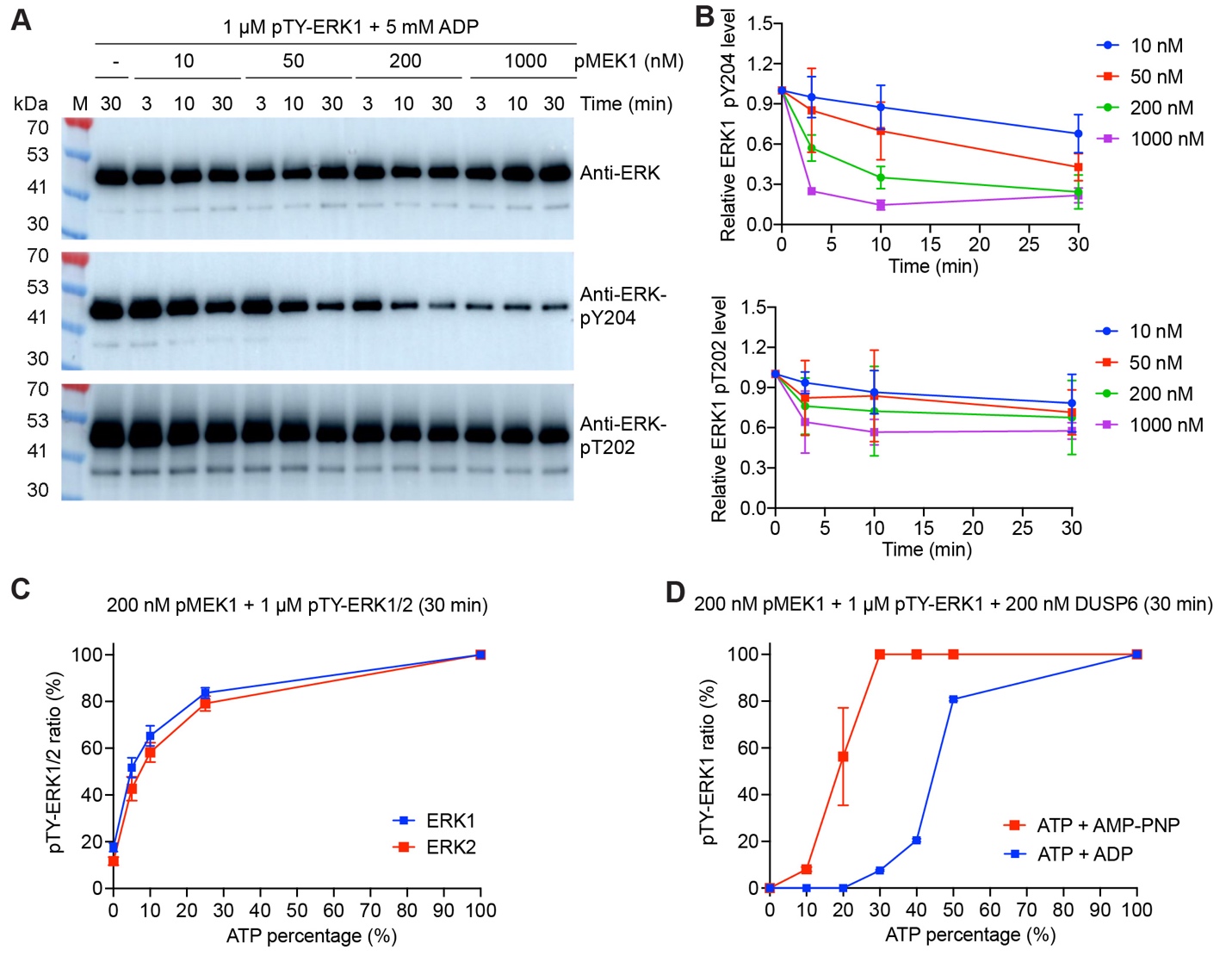

Figure S17. pMEK1-catalyzed dephosphorylation of pTY-ERK1/2 under different nucleotide conditions

(A and B) pMEK1-catalyzed dephosphorylation of pTY-ERK1. pTY-ERK1 was incubated with increasing concentrations of pMEK1 in the presence of 5 mM ADP at 30 °C. At the indicated time points, the phosphorylation of ERK1 T202 and Y204 was evaluated using Western blotting (A) and quantified (B).

(C) Dephosphorylation of pTY-ERK1 and pTY-ERK2 catalyzed by pMEK1 in ATP/ADP mixtures. The total concentration of ATP and ADP was 5 mM in each sample, with varying ATP percentages.

(D) Effect of DUSP6 on pMEK1-catalyzed dephosphorylation of pTY-ERK1 in ATP and ADP or ATP and AMP-PNP mixtures. The total nucleotide concentration was 5 mM in each sample, and the percentage of ATP was varied in different samples.

The reactions in (C) and (D) were carried out at 30 °C for 30 min, then ERK1 phosphorylation levels were determined by LC–MS. The gels in (A) are the results of a representative experiment out of three independent experiments. The data in (B), (C) and (D) represent the mean ± SD of three biological replicates.

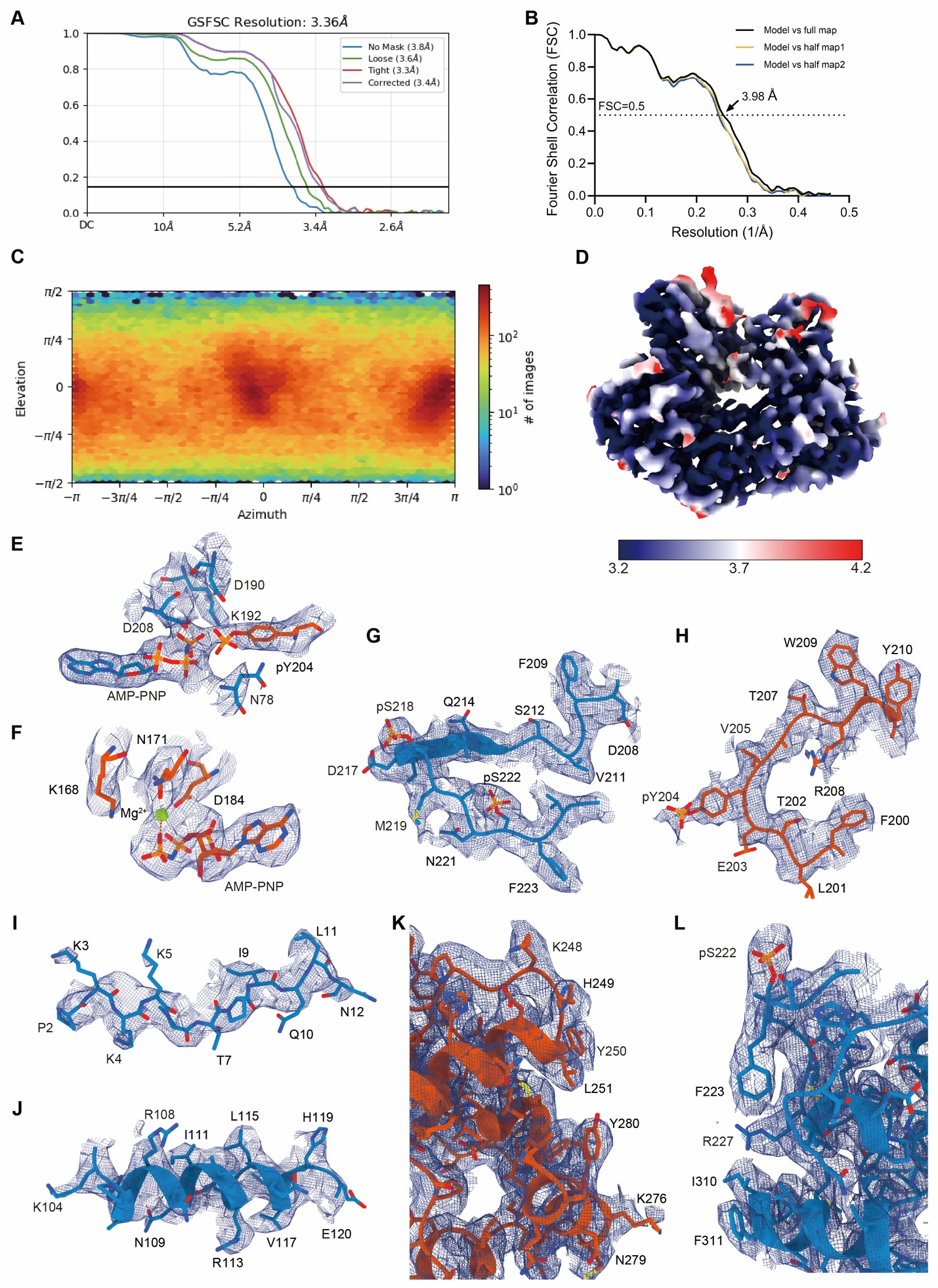

Figure S18. Cryo-EM analysis of the AMP-PNP-bound pMEK1:pY-ERK1 complex

(A) FSC curve for the final reconstruction of the pMEK1:pY-ERK1 complex generated by CryoSPARC. Four curves correspond to the measurements with no mask (3.8 Å), loose mask (3.6 Å), tight mask (3.3 Å) and corrected mask (3.4 Å). (B) FSC curves of the final refined model versus the summed map that it was refined against (black); of the model versus the first half-map (yellow); and of the model versus the second half-map (blue). The small difference between the yellow and blue curves indicates that the refinement of the atomic coordinates did not suffer from overfitting.

(C) Euler-angle distributions of the particles used for the final reconstruction. (D) Local resolution of the reconstructed map.

(E–L) Representative cryo-EM maps of MEK1 (blue) and ERK1 (orange), at a contour level of 0.55. (E, F) ATP binding pockets. (G, H) activation loops. (I) N-terminal ERK docking site on MEK1. (J) αC helix of MEK1. (K, L) C-lobes near interface.

The cryo-EM maps of human pMEK1:pY-ERK1 are shown as blue meshes. All structural figures were made using UCSF ChimeraX (1.9).

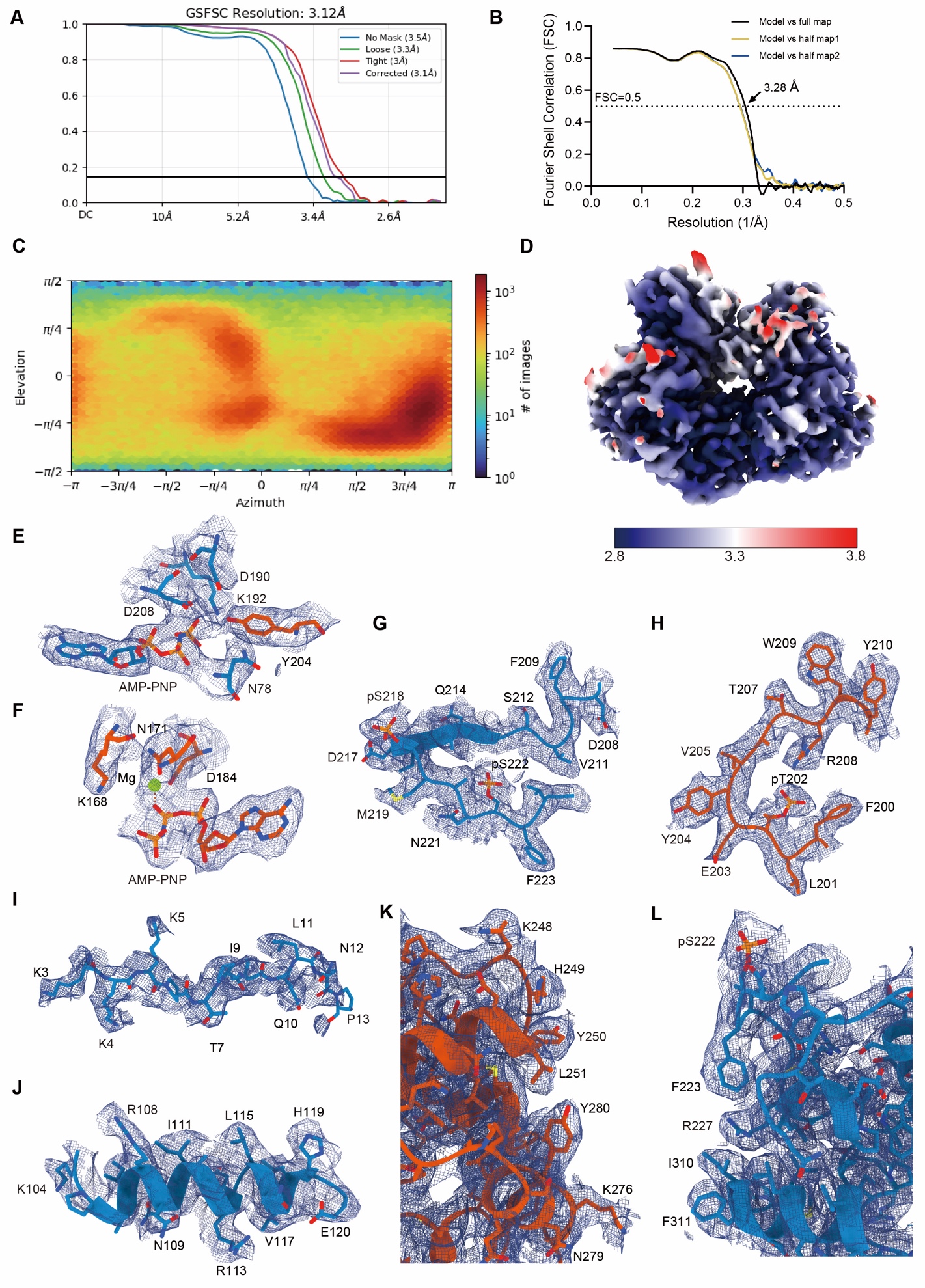

Figure S19. Cryo-EM analysis of the AMP-PNP-bound pMEK1:pT-ERK1 complex (derived from AMP-PNP-bound pMEK1:pTY-ERK1 complex)

(A) FSC curve for the final reconstruction of the pMEK1:pT-ERK1 complex generated by CryoSPARC. Four curves correspond to the measurements with no mask (3.5 Å), loose mask (3.3 Å), tight mask (3.0 Å) and corrected mask (3.1 Å).

(B) FSC curves of the final refined model versus the summed map that it was refined against (black); of the model versus the first half-map (yellow); and of the model versus the second half-map (blue). The small difference between the yellow and blue curves indicates that the refinement of the atomic coordinates did not suffer from overfitting.

(C) Euler-angle distributions of the particles used for the final reconstruction.

(D) Local resolution of the reconstructed map.

(E–L) Representative cryo-EM maps of MEK1 (blue) and ERK1 (orange), at a contour level of 0.52. (E, F) ATP binding pockets. (G, H) activation loops. (I) N-terminal ERK docking site on MEK1. (J) αC helix of MEK1. (K, L) C-lobes near interface.

The cryo-EM maps of human pMEK1:pT-ERK1 are shown as blue meshes. All structural figures were made using UCSF ChimeraX (1.9).

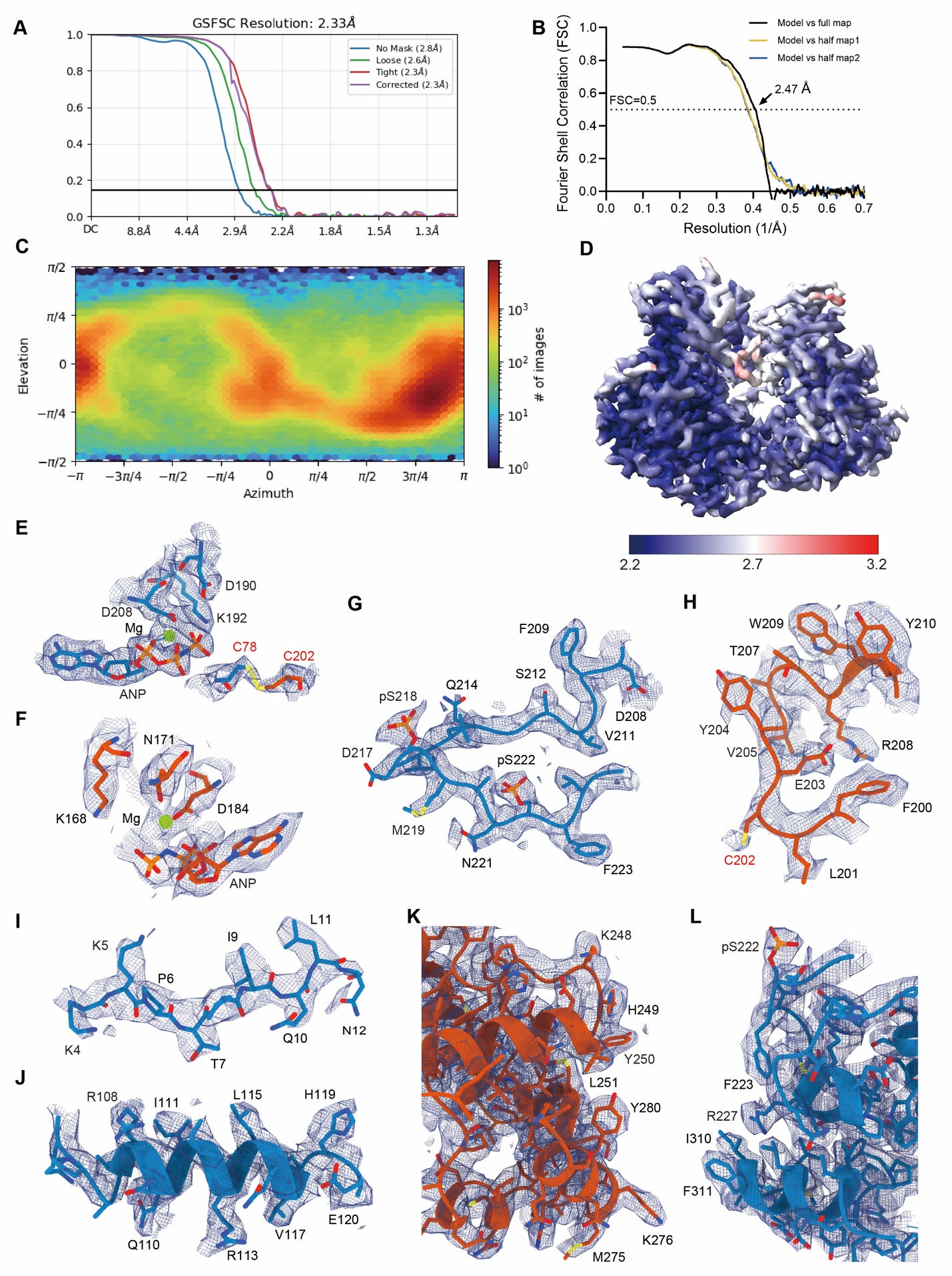

Figure S20. Cryo-EM analysis of the AMP-PNP-bound crosslinked pMEK1(N78C):uERK1(T202C) complex

(A) FSC curve for the final reconstruction of the crosslinked pMEK1(N78C):uERK1(T202C) complex generated by CryoSPARC. Four curves correspond to the measurements with no mask (2.8 Å), loose mask (2.6 Å), tight mask (2.3 Å) and corrected mask (2.3 Å).

(B) FSC curves of the final refined model versus the summed map that it was refined against (black); of the model versus the first half-map (yellow); and of the model versus the second half-map (blue). The small difference between the yellow and blue curves indicates that the refinement of the atomic coordinates did not suffer from overfitting.

(C) Euler-angle distributions of the particles used for the final reconstruction.

(D) Local resolution of the reconstructed map.

(E–L) Representative cryo-EM maps of MEK1 (blue) and ERK1 (orange), at a contour level of 0.08. (E, F) ATP binding pockets. (G, H) activation loops. (I) N-terminal ERK docking site on MEK1. (J) αC helix of MEK1. (K, L) C-lobes near interface.

The cryo-EM maps of human the pMEK1(N78C):uERK1(T202C) complex are shown as blue meshes, point mutations are indicated in red labels. All structural figures were made using UCSF ChimeraX (1.9).

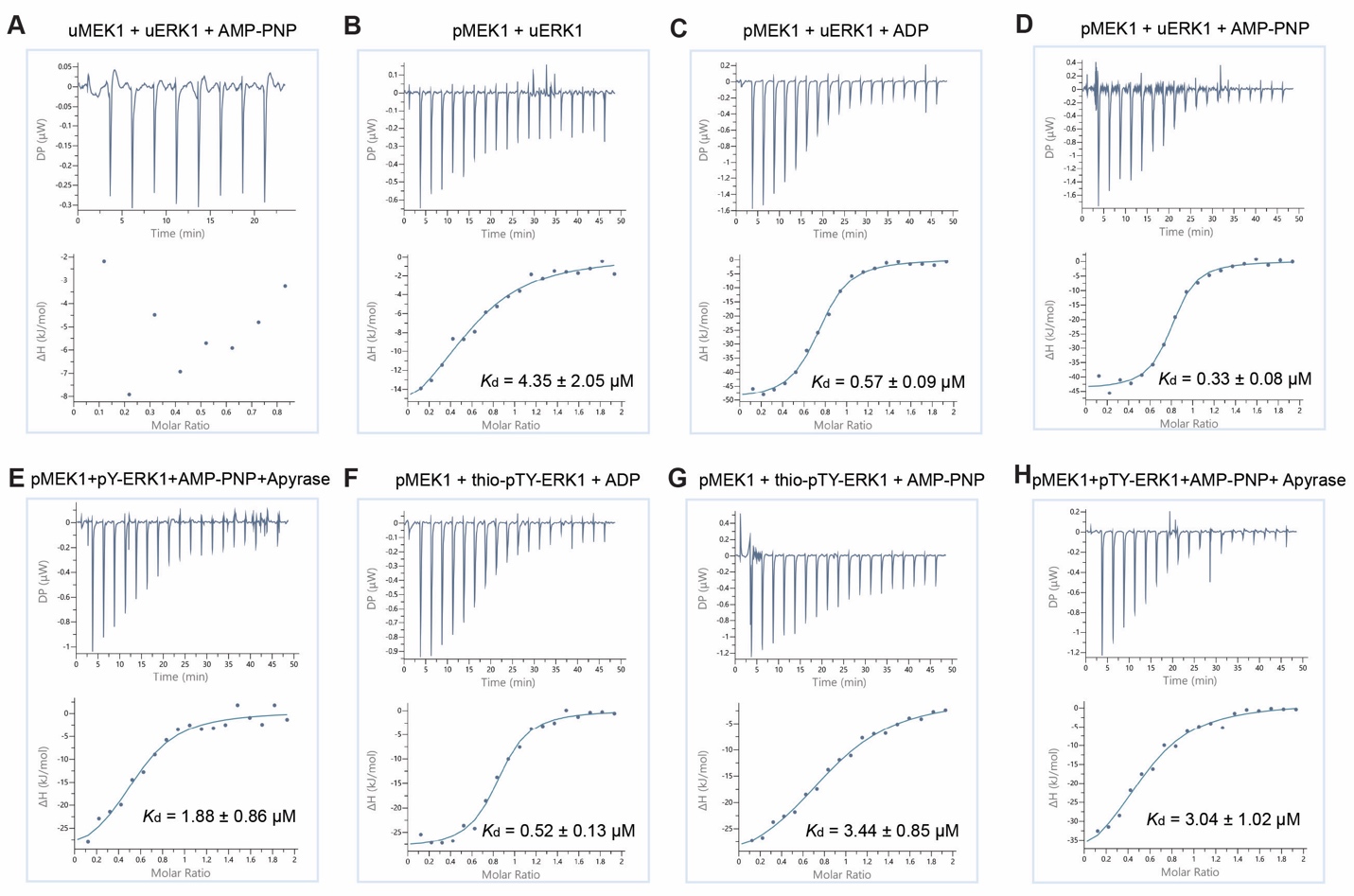

Figure S21. Binding affinity between uMEK1 or pMEK1 and uERK1, pY-ERK1, thio-pTY-ERK1, or pTY-ERK1 in the absence or presence of ADP or AMP-PNP

(A–H The dissociation constants for the binding between uMEK1 and uERK1 in the presence of AMP-PNP (A), between pMEK1 and uERK1 in the absence of nucleotides (B) or in the presence of ADP (C) or AMP-PNP (D), between apyrase treated pMEK1 and pY-ERK1 in the presence of AMP-PNP (E), between pMEK1 and thio-pTY-ERK1 in the presence of ADP (F) or AMP-PNP (G), and between apyrase treated pMEK1 and pTY-ERK1 in the presence of AMP-PNP (H) were measured using Isothermal Titration Calorimetry (ITC).

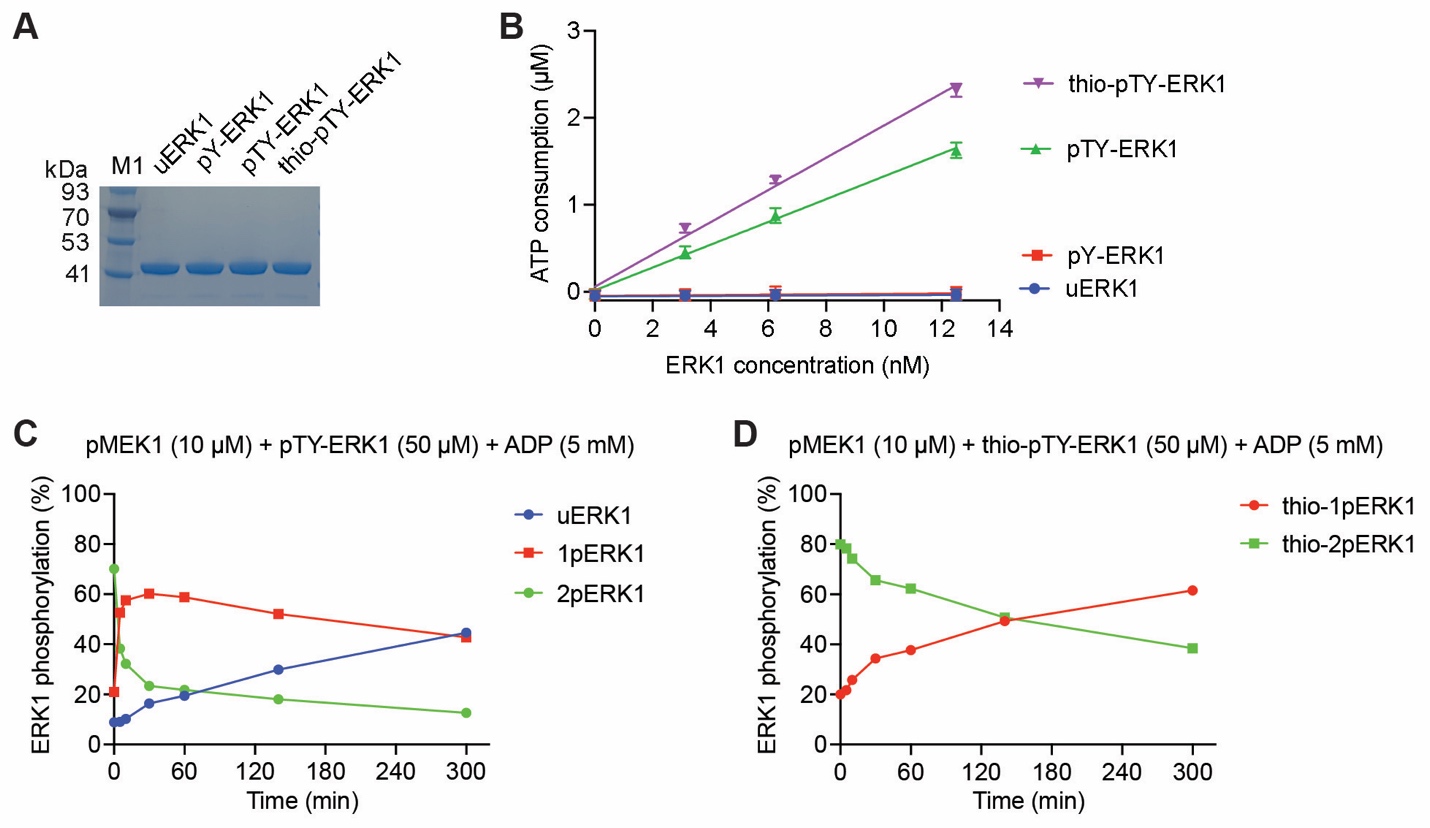
 Figure S22. Characterization of thio-pTY-ERK1

(A) Purified unphosphorylated (uERK1), phosphorylated (pY-ERK1 and pTY-ERK1), and thio-phosphorylated ERK1 (thio-pTY-ERK1) were examined by SDS–PAGE and visualized by Coomassie blue staining.

(B) Comparison of the ATP consumption rate of thio-pTY-ERK1 with those of uERK1, pY-ERK1, and pTY-ERK1. The ATP consumption was measured using an ADP-Glo Kinase Assay. The MBP peptide FFKNIVTPRTPPPSQGK was used as the substrate of ERK1. The data represent the mean ± SD of three independent measurements.

(C and D) pMEK1-catalyzed dephosphorylation of pTY-ERK1 (C) and thio-pTY-ERK1 (D) in an ADP-containing buffer. The percentage of ERK1 in different phosphorylation states was determined by LC-MS.

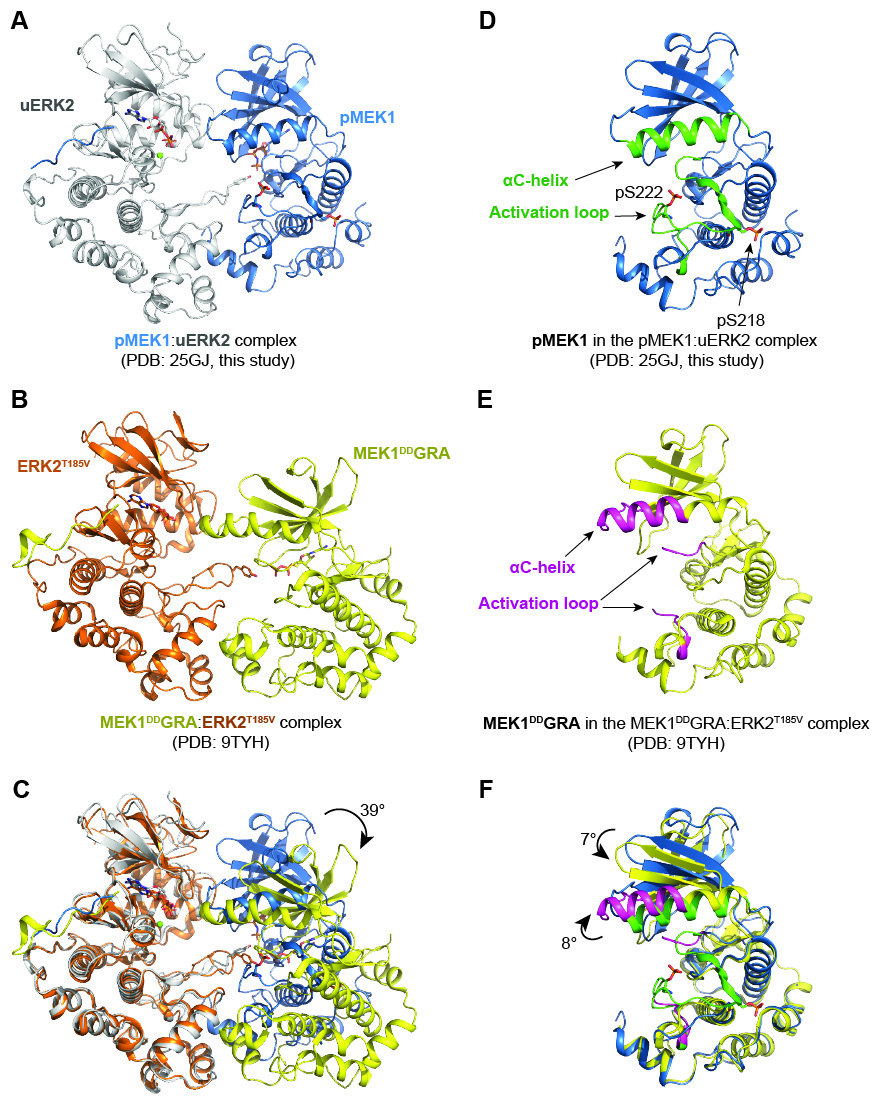
 Figure S23. Comparison of the cryo-EM structure of the pMEK:uERK2 complex (PDB: 25GJ, this study) with that of the MEK1^DD^GRA:ERK2^T185V^ complex (PDB: 9TYH, von Velsen *et al*.^2^)

(A) Cryo-EM structure of the pMEK1:uERK2 complex.

(B) Cryo-EM structure of the MEK1^DD^GRA:ERK2^T185V^ complex. In this complex, residues 3–11 of MEK1 were replaced with the KIM sequence from the *Toxoplasma gondii* effector protein GRA24, S218 and S222 in MEK1 were mutated to aspartate, and T185 in the activation loop of ERK2 was mutated to a valine to stabilize the complex.

(C) Comparison of the cryo-EM structure of the pMEK1:uERK2 complex with that of the MEK1^DD^GRA:ERK2^T185V^ complex. When the C-lobes of ERK2 in the two structures are aligned, the kinase domain of MEK1 in the MEK1^DD^GRA:ERK2^T185V^ complex is rotated by 39° relative to that in the pMEK1:uERK2 complex.

(D) Structure of MEK1 in the pMEK1:uERK2 complex.

(E) Structure of MEK1 in the MEK1^DD^GRA:ERK2^T185V^ complex. The activation loop of MEK1 is largely invisible in this structure.

(F) Comparison of the structure of MEK1 in the pMEK1:uERK2 complex with that in the MEK1^DD^GRA:ERK2^T185V^ complex. When the C-lobes of MEK1 in the two structures are aligned, the αC-helix of MEK1 in the MEK1^DD^GRA:ERK2^T185V^ complex is rotated by 8° relative to that in the pMEK1:uERK2 complex; the β-sheet in the N-lobe of MEK1 is rotated to by 7° to the opposite direction; and the activation loops of MEK1 in the two structures adopt distinct conformations.

**Table S1 | Data collection and refinement statistics (continued on next page).**

|  | pMEK1:uERK1 (EMD-64516, PDB: 9UUR) | pMEK1:uERK1 (from pMEK1:pY-ERK1)  (EMD-64545, PDB: 9UW4) | pMEK1:pT-ERK1 (from pMEK1:pTY-ERK1)  (EMD-64519, PDB: 9UUX) | pMEK1:pY-ERK1  (EMD-64544, PDB: 9UW3) |
| --- | --- | --- | --- | --- |
| **Data collection and processing** |  |  |  |  |
| Microscope | FEI Titan Krios | FEI Titan Krios | FEI Titan Krios | FEI Titan Krios |
| Magnification | 81,000 | 81,000 | 81,000 | 81,000 |
| Voltage (kV) | 300 | 300 | 300 | 300 |
| Detector | Gatan K3 | Gatan K3 | Gatan K3 | Gatan K3 |
| Electron exposure (e^–^ Å^-2^) | 50 | 50 | 50 | 50 |
| Defocus range (μm) | ~-1.1 to -1.8 | ~-1.1 to -1.8 | ~-1.1 to -1.8 | ~-1.1 to -1.8 |
| Pixel size (Å) | 1.0773 | 1.087 | 1.0773 | 1.0773 |
| Symmetry imposed | C1 | C1 | C1 | C1 |
| Initial particle images (no.) | 24,513,458 | 25,107,479 | 4,226,273 | 11,179,582 |
| Final particle images (no.) | 319,939 | 235,273 | 555,122 | 216,414 |
| Map resolution (Å) | 3.36 | 3.20 | 3.12 | 3.36 |
| FSC threshold | 0.143 | 0.143 | 0.143 | 0.143 |
| Map resolution range (Å) | 2.8-8.8 | 2.7-8.6 | 2.7-7.9 | 2.9-5.9 |
| **Refinement** |  |  |  |  |
| Initial model used | AlphaFold3 | AlphaFold3 | AlphaFold3 | AlphaFold3 |
| Model resolution (Å) | 3.76 | 3.54 | 3.28 | 3.98 |
| FSC threshold | 0.5 | 0.5 | 0.5 | 0.5 |
| Model composition |  |  |  |  |
| Nonhydrogen atoms | 5300 | 5269 | 5310 | 5204 |
| Protein residues | 657 | 652 | 657 | 643 |
| Ligand | 2 | 2 | 2 | 2 |
| Water | 0 | 0 | 0 | 0 |
| Ion | 1 | 1 | 1 | 1 |
| Root mean square deviations |  |  |  |  |
| Bond lengths (Å) | 0.008 | 0.008 | 0.008 | 0.008 |
| Bond angles (°) | 1.212 | 1.186 | 1.172 | 1.212 |
| Validation | | |  |  |
| MolProbity score | 1.91 | 1.76 | 1.95 | 1.88 |
| Clashscore | 8.50 | 5.60 | 8.39 | 8.07 |
| Poor rotamers (%) | 0.00 | 0.00 | 0.35 | 0.18 |
| Ramachandran plot |  |  |  |  |
| Favored (%) | 92.98 | 92.77 | 91.69 | 93.27 |
| Allowed (%) | 7.02 | 7.23 | 8.31 | 6.73 |
| Disallowed (%) | 0.00 | 0.00 | 0.00 | 0.00 |

**Table S1 | Data collection and refinement statistics (continued).**

|  | pMEK1:uERK2  (EMD-80086, PDB: 25GJ) | pMEK1:uERK1  (ADP+AlF_3_)  (EMD-80120, PDB: 25HU) | pMEK1 N78C:uERK1 T202C  (EMD-82382, PDB:43YB) |
| --- | --- | --- | --- |
| **Data collection and processing** |  |  |  |
| Microscope | FEI Titan Krios | FEI Titan Krios | FEI Titan Krios |
| Magnification | 81,000 | 81,000 | 215,000 |
| Voltage (kV) | 300 | 300 | 300 |
| Detector | Gatan K3 | Gatan K3 | Falcon 4i |
| Electron exposure (e^–^ Å^-2^) | 50 | 50 | 50 |
| Defocus range (μm) | ~-1.1 to -1.8 | ~-1.1 to -1.8 | ~-1.0 to -1.9 |
| Pixel size (Å) | 1.0773 | 1.0773 | 0.57 |
| Symmetry imposed | C1 | C1 | C1 |
| Initial particle images (no.) | 12,045,667 | 11,784,916 | 52,692,322 |
| Final particle images (no.) | 609,769 | 403,726 | 1,082,206 |
| Map resolution (Å) | 2.93 | 2.96 | 2.33 |
| FSC threshold | 0.143 | 0.143 | 0.143 |
| Map resolution range (Å) | 2.6-3.8 | 2.5-5.5 | 2.1-3.2 |
| **Refinement** |  |  |  |
| Initial model used | AlphaFold3 | AlphaFold3 | AlphaFold3 |
| Model resolution (Å) | 3.01 | 3.15 | 2.47 |
| FSC threshold | 0.5 | 0.5 | 0.5 |
| Model composition |  |  |  |
| Nonhydrogen atoms | 5103 | 5033 | 5268 |
| Protein residues | 630 | 620 | 645 |
| Ligand | 2 | 3 | 2 |
| Water | 0 | 0 | 0 |
| Ion | 1 | 1 | 2 |
| Root mean square deviations |  |  |  |
| Bond lengths (Å) | 0.007 | 0.005 | 0.006 |
| Bond angles (°) | 1.218 | 1.051 | 0.701 |
| Validation | | |  |
| MolProbity score | 1.75 | 1.57 | 1.42 |
| Clashscore | 6.67 | 3.47 | 3.71 |
| Poor rotamers (%) | 0.54 | 0.00 | 0.00 |
| Ramachandran plot |  |  |  |
| Favored (%) | 94.46 | 93.40 | 96.20 |
| Allowed (%) | 5.21 | 6.60 | 3.80 |
| Disallowed (%) | 0.33 | 0.00 | 0.00 |
